# Long somatic DNA-repeat expansion drives neurodegeneration in Huntington disease

**DOI:** 10.1101/2024.05.17.592722

**Authors:** Robert E. Handsaker, Seva Kashin, Nora M. Reed, Steven Tan, Won-Seok Lee, Tara M. McDonald, Kiely Morris, Nolan Kamitaki, Christopher D. Mullally, Neda Morakabati, Melissa Goldman, Gabriel Lind, Rhea Kohli, Elisabeth Lawton, Marina Hogan, Kiku Ichihara, Sabina Berretta, Steven A. McCarroll

## Abstract

Huntington Disease (HD) is a fatal genetic disease in which most striatal projection neurons (SPNs) degenerate. The central biological question about HD pathogenesis has been how the disease-causing DNA repeat expansion (CAG_n_) in the *huntingtin* (*HTT*) gene leads to neurodegeneration after decades of apparent latency. Inherited *HTT* alleles with a longer CAG repeat hasten disease onset; the length of this repeat also changes over time, generating somatic mosaicism, and genes that regulate DNA-repeat stability can influence HD age-at-onset. To understand the relationship between a cell’s CAG-repeat length and its biological state, we developed a single-cell method for measuring CAG-repeat length together with genome-wide RNA expression. We found that the *HTT* CAG repeat expands from 40-45 CAGs to 100-500+ CAGs in HD-vulnerable SPNs but not in other striatal cell types, with these long DNA-repeat expansions acquired at different times by individual SPNs. Surprisingly, somatic expansion from 40 to 150 CAGs had no apparent effect upon gene expression – but neurons with 150-500+ CAGs shared profound gene-expression changes. These expression changes involved hundreds of genes, escalated alongside further CAG-repeat expansion, eroded positive and then negative features of neuronal identity, and culminated in expression of senescence/apoptosis genes. Rates of striatal neuron loss across HD stages reflected the rates at which neurons entered this biologically distorted state. Our results suggest that *HTT* CAG repeats in striatal neurons undergo decades of biologically quiet expansion, then, as they asynchronously cross a high threshold, cause SPNs to degenerate quickly and asynchronously. We conclude that, at any moment in the course of HD, most neurons have an innocuous (but unstable) *huntingtin* gene, and that HD pathogenesis is a DNA process for almost all of a neuron’s life.

## Introduction

Huntington Disease (HD) is a fatal genetic neurodegenerative disease. Most people who inherit HD-causing alleles have no symptoms for decades, then develop uncontrolled movements (chorea), cognitive and psychiatric symptoms; the motor symptoms progress to severe impairment, rigidity and lethality. Persons with HD have severe atrophy of the striatum in which they have lost its principal neurons, striatal projection neurons (SPNs, also called medium spiny neurons or MSNs). No treatments are known to prevent or slow HD.

HD segregates in families in a dominant manner; its genetic cause is an inherited DNA repeat within the *huntingtin* (*HTT*) gene (CAG_n_, encoding polyglutamine) (MacDonald *et al*., 1993). All persons with HD have inherited a germline allele with 36 or more CAGs (36-55 in 98% of cases, 40-49 in 90%), versus the 15-30 CAGs that are inherited commonly (Gusella, Lee and MacDonald, 2021). Longer CAG repeats tend strongly (though with inter-individual variation) to lead to earlier HD onset (Langbehn *et al*., 2004).

It is unknown why stretches of 40 or more CAGs lead almost inevitably to HD. According to a longstanding hypothesis that has guided most biological studies, the encoded polyglutamine causes cumulative cellular damage, with longer polyglutamine stretches more damaging than shorter ones. However, the observation that individuals who inherit two HD-causing alleles do not develop disease symptoms significantly earlier in life (Wexler *et al*., 1987; Myers *et al*., 1989; Cubo *et al*., 2019) has always been challenging to reconcile with the prevailing biological model.

Three central aspects of HD remain unexplained: its cell-type-specific pathology, its long latency, and the series of events by which inherited alleles lead to neurodegeneration.

First, neurodegeneration in HD is highly cell type specific; in the striatum, most SPNs are lost, while interneurons and glia survive, even though all of these cell types express HTT.

Second, HD symptoms generally take decades to manifest. Persons who have inherited common HD-causing alleles (40-45 CAGs) reach adulthood with normal striatal volumes and normal scores on cognitive and motor tests; neuroimaging and fluid biomarkers begin to detect changes only about 10-20 years before onset of clinical motor symptoms (Scahill *et al*., 2020; Tabrizi *et al*., 2022). This long latency is often attributed to a slowly cumulative toxicity of expanded polyglutamine repeats.

A third mystery involves how inherited *HTT* alleles lead to HD. The encoded protein (HTT) has many biological functions; loss, over-expression, and genetic manipulation of *HTT* produce diverse phenotypes in many species and cell types (Saudou and Humbert, 2016). Diverse biological hypotheses are considered plausible for HD, with recent studies focusing on embryonic development (Braz *et al*., 2022), mitochondria (Lee *et al*., 2020), vascular cells (Garcia *et al*., 2022), microglia (Wilton *et al*., 2023), and long-range circuitry effects (Pressl *et al*., 2024).

An important clue may reside in the long-observed phenomenon of somatic mosaicism in HD. The length of the CAG repeat varies somatically; this somatic mosaicism is pronounced in the brain (Telenius *et al*., 1994; Kennedy *et al*., 2003), is greater in neurons than in glia (Shelbourne *et al*., 2007; Mätlik *et al*., 2024), and is greater in persons with earlier-than-expected motor symptom onset (Swami *et al*., 2009). The biological significance of somatic mosaicism in *HTT* has been debated for 30 years, with a dominant view that somatic instability simply modifies the inherent toxicity of inherited *HTT* alleles. However, the recent discovery of common human alleles that modify age-at-onset (Genetic Modifiers of Huntington’s Disease (GeM-HD) Consortium., 2019) suggests that much disease-significant biology may involve changes at a DNA level (Hong *et al*., 2021). HD motor onset is delayed by a synonymous CAG-to-CAA variant (within the CAG repeat) that reduces the repeat’s instability without shortening the encoded polyglutamine (Genetic Modifiers of Huntington’s Disease (GeM-HD) Consortium., 2019; Wright *et al*., 2019). Age of HD onset is also shaped by common genetic variation at many DNA maintenance genes, including *MSH3*, *FAN1*, *MLH1*, *LIG1*, *PMS1,* and *PMS2* (Genetic Modifiers of Huntington’s Disease (GeM-HD) Consortium, 2015; Genetic Modifiers of Huntington’s Disease (GeM-HD) Consortium., 2019). Proteins encoded by these genes affect DNA-repeat stability (Manley *et al*., 1999; Wheeler *et al*., 2003; Dragileva *et al*., 2009; Kovalenko *et al*., 2012; Pinto *et al*., 2013; Tomé *et al*., 2013; Kim *et al*., 2020; Loupe *et al*., 2020; Goold *et al*., 2021).

In this work, to uncover the pathophysiological process in HD and its relationship to the CAG repeat in *HTT*, we developed a molecular approach to measure this repeat at single-cell resolution, concurrent with the same cell’s genome-wide RNA expression. This approach allowed us to recognize biological changes that appear to be driven directly by the CAG repeat.

## Results

### SPN vulnerability, HTT expression and case-control differences in HD

We first used conventional single-nucleus RNA sequencing (snRNA-seq) to analyze RNA expression in 613,670 nuclei sampled from the anterior part of the caudate nucleus – the largest component of the striatum, and the region most affected in HD – from 56 persons with HD and 53 unaffected controls (mean 5,630 cell nuclei per donor). Each nucleus was assigned to one of seven major cell types, based on the RNAs it expressed (**Supplementary Fig. 1**).

The loss of SPNs throughout HD was clear in reduced numbers of SPNs as a fraction of all cells (**Fig. 1a-c**). To analyze together persons with many different ages and inherited CAG-repeat lengths, we used CAG-Age-Product (CAP) score, a common estimate of onset and progression in HD (Zhang *et al*., 2011), which is calculated as *age* * (*inheritedCAGlength* - 33.66). Persons with CAP scores up to 300 (generally corresponding to the long latent period prior to clinical motor onset) tended to have SPN proportions just slightly lower than the average unaffected control brain donor (**Fig. 1b**, **Supplementary Fig. 2**). SPNs appeared to be lost at a much-greater rate in donors with CAP score greater than 400 (donors with manifest HD symptoms), as evidenced by a steep downward slope in the relationship of SPN abundance to CAP score (**Fig. 1b**, **Supplementary Fig. 2**). Almost all persons with a CAP score greater than 600 (corresponding to HD stages with greatly advanced caudate atrophy) appeared to have lost >80% of their SPNs. These results are consistent with findings that caudate atrophy commences subtly about 10-20 years before the onset of motor symptoms, then escalates (Scahill *et al*., 2020; Tabrizi *et al*., 2022).

**Figure 1.**
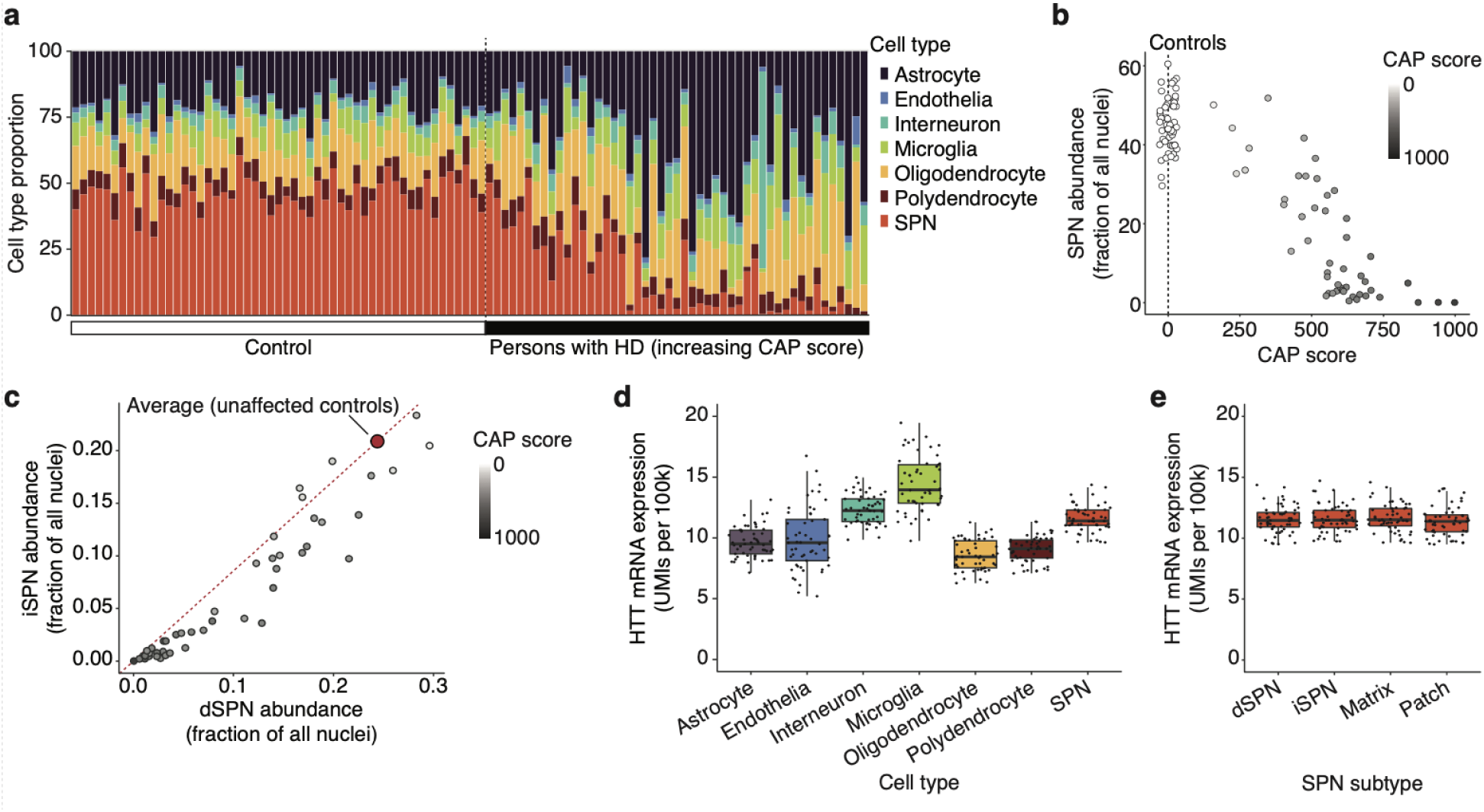
Trajectories of SPN loss in persons with HD. (**a**) Caudate cell-type proportions in each donor, show the loss of SPNs in persons with HD. In the figure, control donors are in the left half, and persons with HD are ordered from left to right by increasing CAP score. (**b**) SPN loss with HD progression (increasing CAP score). Unaffected controls in white; among persons with HD, darker shades of gray represent increasing CAP score. The same data are shown on a log scale in **Supplementary Fig. 2**. (**c**) Decline in iSPNs (D2 SPNs) and dSPNs (D1 SPNs) with HD progression. Gray shading as in **b**. Quantities plotted are fractions of all nuclei (of all cell types) sampled. (**d,e**) Expression of *HTT* transcripts (units: UMIs per 100k) in the nuclei of (**d**) striatal cell types and (**e**) SPN subtypes, among 53 control (unaffected) donors.

Two SPN subtypes defined by their connectivity and gene expression are direct-pathway SPNs (dSPNs) and indirect-pathway SPNs (iSPNs). iSPNs comprised 47% (+/- 6%) of the SPN population in controls, but a smaller fraction in persons with HD (p = 8 x 10^-6^, Wilcox test, Fig. 1c), indicating that iSPNs become vulnerable earlier on average than dSPNs. Since iSPNs inhibit motor programs while dSPNs initiate them, the earlier average loss of iSPNs (which is consistent with stereological measurements (Albin *et al*., 1990)) may underlie the prominence of chorea (involuntary movements) as an early motor symptom in HD (Albin *et al*., 1990). (Relative losses of patch (striosomal) and matrix (extra-striosomal) SPNs more somewhat more variable across individuals, **Supplementary Fig. 3**.)

A longstanding hypothesis for HD pathology involves continuous lifelong damage from a toxic mutant HTT protein; we thus sought to better understand whether *HTT* expression levels (Landwehrmeyer *et al*., 1995; Schilling *et al*., 1995) could help explain the profound vulnerability of SPNs or the more-modest relative vulnerabilities of iSPNs (relative to dSPNs). Biallelic expression levels of *HTT*, as a fraction of all mRNA transcripts, were slightly lower in SPNs than in interneurons, and only modestly higher in SPNs than in most glia (Fig. 1d). *HTT* expression levels in dSPNs and iSPNs were indistinguishable (p = 0.56, paired t-test, Fig. 1e). Individuals varied in *HTT* expression levels, but accelerated SPN loss (relative to CAP score) did not associate with higher *HTT* expression (**Supplementary Fig. 4**).

In every caudate cell type, thousands of genes were differentially expressed (on average) between persons with HD and unaffected individuals (**Supplementary Note 1**). This broadly altered gene expression potentially reflected the consequences of HD, which causes atrophy and de-vascularization of the caudate and a changed context for the remaining cells. Indeed, almost all such changes also associated (in an HD-cases-only analysis) with the extent of a donor’s earlier SPN loss (**Supplementary Note 1)**.

### Measuring somatic CAG-repeat expansion alongside RNA expression

We turned to investigating whether there were cell-autonomous gene expression changes that associated with a cell’s own CAG-repeat length.

We developed a molecular approach for selectively amplifying and analyzing the CAG repeat of *HTT* RNA transcripts (together with cell and molecular barcodes) alongside analysis of genome-wide RNA expression in the same cell nuclei (Fig. 2a). In our approach, each CAG-length measurement is matched (using the cell barcodes) to the gene-expression profile of the cell from which it is derived, and thus to the identity and biological state of that cell (Fig. 2a). We deeply sampled nuclei from the caudate of six persons with clinically manifest HD (**Supplementary Table 1**). We were able to acquire measurements for approximately 10% of SPNs and interneurons (which have the largest nuclei and snRNA libraries) and smaller fractions (2-5%) of non-neuronal cell types.

**Figure 2.**
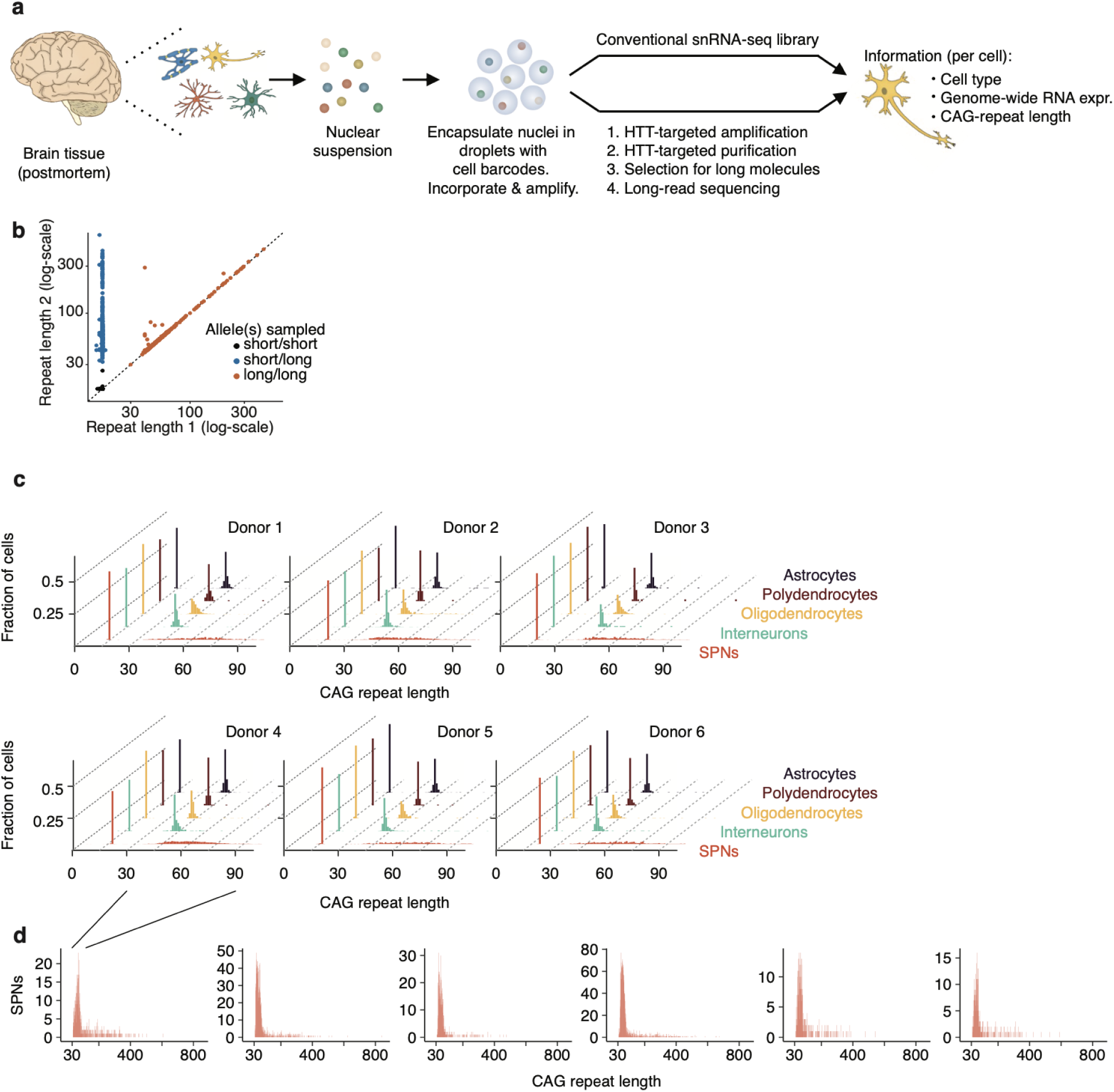
Single-cell analysis of *HTT* CAG-repeat length (via *HTT* RNA transcripts) and genome-wide RNA expression in the same nuclei. (**a**) Molecular approach. Two sequencing libraries are prepared from the same set of barcoded nuclear cDNAs. The first is a conventional snRNA-seq library. The second library samples the CAG-repeat sequence in the first exon of the *HTT* gene and is analyzed by long-read sequencing. The presence of shared cell barcodes in the two libraries allows each CAG-repeat sequence to be matched to the RNA-expression profile of the nucleus from which it was sampled. (**b**) Concordance between pairs of measurements of CAG-repeat length from different *HTT* RNA transcripts (with different UMIs) in the same nucleus (same cell barcode). For each such measurement-pair, the longer of the two CAG-repeat measurements is shown on the y-axis. Nuclei in which both measurements are from the HD-causing allele (orange) make it possible to measure precision and error rate. (**c**) Distributions of CAG-repeat length measurements by donor and cell type; each of the six plots is from a different person with HD. (**d**) For each donor in **c** the distributions of CAG-repeat length measurements in SPNs, showing only the long (HD-causing) allele and the much-wider range of CAG-repeat lengths SPNs attain. See also **Supplementary Fig. 6-8** and **Supplementary Note 2** for additional analyses and visualizations.

We also developed companion analytical approaches. Despite the well-known distorting effects of PCR upon DNA repeats and molecular-size distributions, sets of sequence reads with the same cell barcode and molecular barcode (i.e. from the same *HTT* RNA transcript) exhibited repeat-length consensus (**Supplementary Fig. 5**), which we used in downstream analyses.

When CAG-repeat length could be measured on multiple *HTT* transcripts (with distinct molecular barcodes) in the same nucleus, these measurements also agreed (Fig. 2b).

### Long somatic CAG-repeat expansions in SPNs

The *HTT* CAG repeat exhibited different lengths in different cells and cell types. Astrocytes, oligodendrocytes, polydendrocytes (OPCs), microglia, and interneurons exhibited modest CAG-repeat instability, with almost all cells having a CAG repeat within a few units of the inherited length (Fig. 2c, **Supplementary Note 2**). However, SPNs exhibited extensive somatic expansion of the HD-causing allele (Fig. 2c,d). This pattern was present in all (6/6) of the persons with HD whose caudate we deeply sampled by this approach (Fig. 2c-d, **Supplementary Fig 6,7**). The distinction between SPNs and striatal interneurons was particularly notable, since all are inhibitory (GABAergic) neurons that share a developmental lineage. (Among interneurons, cholinergic interneurons exhibited more expansion than other interneurons, though much less expansion than SPNs (**Supplementary Note 2**).)

Somatic expansion was allele-specific, strongly affecting the HD-causing allele but not the other inherited allele (Fig. 2, **Supplementary Fig. 6**), suggesting that somatic instability is strongly affected in *cis* by an allele’s own CAG-repeat length.

Individuals’ distributions of SPN CAG-repeat lengths exhibited a characteristic shape that visually resembled the profile of an armadillo (**Supplementary FIg. 8**). The bulk of the distribution (the “body”) reflected substantial expansion in almost all SPNs: 95-98% of each donor’s SPNs had expanded beyond the inherited (germline) length, reaching a median CAG-repeat length of 60-73 CAGs (20-31 CAGs longer than the same donors’ germline *HTT* alleles of 40-43 CAGs).

The second feature (the “tail”) involved a prominent minority of SPNs with far-longer expansions (100-500+ CAGs) (Fig. 2d). In SPNs from each donor, this long right tail commenced at about 100 CAGs and tapered only slowly across a wide range (100 to 500+ CAGs), suggesting a second, qualitatively faster phase of somatic expansion that commenced at about 100 CAGs (Fig. 2d). We call these phase A and phase B, and further discuss them in a later section.

Our detection of many SPNs with a long CAG repeats (100-842 CAGs) (Fig. 2d) contrasted with earlier human HD studies, many of which detect repeat expansion only in the 35-100 range (Mätlik *et al*., 2024; Pressl *et al*., 2024). Critical in recognizing the abundance of these long CAG-repeat expansions was incorporating unique molecular identifiers (UMIs) into first-strand cDNAs to address the tendency of subsequent PCR to amplify shorter DNA sequences exponentially more efficiently than longer ones. Notably, a 2003 study, which had utilized small-pool PCR to address this same distorting effect, had also observed many molecules with long (100-1000 CAG) CAG-repeat tracts in HD brain tissue (Kennedy *et al*., 2003). The innovation and observation of (Kennedy *et al*., 2003) were perhaps insufficiently appreciated at the time.

### Expansion to 150 CAGs without consequence

To recognize how a cell’s gene expression is affected by its own *HTT* CAG-repeat length, we identified allelic series of SPNs naturally arising from the mosaicism within each individual person with HD. These six allelic series consisted of 467 to 2,337 SPNs per person, with the CAG-repeat lengths collectively spanning 35 to 842 CAGs. By performing each comparison within-person rather than across people, we controlled for the profound non-cell-autonomous effects of each donor’s disease state (**Supplementary Note 1**).

Surprisingly, our analyses detected no apparent cell-autonomous effects of CAG-repeat expansion across 36 to 150 CAGs; however, the same analyses found that longer expansions (>150 CAGs) were associated with altered expression of hundreds of genes (Fig. 3, **Supplementary Note 3**). This conclusion was supported by many kinds of analyses: simple correlations of gene-expression profiles (Fig. 3a, **Supplementary Fig. 9**); non-parametric statistical tests (as shown in volcano plots and genome-wide distributions of gene-level test statistics; **Fig. 3b,c**, **Supplementary Fig. 10,11**); and regression of gene-expression measurements against CAG-repeat length (**Supplementary Note 3**). None of these analyses detected any cell-autonomous consequences of repeat expansion to 150 CAGs, but all detected profound effects of CAG-repeat expansion beyond 150 CAGs (**Supplementary Fig. 9-12**).

**Figure 3.**
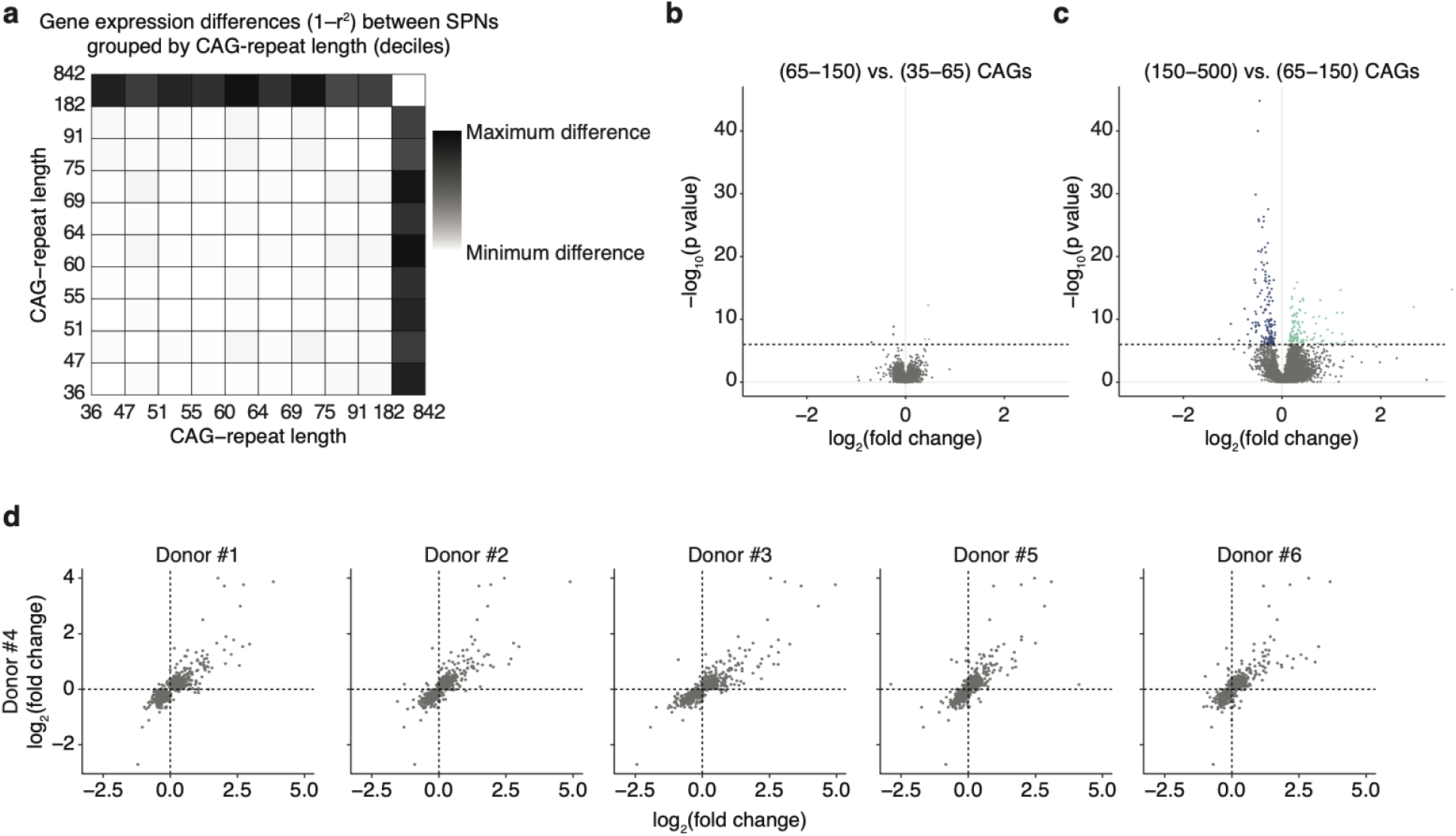
An apparently high CAG-length threshold for effects of the *HTT* CAG repeat. (**a**) Magnitude of gene-expression differences (one minus the correlation coefficient) when comparing sets of SPNs (from the same tissue sample) grouped into deciles based on the CAG-repeat length of their HD-causing *HTT* allele. Gray scale: black indicates maximal difference observed in any comparison; white indicates no difference. **Supplementary Fig. 9** has similar analyses of SPNs from six persons with HD. (**b,c**) Comparisons (volcano plots) of gene expression between sets of SPNs, sampled from the same person with HD but with CAG-repeat lengths in different ranges. p-values (y-axis) are derived from a Wilcoxon text across the individual SPNs in each group. Fold-changes (x-axis) are the log_2_-ratio of the group averages. (**d**) Consistency of long(150)-repeat-expansion associated gene-expression changes among individual persons with HD. Each panel is a pairwise comparison of SPN gene-expression data involving two persons with HD (x- and y-axes), in which the values on the two axes are the log_2_-fold-changes in gene expression when comparing an individual’s SPNs with >150 CAGs to the same individual’s SPNs with <150 CAGs. Genes whose expression levels change significantly with repeat expansion in at least one of the donors are shown. These and more analyses of gene expression in SPNs sampled from six different persons with HD are in **Supplementary Fig. 9-12** and **Supplementary Note 3**.

### Continuously escalating changes (phase C) beyond 150 CAGs

We found that not only the high CAG-repeat-length threshold (∼150), but also the ensuing gene-expression changes, were highly similar from person to person (Fig. 3d, **Supplementary Fig. 12**). Analysis identified hundreds of genes whose expression levels were affected by CAG-repeat length (**Supplementary Note 3**). Notably, these genes exhibited two kinds of relationships to CAG-repeat length: one set exhibited continuous changes in expression levels as the CAG repeat further expanded beyond 150 CAGs; another set exhibited more-discrete and dramatic changes in a specific subset of SPNs with still-longer CAG-repeat expansions (generally >250 CAGs).

More than 500 genes exhibited incipient and escalating gene-expression distortion to the extent the CAG repeat had beyond 150 units (Fig. 4a, **Supplementary Fig. 13, 14**). We refer to this as phase C (continuous change), and to the affected genes as C- (down-regulated) and C+ (up-regulated) genes.

**Figure 4.**
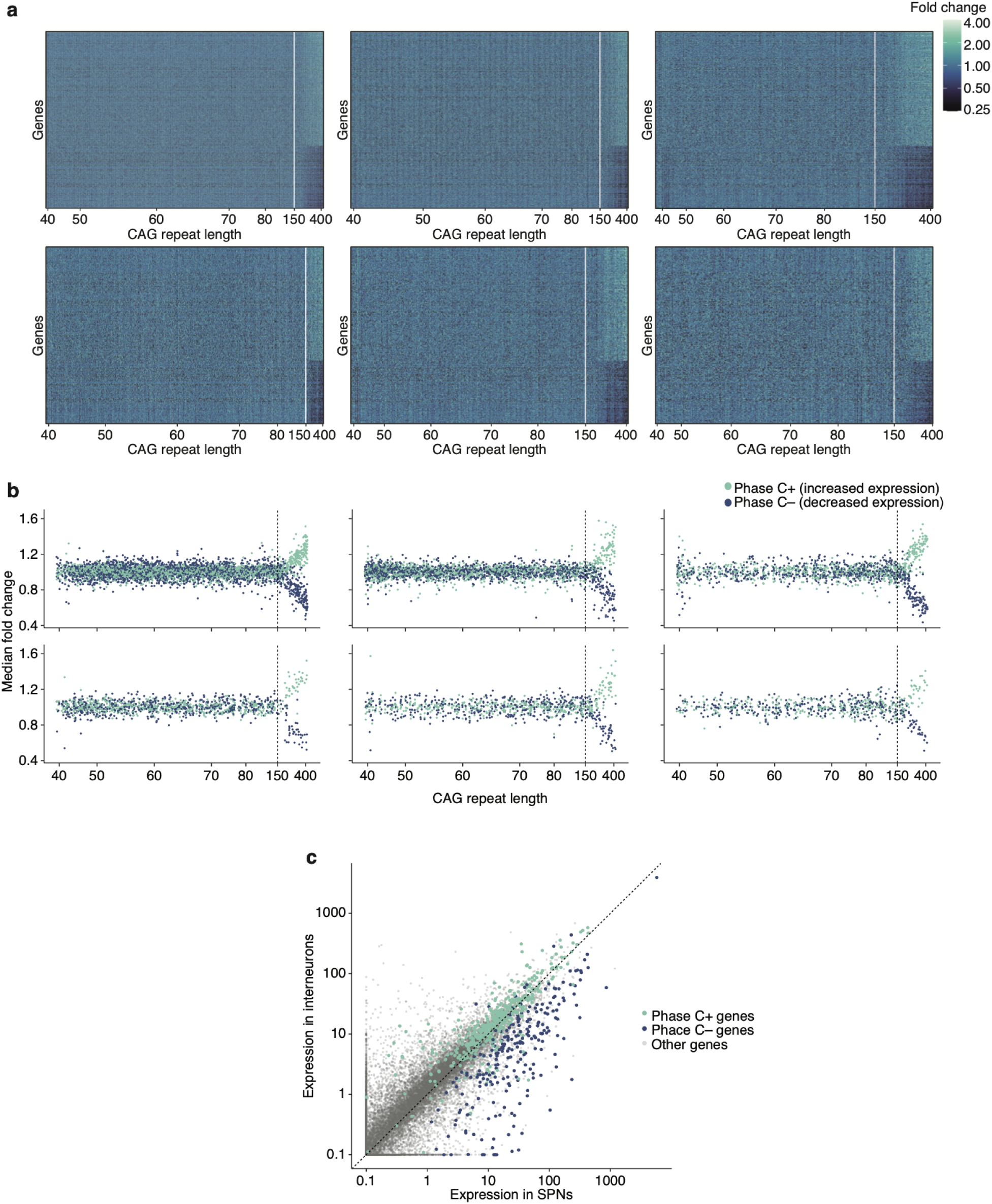
Gene-expression change in SPNs with somatic expansion beyond 150 CAG repeats. (**a**) On each plot, a specific donor’s individual SPNs are ordered from left to right by their CAG-repeat length (thus corresponding to the columns of the heatmap). Each row shows expression data for a specific gene in each of these SPNs. (The genes shown are those found to change in expression with repeat expansion.) Shades of each facet show the level of expression of that gene in that SPN, relative to the average SPN with repeat length < 150 in that donor (lighter shades: elevated expression; darker shades: reduced expression). Example trajectories for individual genes are in **Supplementary Fig. 14**. **(b)** As in **a**, on each plot, individual SPNs are ordered from left to right by their CAG-repeat length. Each SPN is represented by both a blue and a green point: blue points show the median fold-change of a set of 203 genes that decrease in expression with repeat expansion (C-genes). Green points show the median fold-change of a set of 369 genes that increase in expression with repeat expansion (C+ genes). (**c**) Gene-expression features of SPN identity and their relationship to phase C gene-expression changes. Gray points: all genes (cell-type-specific expression levels in unaffected individuals, measured in UMIs per 100k). Colored circles: Genes whose expression levels decline (blue) or increase (green) in SPNs as the *HTT* CAG repeat expands beyond 150 units (in phase C).

Repeat-length-associated expression changes were almost undetectable at 150-180 repeats, but analyses that drew upon all the genes together indicated that these changes had commenced by about 150 repeats (Fig. 4b, **Supplementary Note 3**). This pattern was shared across persons with HD (Fig. 4b) and was shared by direct and indirect SPNs, and by patch (striosomal) and matrix (extra-striosomal) SPNs (**Supplementary Fig. 13**).

The genes whose expression declined in SPNs with long repeat expansions were among the most strongly expressed genes in healthy SPNs, and were genes whose high expression levels in SPNs distinguished SPNs from other types of inhibitory neurons (Fig. 4c). These included *PDE10A, PPP2R2B, PPP3CA, PHACTR1, RYR3*, and more than 100 other genes that normal SPNs express more strongly than striatal interneurons do (Fig. 4c). This suggests that a core biological change in phase C involves the erosion of SPN-identity features that distinguish SPNs from other kinds of inhibitory neurons. Many of these genes also have important physiological functions; for example, genes encoding the potassium channel subunits KCNA4, KCNAB1, KCND2, KCNH1, KCNK1, KCNQ5 and KCTD1 all declined in expression during phase C, a change that might affect SPN physiology.

*HTT* expression itself did not associate with an SPN’s own CAG-repeat length (**Supplementary Fig. 15**), though this does not preclude altered post-transcriptional processing (Sathasivam *et al*., 2013) that snRNA-seq does not measure.

Though the relationship of phase-C changes to a cell’s own CAG-repeat length was strong and clear (Fig. 4), such changes appear to have been hard to recognize in bulk and sorted-SPN studies because they arise asynchronously in sparse SPNs. Earlier studies have focused primarily on changes which our own analysis suggested were the result of earlier SPN loss, as they were experienced equally by all remaining SPNs (regardless of CAG-repeat length) and to an extent predicted by a donor’s earlier SPN loss (**Supplementary Note 1**).

We found stronger alignment between our findings and analyses of a specific HD mouse model (Q175), which begins life with a CAG-repeat tract of about 175 CAGs in all cells. In such mice, SPNs, interneurons and glia all exhibit reduced expression of genes that distinguish them from one another (Malaiya *et al*., 2021). The presence of this long (175 CAG) repeat in all cell types (rather than just SPNs) may also explain the diverse cell-type-specific pathologies, including vascular and oligodendrocyte pathologies (Ferrari Bardile *et al*., 2019; Garcia *et al*., 2022), exhibited by Q175 mice.

### De-repression crisis (phase D)

A distinct set of genes that are normally repressed in SPNs also exhibited repeat-length-dependent change, but with a different pattern. These genes remained repressed even in most SPNs with long expansions (>150 repeats), but tended to become de-repressed in those SPNs in which the phase-C changes had progressed to the greatest degree (**Fig. 5a**,b). In the cells in which de-repression had occurred, it tended to involve very many genes concurrently (Fig. 5a). We refer to this state as “de-repression crisis” (phase D).

**Figure 5.**
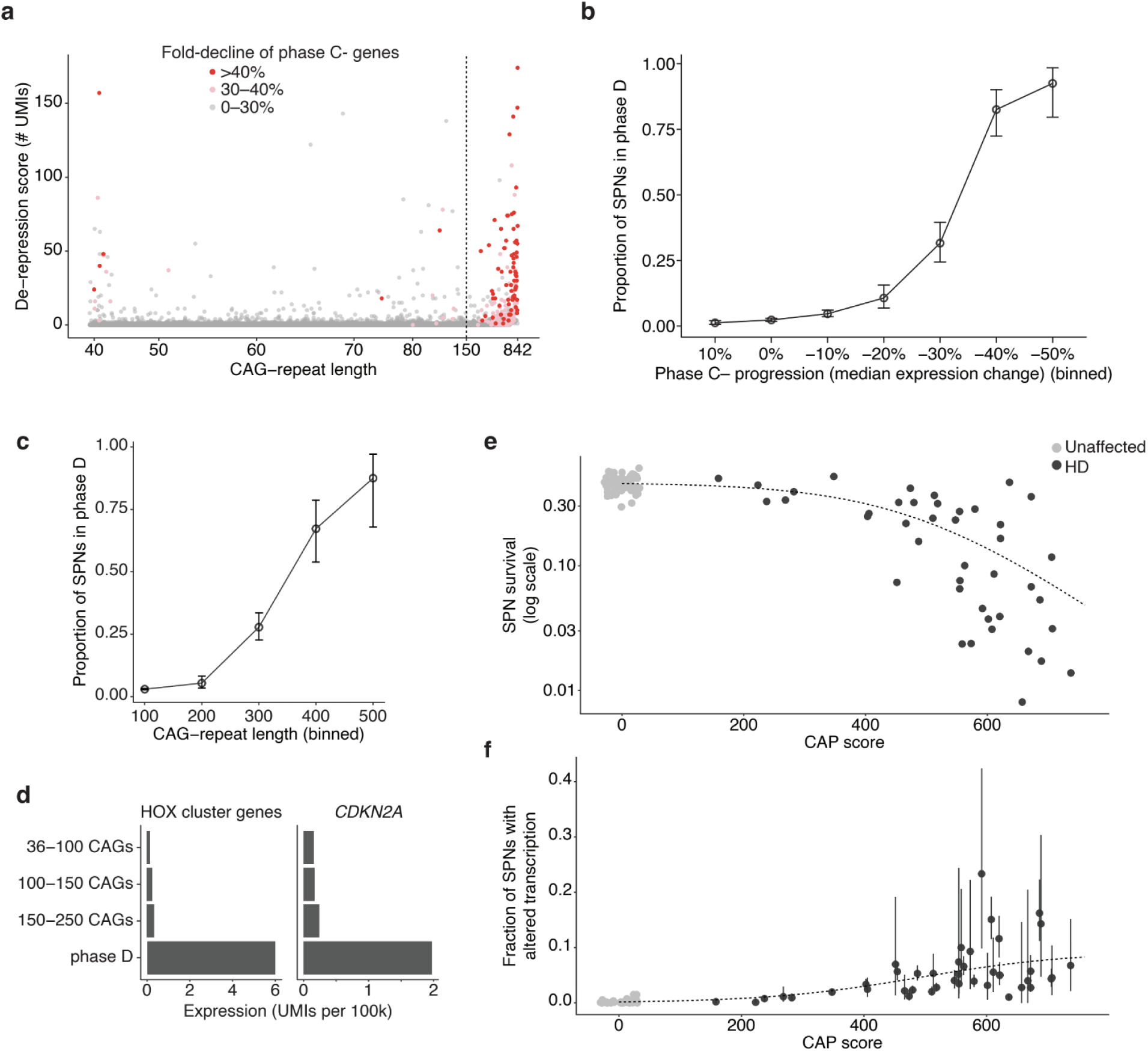
De-repression crisis (phase D) and subsequent SPN elimination (phase E). (**a**) De-repression of genes that are normally silent in SPNs. Points represent individual SPNs; red and pink points are those SPNs whose phase-C expression changes (Fig. 4b) have progressed beyond the threshold values shown. y-axis: de-repression score, the number of transcripts (UMIs) detected from 173 “phase D” genes that are normally silent in SPNs. An additional visualization of this relationship is in **Supplementary Fig. 16**. (**b**) Fraction of SPNs exhibiting this de-repression phenotype, in relationship to the increasing dysregulation (reduced expression) of the phase C-genes. (**c**) Fraction of SPNs exhibiting de-repression phenotype, in relation to CAG-repeat length (**d**) Expression of HOX cluster genes (left) and *CDKN2A* (right) in SPNs of persons with HD (units: UMIs per 100k in a metacell combining the individual SPNs). (**e,f**) SPN loss and transcriptionopathy as HD progresses. (**e**) Relationship of SPN survival (shown on a logarithmic scale) to HD progression as indexed by CAP score. The dashed curve shows a logistic function fit to the SPN-survival data; its slope (derivative) estimates average rates of SPN loss. Data from 56 control (unaffected) donors are shown on the left side of the plot with CAP scores of zero. (**f**) Relationship of the frequency of SPN phase-C transcriptionopathy (y-axis) to HD progression as indexed by CAP score. The dashed curve is the negative of the derivative of the SPN-survival curve from the panel above (i.e., is proportional to the estimated rate of SPN loss).

Phase D was associated with still-longer CAG repeats (Fig. 5a). De-repression was rare (<4%) even among SPNs with 150-250 CAGs, but it became very common (>50%) in SPNs with 350 or more CAGs (Fig. 5c). Notably, in nuclei in which phase-D changes were detected, the number of de-repressed genes bore little relationship to CAG-repeat length (**Supplementary Fig. 16**), a pattern distinct from the phase-C changes, which were well-predicted by an SPN’s CAG-repeat length at the time of analysis (Fig. 4b). We interpret this to mean that, while phase-C changes proceed on a time scale similar to that of fast CAG-repeat expansion (beyond 150 CAGs), phase-D changes progress with far-faster kinetics once underway.

The 173 genes we found to be de-repressed in phase D had distinct biological features in common. They included most of the genes at the HOXA, HOXB, HOXC, and HOXD loci, as well as noncoding RNAs (*HOTAIR, HOTTIP, HOTAIRM1*) at these same loci. These genes are involved in cell specification in the brain and other organs (Hobert, 2021) and are normally expressed during early embryonic development but not in adult neurons. The de-repressed genes at other loci included dozens of transcription-factor genes (including *FOXD1, IRX3, LHX6, LHX9, NEUROD2, ONECUT1, POU4F2, SHOX2, SIX1, TCF4, TBX5, TLX2, ZIC1, ZIC4*) that are normally expressed in other neural cell types but not in SPNs.

The de-repression of so many transcription-factor genes might lead to expression of genes normally expressed in other neural cell types. Indeed, phase-D SPNs expressed many genes that are normally expressed in interneurons (*CALB2, SST, KCNC2*), in glutamatergic (excitatory) neurons (*SLC17A6, SLC17A7, SLC6A5*), in astrocytes (*SLC1A2*), in oligodendrocyte progenitor cells (*VCAN*), or in oligodendrocytes (*MBP*). These changes suggested that SPNs in phase D were losing negative as well as positive features of SPN cell identity.

Two of the most strongly de-repressed genes in phase D were *CDKN2A* (Fig. 5d) and *CDKN2B*, which encode proteins (p16(INK4a) and p15(INK4b)) that promote senescence and apoptosis (Igney and Krammer, 2002; Gil and Peters, 2006; Herranz and Gil, 2018; Yuile *et al*., 2023). Ectopic expression of *Cdkn2a* is toxic to neurons (Finneran *et al*., 2023). De-repression of *CDKN2A* and *CDKN2B* in phase-D SPNs may be an imminent cause of their death.

Interestingly, inactivation of the Polycomb Repressor Complex 2 (PRC2) in adult mice causes a similar set of gene-expression changes, including de-repression of Hox genes, other transcription factors, and *Cdkn2a* and *Cdkn2b*, leading within months to SPN loss, motor function decline and lethality (von Schimmelmann *et al*., 2016).

### Elimination phase (phase E)

The above results suggested that the transcriptional changes in long-repeat (phase C,D) SPNs might lead to their death. Though we cannot observe the same SPNs at multiple points in time, we sought to learn from comparisons across donors who passed away at different stages of caudate atrophy and SPN loss. To do this, we used CAP score as a measure of HD progression (as in Fig. 1) in order to bring donors with a variety of ages and inherited CAG-repeat lengths into a single analysis (**Fig. 5e**,**f**).

Across HD progression, the rate of SPN loss can be estimated from the slope of the decline in SPN abundance (on a logarithmic scale) (Fig. 5e**,f**). This inferred rate of SPN loss exhibited clear proportionality to the fraction of donors’ SPNs whose RNA expression indicated the presence of phase-C transcriptional changes (Fig. 5f, **Supplementary Fig. 17**). In addition, in donors with precocious SPN loss (relative to CAP score), larger fractions of SPNs exhibited this transcriptionopathy (**Supplementary Fig. 18**).

### Insights from computational modeling of DNA repeat-expansion dynamics

Our experimental results were hard to reconcile with conventional biological models of HD in which most or all SPNs endure a toxic mutant HTT *simultaneously*. Could a model of *sequential* SPN toxicity – in which, at any one time, most SPNs have a biologically innocuous HTT whose repeat length is far below a high toxicity threshold – plausibly explain the relentless loss of SPNs in HD (Fig. 1)? Could the decades-long latent period before symptom onset be reconciled with the subsequent, fast loss of SPNs (Fig. 1)?

To address these and other questions, and to better appreciate the dynamic processes that might give rise to clinical observations and end-of-life biological measurements, we turned to computationally modeling of repeat-expansion dynamics over the human lifespan, seeking to understand whether simple models based on an emerging understanding of DNA repeat-expansion mechanisms (Iyer and Pluciennik, 2021; Phadte *et al*., 2023) (Fig. 6a) would generate repeat-length distributions and cell-loss trajectories consistent with our experimental results.

**Figure 6.**
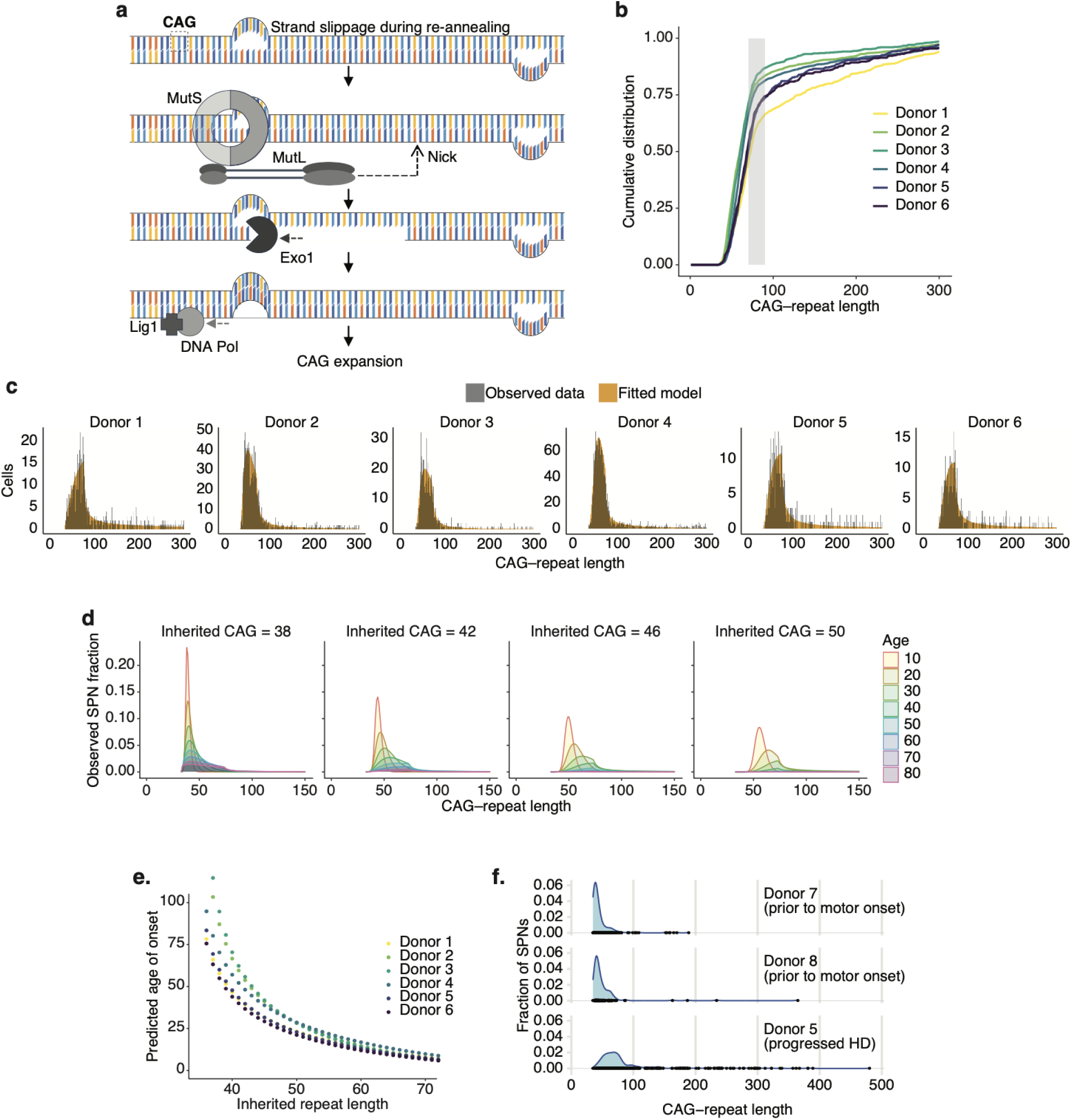
Computational modeling of somatic CAG-repeat expansion dynamics. (**a**) Schematic illustrating mechanisms (established in earlier work, (Iyer and Pluciennik, 2021)) for non-replicative DNA-repeat expansion in post-mitotic cells. Extrahelical DNA extrusions (“slip-out” structures) form from mispairing within the CAG-repeat tract after strand separation (e.g. due to transcription). The MutSβ complex can bind to these transient structures, initiating DNA excision and resynthesis, which (when initiated on the strand opposite the slip-out) can result in the incorporation of an extra repeat unit. Occurring multiple times across a human lifetime, this mutational process results in a progressive expansion of the DNA repeat. (**c**) Cumulative distributions of CAG-repeat length measurements in SPNs from six deeply sampled donors. The gray shaded region highlights the range (70-90 CAGs) over which somatic expansion appears to greatly accelerate. (**c**) Distributions of CAG-repeat length measurements in SPNs from these same donors (black) overlaid with the results of stochastic models (orange) for which a few key parameters (such as mutation rate) have been fitted to each donor’s repeat-length and SPN-loss data. (**d**) Effect of changing a single variable (inherited/germline CAG-repeat length) in the model for a typical donor, keeping the other fitted parameters fixed. Each curve indicates the predicted CAG-repeat length distribution for surviving SPNs at each decade (ages 10 to 80). (**e**) Model-estimated relationship between inherited germline CAG-repeat length and age at clinical motor onset. As a proxy for age of onset, we used the predicted time at which 25% of a donor’s SPNs have been lost. We estimated each donor’s age of onset proxy at different hypothetical inherited repeat lengths. The shapes of the resulting curves approximate the known relationship between inherited repeat length and age of HD onset. (**f**) Observed CAG-repeat length distributions for two donors with HD-causing alleles who passed away prior to clinical motor symptom onset based on review of their medical records (**Supplementary Table 1**). Data from a typical symptomatic donor (donor 5) is shown at bottom for comparison (distributions for several other symptomatic donors are in Fig. 2d).

In post-mitotic cells, DNA-repeat length-change mutations are thought to result from occasional strand misalignment (mispaired repeats) after transcription or transient helix destabilization (Corless and Gilbert, 2016). Mispaired repeats create extrahelical extrusions (“slip-out” structures) (Fig. 6a). Small extrahelical extrusions are recognized by DNA mismatch repair (MMR) complexes, which initiate repair pathways (Iyer and Pluciennik, 2021) that involve nicking, excision, and resynthesis of one of the two strands. If the two slip-out structures are farther apart than this excision distance, then resynthesis results in a length-change mutation (Fig. 6a) – an expansion or contraction, depending on which strand has been nicked and excised. (MMR complexes have a strand bias that leads to expansions more frequently than contractions (Phadte *et al*., 2023)). Experimental observations indicate that repeat expansion tends to occur in small increments (Dragileva *et al*., 2009; Goold *et al*., 2021).

Our simulations adhered as closely as possible to this emerging understanding. We assumed that all SPNs initially had the same (germline) *HTT* allele; that length-change mutations were stochastic expansions or contractions of a small number of CAG units; that the likelihood of mutation increased with repeat length; and that SPN loss occurred among SPNs with >150 repeats. We found mutation-rate and expansion-contraction-bias parameters that optimized the likelihood of the observed data from each person with HD, including the distribution of SPN CAG-repeat lengths and SPN loss at the age of death and brain donation. These simulations are described in detail in **Supplementary Note 4**, and available as <u>Supplementary Movies</u>.

The most challenging aspect of the repeat-length data to explain was its armadillo shape (**Supplementary Fig. 8**) – the simultaneous presence of a large majority of SPNs with 40-100 CAGs, and a small minority of SPNs with far more (100-800+) CAGs. All the donors we analyzed exhibited this transition across the same CAG-repeat length range of about 70-90 CAGs (Fig. 6b). Models in which the increase in the mutation rate was a linear, quadratic, higher-order polynomial, or log-normal function of repeat length did not generate this shape. However, models with two phases of expansion – a slow phase (phase A) that transitioned into a much-faster phase (phase B) – generated data that closely matched the experimental data (Fig. 6c, **<u>Supplementary Movie 1</u>**). Our models estimated this transition as occurring over a similar repeat-length interval (70-90 CAGs) in each donor, with the mutation rate increasing at least ten-fold over this range (beyond its general pattern of continuous increase with repeat length). We note that, at this length scale (70+ CAGs, 210+ bp), otherwise-mobile slip-out structures (Fig. 6a) may with increasing likelihood be separated by an intervening nucleosome, greatly increasing the likelihood that they are surveilled by MMR complexes before they resolve on their own.

Fitting the experimental data did not require assuming single-cell heterogeneity in mutation rates: we found that asynchronous SPN toxicity could be explained simply by the asynchronous passage of SPNs from phase A to the subsequent, faster phase B. This asynchronicity arose from that (i) length-change mutations were initially rare events (occurring less than once per year per cell across 36-55 CAGs), and (ii) such mutations, upon occurring, increased the probability of subsequent mutations.

A fundamental relationship in HD is the association of longer inherited alleles with symptom onset earlier in life – a relationship that is steep in the 36-45 CAG range (across which each extra inherited CAG accelerates symptom onset by more than a year) and has long been thought to reflect increasing mHTT toxicity in this range. Our simulations also produced this relationship, but for a different reason: slightly longer inherited alleles bypassed the CAG-repeat lengths at which somatic expansion is most infrequent (occurring less than once per year) (Fig. 6d**,e**, **<u>Supplementary Movie 2</u>**). We note that this was previously predicted on theoretical grounds by Kaplan et al. (Kaplan, Itzkovitz and Shapiro, 2007).

Simulation results suggested that the earlier loss of iSPNs relative to dSPNs (Fig. 1c) – which had not been explained by *HTT* expression levels (Fig. 1f) – may instead be explained by a modestly higher (∼15%) rate of somatic expansion in iSPNs (**<u>Supplementary Movie 3</u>**).

A long-standing mystery about HD involves the long latent period (generally decades) in which persons have no apparent symptoms (ISS Stage 0-1, see (Tabrizi *et al*., 2022)). Our simulations predicted that persons in this stage might in fact have substantial somatic expansion, but with almost all SPNs still in phase A. To test this, we analyzed caudate tissue from two persons with HD who had passed away and contributed their brains for research prior to clinical motor diagnosis and/or without apparent neuropathology upon autopsy. Distributions of CAG-repeat lengths in their SPNs indeed exhibited substantial somatic expansion, but few cells with long (>100) expansions (Fig. 6f).

We also found that explaining a long-puzzling feature of HD – the transition from slow to rapid atrophy of the caudate – did not require common assumptions of a non-cell-autonomous disease-escalating process (such as inflammation or spreading prions). Rather, the period of more-rapid decline corresponded to the period in which the bulk of a person’s SPNs reached the end of phase A and more quickly traversed the subsequent, pathological phases (whereas only a few, precociously expanding SPNs did this during the latent stage).

Our simulations suggest that the average SPN in a person with the most common HD-causing inherited allele (42 repeats) spends 96.4% (s.d. 2.0%) of its life below the threshold of 150 CAG repeats at which distorted gene expression appears to commence – i.e., with what our experimental results suggest is an innocuous mutant *huntingtin* gene.

### A pathogenesis model: Extra-long repeats acquire toxic effect (ELongATE)

Our results suggest that an SPN’s own CAG repeat becomes toxic only when quite long (>150 CAGs), and that this repeat length is necessary and sufficient for pathology. We propose a model of HD pathogenesis involving a series of phases driven cell-autonomously by a neuron’s own expanding *HTT* allele (Fig. 7, **Table 1**). We call this dynamic ELongATE (extra-long repeats, acquired toxic effect).

**Figure 7.**
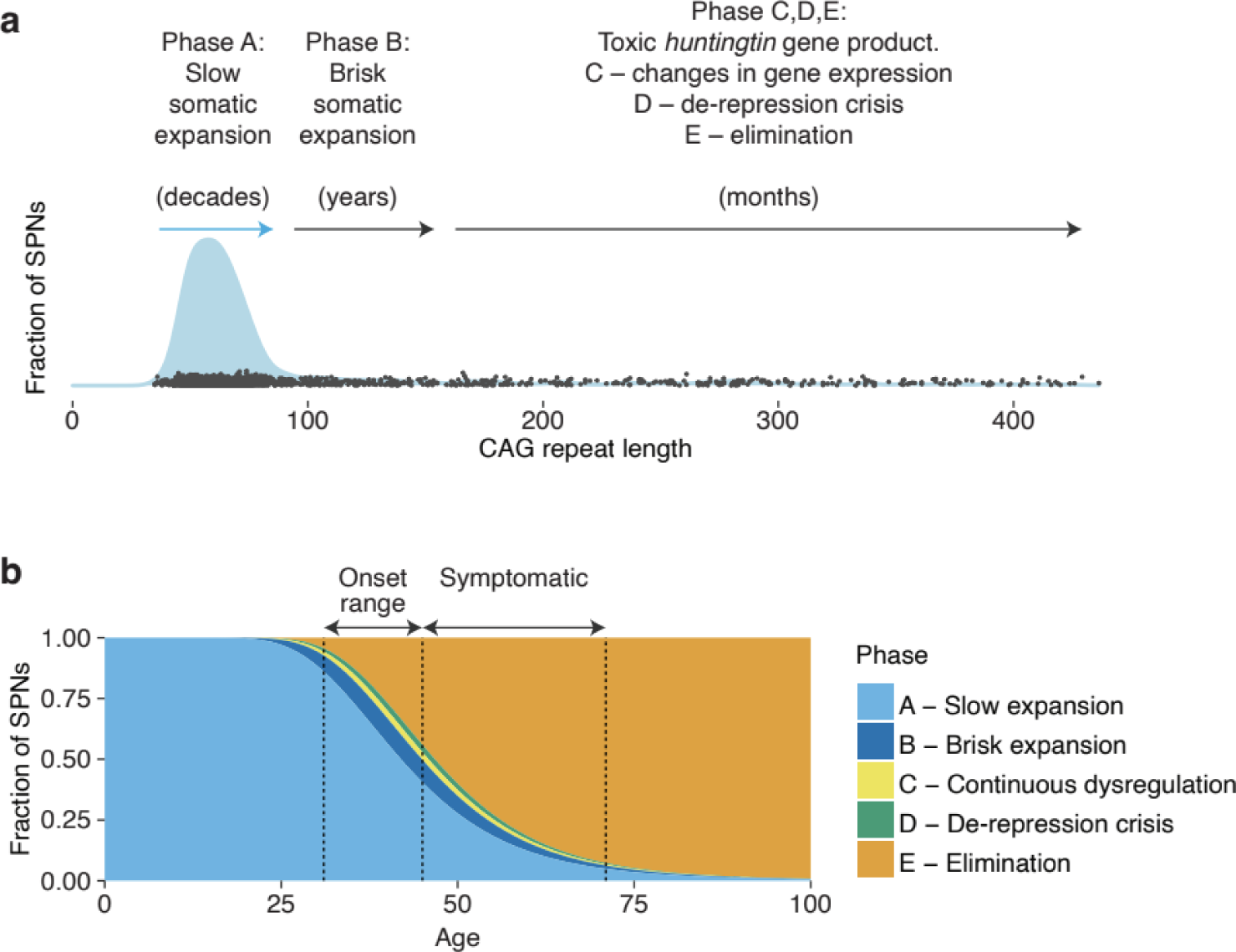
ELongATE (extra-long repeats acquire toxic effect): a model for neuropathology in HD. (**a**) Individual neurons pass asynchronously through five key pathological phases, spending >95% of their lives in a long period of DNA-repeat expansion (a “ticking DNA clock”, phases A and B) with a biologically harmless (but unstable) *HTT* gene. Individual neurons asynchronously exit phase A and proceed through the subsequent, faster phases **(b)** Prediction of the fraction of SPNs in each of the five phases (**Table 1**), across the latent, peri-onset and then progressive stages of HD. The estimated trajectories are based on the data from a representative donor. The indicated ranges for clinical motor onset and escalating symptoms are approximate. The illustrated onset range, representing between 20% to 50% SPN loss, as inferred from available medical records in the patients we have analyzed.

**Table 1.** Proposed phases in HD pathology at the single-neuron level. Time estimates are for persons who inherit the more common HD-causing alleles (40 to 45 CAG repeats).

| Phase | CAG-repeat length | Somatic expansion dynamic | Time in phase (est.) | Gene expression changes associated with neuron's own *HTT* CAG repeat length (relative to nearby neurons in the same tissue) |  |  |
| --- | --- | --- | --- | --- | --- | --- |
|  |  |  |  | Genes affected | Dynamic | Functional theme |
| **A** slow DNA-repeat expansion | 36 to 80 | Slow (initially <1/yr) but increasing | Decades | — | — | — |
| **B** brisk DNA-repeat expansion | 80 to 150 | Fast | Years | — | — | — |
| **C** continuous gene-expression change | > 150 | Fast | Months | Hundreds | Gradual, escalating | Cell-identity erosion |
| **D** de-repression crisis | >> 150 | Fast | Months | Hundreds | Discrete, concurrent | De-repression crisis |
| **E** eliminated |  |  |  | (Atrophy-associated gene-expression changes in surviving cells) |  |  |

In the first phase (phase A, when a neuron has 36 to 80 CAGs), an SPN undergoes decades of slow-but-accelerating repeat expansion. We estimate that an SPN takes 50 years (on average) to expand from 40 to 60 CAGs, then another 12 years to expand from 60 to 80, but with variability both cell-to-cell and person-to-person (**Supplementary Note 2**). Phase A could be compared to a slowly and capriciously ticking DNA clock.

As a neuron enters the second phase (phase B, 80 to 150 CAGs), the rate of expansion greatly accelerates, and the tract may now expand to 150 CAGs in just a few years. Still, as in phase A, the neuron’s *HTT* CAG repeat does not appear to affect its own gene expression. Phase B could be compared to a more rapidly, predictably ticking DNA clock.

As a neuron enters the third phase (phase C, 150+ repeat units), hundreds of genes begin to change in expression levels. These changes are initially tiny, but they escalate alongside further repeat expansion (Fig. 3**,4**), eroding gene-expression features of SPN identity (Fig. 4c).

In its fourth phase (phase D), an SPN de-represses scores of genes that are typically expressed in other neural cell types or in embryonic development. Phase-D neurons also de-repress *CDKN2A* and *CDKN2B*, which encode proteins that promote senescence and apoptosis.

In the final phase, an SPN is eliminated (phase E). Such cells do not appear in CAG-length and gene-expression data, but the effect of their earlier loss is apparent in effects on gene expression in remaining cells of all types (including SPNs) (**Supplementary Note 1**).

Importantly, individual SPNs enter the fast phases (B,C,D,E) at different times, an asynchrony which our modeling suggests can be explained largely by the variable amounts of time that individual neurons take to traverse phase A. Phase A introduces this asynchrony because each neuron’s expansion results from low-frequency stochastic length-change mutations (initially occurring less than once per year), with each expansion event increasing the likelihood of subsequent such events.

## Discussion

### Three biological questions

Biological research on HD has long been animated by three questions. What is toxic to cells about the inherited alleles that cause HD? Why is this toxicity so cell-type-specific? And why are HD symptoms preceded by a decades-long latent period? Our experiments and analyses suggest surprising answers to all three questions.

The surprising answer to the first puzzle – the biological nature of the toxicity of inherited HD-causing alleles – is that such alleles are in fact innocuous to SPNs, and remain so even after decades of somatic expansion (phase A in Fig. 7). We found no effect of CAG-repeat length on SPN gene expression across a wide range of repeat-lengths (36-150) including those inherited by almost all patients – but we found profound, and likely quite toxic, gene-expression changes in SPNs with longer (>150) repeats. A potential interpretation is that the apparent threshold for an inherited *HTT* allele to be disease-causing (36-40 CAGs) reflects not that such alleles encode toxic RNAs or proteins, but that they are sufficiently unstable as to be likely to expand beyond 150 repeats within a human lifetime. We propose that the key question is not, “What is toxic about inherited *HTT* alleles?”, but rather, “What toxicity is acquired with expansion beyond about 150 repeats?“. Future molecular research on *huntingtin* RNA and protein might optimally focus on phenomena that change at long (∼150) repeat lengths as opposed to the modest repeat lengths (35-100) observed in most cells.

We propose that the answer to the second question – the cell-type specificity of cell death in HD – is that the CAG repeat reaches the high toxicity length threshold only in certain cell types (Fig. 2c). The key question is perhaps not, “why are SPNs more vulnerable to a toxic HTT protein”, but rather “why does somatic instability vary by cell type?”

The surprising answer to the third puzzle – the apparently slow toxicity caused by mutant HTT – may be that once a neuron develops a harmful *HTT*, that neuron’s decline is not actually slow. Our results suggest that, once the toxicity threshold is crossed and cell-autonomous biological changes start, these changes progress to cell death over months rather than decades – one to two orders of magnitude faster than previously thought. Individual neurons thus tend to experience their own *huntingtin* toxicity asynchronously.

Scores of phenomena have been described in animal and cellular models of HD and proposed to explain or contribute to HD pathogenesis. A long-standing challenge has been to identify which of these are disease-driving mechanisms, which are reactive mechanisms, and which arise only in models but are not features of HD in humans. The potential centrality of an ELongATE dynamic (Fig. 7) in HD is strongly supported by human-genetic findings of HD “modifiers”, common alleles that affect the age at which HD motor symptoms commence.

Known modifier effects arise from *MSH3, FAN1, MLH1, LIG1, PMS1*, and *PMS2 (Genetic Modifiers of Huntington’s Disease (GeM-HD) Consortium., 2019)*, genes that are functionally united not only by roles in DNA repair, but by more-specific effects upon the long-term stability of DNA repeats (Manley *et al*., 1999; Wheeler *et al*., 2003; Dragileva *et al*., 2009; Kovalenko *et al*., 2012; Pinto *et al*., 2013; Tomé *et al*., 2013; Kim *et al*., 2020; Loupe *et al*., 2020; Goold *et al*., 2021; Phadte *et al*., 2023).

It will be important to understand whether the neuron-autonomous, CAG-repeat-driven disease dynamic that we have described in the striatum also explains HD pathology in other brain areas, where other factors (such as connectivity to SPNs) have been proposed to be necessary for pathology.

### Therapeutic implications

The most significant implication of our findings may be for developing therapies for HD and perhaps other DNA-repeat disorders.

The focus of almost all therapies in advanced clinical development for HD is on lowering HTT expression – by diverse approaches that include antisense oligonucleotides, small interfering RNAs, splicing modulation, and gene editing (Tabrizi, Ghosh and Leavitt, 2019). Under conventional models for HD pathology, HTT lowering has a compelling rationale: if inherited HD-causing alleles encode a toxic protein (or become toxic after just modest somatic expansion), and if the cell-biological process by which such alleles lead to neuronal death is decades-long, then even a partial reduction in HTT production might greatly delay disease. However, HTT-lowering treatments have so far been unsuccessful in HD clinical trials (*Nature Reviews Drug Discovery*, 2021, *Science*, 2021; McColgan *et al*., 2023).

Our model for HD pathogenesis (Fig. 7, **Table 1**) suggests two potentially important challenges for the HTT-lowering therapeutic approach. First, at any time, very few SPNs may actually have a toxic HTT protein from whose lowering they might benefit (Fig. 7). (At the same time, most neurons may be deriving positive biological function from HTT (Burrus *et al*., 2020).) Second, even once an SPN arrives at cell-biological toxicity (phases C,D in Fig. 7) and may benefit from HTT lowering, its expected lifetime (if untreated) may be months rather than decades. In short, HTT-directed therapeutic efforts will need to address the possibility that HTT toxicity is brief, asynchronous and intense, rather than long, synchronous and indolent.

Our conclusion that HD pathogenesis is a DNA process for >95% of a neuron’s life (Fig. 7) suggests potentially greater focus on trying to slow somatic expansion. Experimental reduction in the function of MMR genes (including *MSH3*, *MSH2*, *MSH6* and *PMS1*) can stabilize DNA repeats in mice and/or cultured cells (Manley *et al*., 1999; Wheeler *et al*., 2003; Dragileva *et al*., 2009; Kovalenko *et al*., 2012; Pinto *et al*., 2013; Tomé *et al*., 2013; Ferguson *et al*., 2023) and thus might pre-empt the somatic-genetic cause of HD pathology. However, much uncertainty has surrounded the therapeutic window that such an approach could have. Our results suggest that the therapeutic window might be quite wide: if a cell spends 95% of its life in phase A, then even modestly slowing somatic expansion might substantially postpone HD symptom onset.

What about persons who already have early HD symptoms? Surprisingly, our results predict that even when a person with HD has lost 25% of their SPNs, more than 90% of still-living SPNs still have a huntingtin gene that is not yet biologically harmful (Fig. 6**,7**, **Supp. Note 3**). A future somatic-expansion-directed therapy thus might be able to slow or stop HD progression even in persons who already have early HD symptoms. This would allow the efficacy of such therapy to be evaluated in patients with HD symptoms, a faster and more straightforward path to clinical evaluation than a long-term prevention trial.

### Implications for other DNA-repeat disorders

The dynamic we have described might also apply in other DNA-repeat disorders. More than 40 human diseases are caused by inherited expansions of DNA repeats in protein-coding sequences, introns, UTRs, or promoters (Paulson, 2018; Rajagopal *et al*., 2023). Several of these diseases involve age-associated mosaicism and mid-life onset (Monckton *et al*., 1995; Morales *et al*., 2012, 2020; Campion *et al*., 2022). Many – including Myotonic dystrophy 1, X-linked dystonia Parkinsonism, Friedrich ataxia, and six forms of spino-cerebellar ataxia (SCA1, SCA2, SCA3, SCA6, SCA7, SCA11) – are also (like HD) delayed or hastened by common genetic variation at genes that regulate somatic DNA-repeat stability (Morales *et al*., 2016; Laabs *et al*., 2021; Rajagopal *et al*., 2023). If these disorders share a dynamic in which toxicity is acquired via long somatic repeat expansions, then a therapy that slows somatic DNA-repeat expansion might prevent many human DNA-repeat disorders.

## Acknowledgments

We are grateful to the patients and families whose donations of brain tissue to the NIH NeuroBioBank enabled this work. We are grateful for support from the CHDI Foundation, the Harvard Medical School Department of Genetics, and the Harvard Ludwig Initiative in Neurodegenerative Disease. We are grateful to Tom Vogt, Vahri Beaumont, Jian Chen, and Cristina Sampaio for helpful advice and suggestions throughout this project’s conception and execution; to Darren Monckton, for helpful advice on amplifying and sequencing DNA repeats; to Sarah Tabrizi, Jim Gusella, and Marcy MacDonald, for helpful advice on HD; to John Warner, for helpful advice on computational modeling; and to Christina Usher, for contributions to the manuscript figures.

## Supplementary Materials

**Supplementary Figure 1**. Single-nucleus RNA-seq analysis of brain tissue from 56 persons with HD and 53 controls.

**Supplementary Figure 2**. SPN loss with HD progression (logarithmic scale).

**Supplementary Figure 3**. Decline in patch (striosomal) and matrix (extra-striosomal) SPNs with HD progression.

**Supplementary Figure 4**. HTT expression levels and rates of SPN loss.

**Supplementary Figure 5**. CAG-repeat length measurement from a set of sequence reads derived from individual HTT RNA transcripts.

**Supplementary Figure 6**. Allele specificity of somatic expansion.

**Supplementary Figure 7**. Alternative visualizations of the cell-type-specific CAG-repeat length data.

**Supplementary Figure 8**. Length distributions for the HD-causing CAG repeat in SPNs, in six persons with HD that were deeply sampled.

**Supplementary Figure 9**. Appearance of gene-expression changes in SPNs with the longest CAG-repeat expansions.

**Supplementary Figure 10**. Changes in SPN gene expression with somatic CAG-repeat expansion (volcano plots).

**Supplementary Figure 11**. Changes in SPN gene expression with somatic CAG-repeat expansion (p-value distributions).

**Supplementary Figure 12**. Similarity of long-repeat-expansion-associated gene-expression changes across persons with HD.

**Supplementary Figure 13**. Shared thresholds for gene-expression changes across SPN subtypes and individual persons with HD.

**Supplementary Figure 14**. Expression levels of six example phase C-genes in the individual SPNs of persons with HD.

**Supplementary Figure 15**. Expression levels of HTT in the individual SPNs of persons with HD.

**Supplementary Figure 16**. Relationship of de-repression (phase D) to CAG-repeat length and progression of phase-C gene-expression changes (additional visualization, deeply sampled donors).

**Supplementary Figure 17**. Relationship of de-repression (phase D) to CAG-repeat length and progression of phase-C gene-expression changes (additional visualization, case-control cohort). **Supplementary** Figure 18. Relationship of phase C-gene-expression changes to SPN loss.

**Supplementary Table 1**. Donor characteristics for deeply sampled donors

**Supplementary Table 2**. SPN loss estimates for deeply sampled donors.

**Supplementary Note 1**. Case-control gene-expression differences and their relationship to cell-autonomous and -non-autonomous effects of the CAG repeat in HD.

**Supplementary Note 2**. Somatic expansion in caudate cell types other than SPNs. **Supplementary Note 3**. Modeling the effect of CAG-repeat length on gene expression in SPNs. **Supplementary Note 4**. Modeling and simulation of SPN CAG-repeat expansion dynamics.

## Supplementary Figures

**Supplementary Figure 1.**
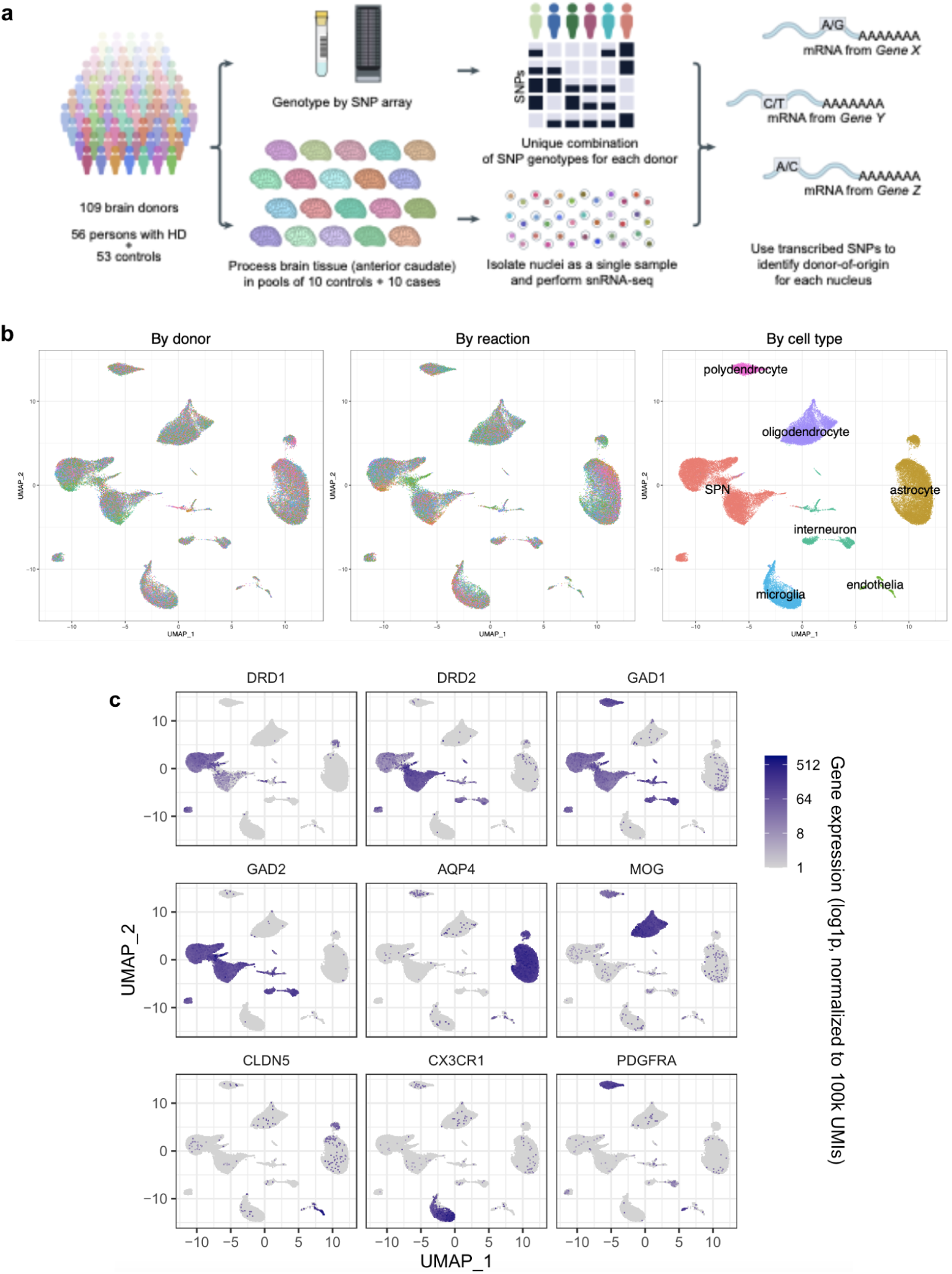
Single-nucleus RNA-seq analysis of brain tissue from 56 persons with HD and 53 controls. (**a**) “Cell village” workflow by which we perform snRNA-seq on sets of 20 donors at once. Image is only lightly modified from (Ling *et al*., 2024), where we describe this approach. (**b**) Multi-dimensional single cell RNA expression data for the 613 thousand striatal nuclei was projected into a two-dimensional space using the UMAP algorithm and then colored based on their donor-of-origin (left), village-of-origin (center), or assigned cell type (right), which was based on their genome-wide RNA-expression patterns. (**c**) The accuracy of these assignments can be visualized via the expression patterns of known cell-type-specific marker genes.

**Supplementary Figure 2.**
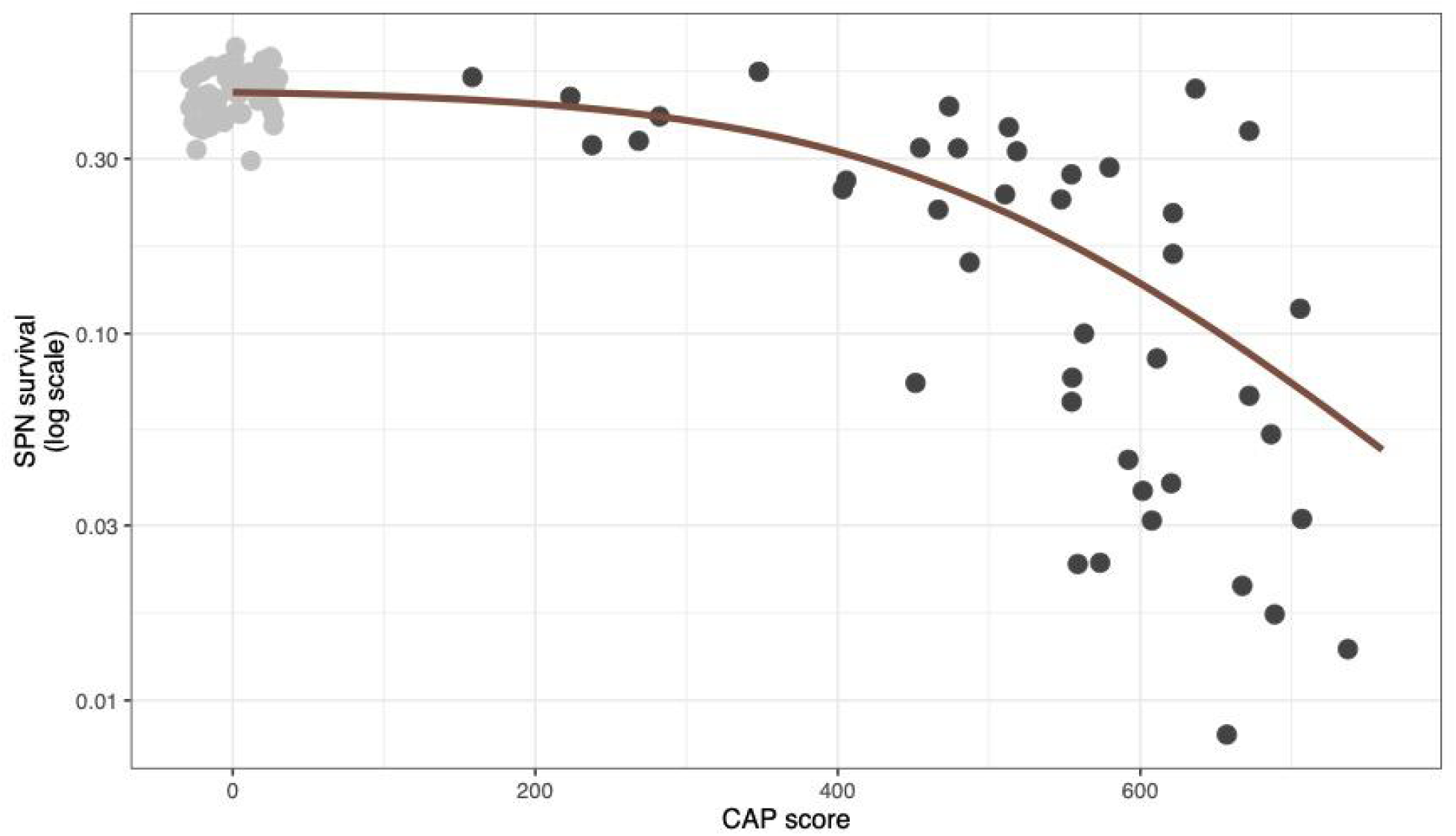
SPN loss with HD progression (logarithmic scale). Same data as in Fig. 1b, but shown here on a logarithmic scale, in which the slope of the relationship estimates rates of SPN loss with increasing CAP score. Unaffected control donors are shown as gray circles with CAP score of zero (jitter has been added to reduce overplotting). The red curve is from a fit of a logistic function to the SPN survival curve (before log-transformation, i.e. as in Fig. 1a).

**Supplementary Figure 3.**
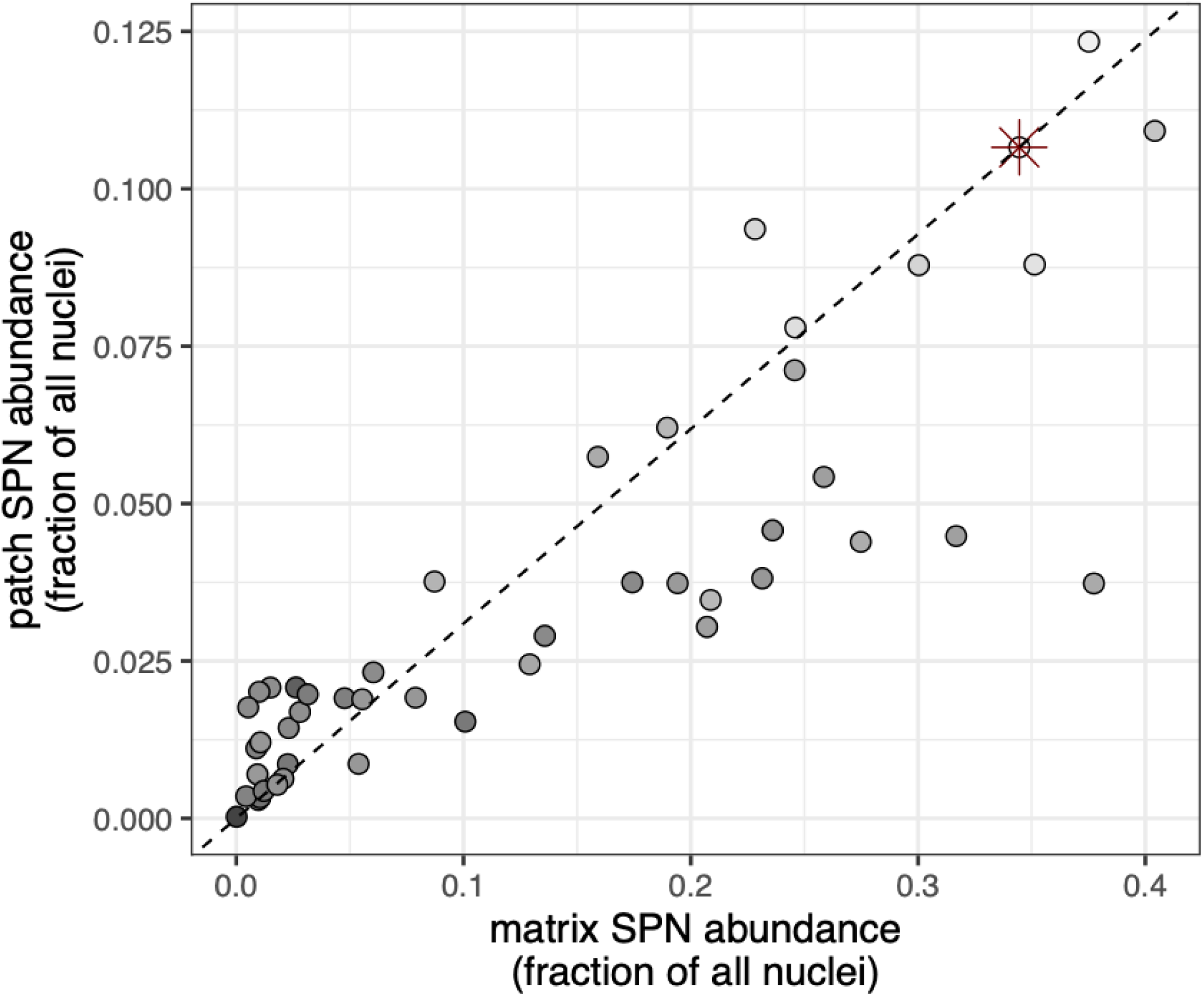
Decline in patch (striosomal) and matrix (extra-striosomal) SPNs with HD progression. Gray scale represents CAP score as in Fig. 1. Red asterisk denotes the median of 56 unaffected control donors. Gray scale represents CAP score as in Fig. 1.

**Supplementary Figure 4.**
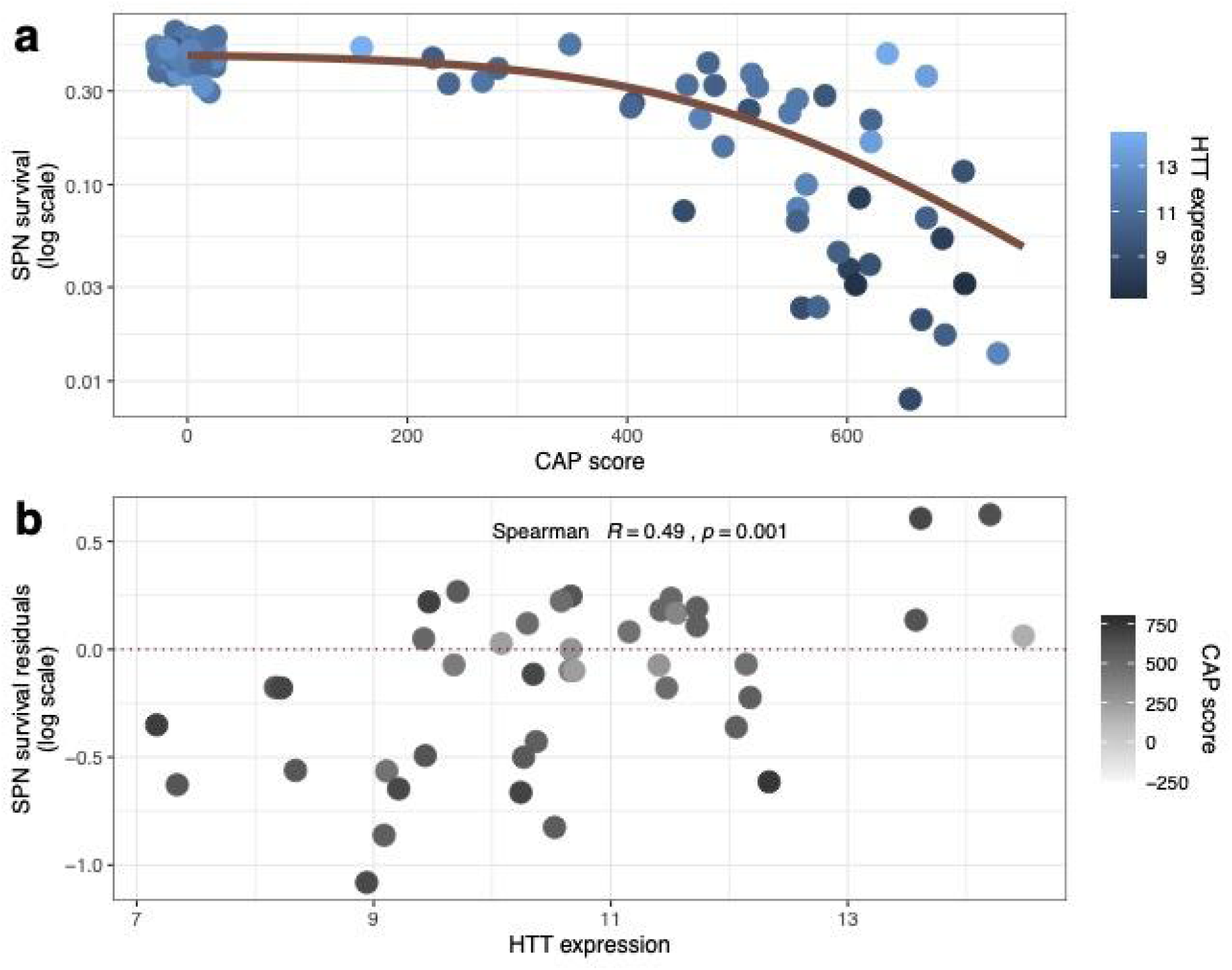
*HTT* expression levels and rates of SPN loss. (**a**) SPN survival rates (on a logarithmic scale, y-axis) are plotted against CAP score, an estimate of age-expected HD progression. The red curve is from a fit of a logistic function to the SPN survival curve. Points are colored to reflect individual donors’ expression levels of *HTT* in SPNs. (**b**) Residuals of the relationship in **a** are plotted against individual donors’ *HTT* expression levels. Gray shading represents CAP score. Note that individuals’ *HTT* expression levels actually show a modest positive relationship to SPN survival (rather than the negative relationship predicted by the “cumulative lifetime damage” model, in which individuals with higher expression levels would exhibit earlier/faster SPN loss), though this arises substantially from the donors in the lower-left part of the plot – donors with very high CAP scores and extreme caudate atrophy.

**Supplementary Figure 5.**
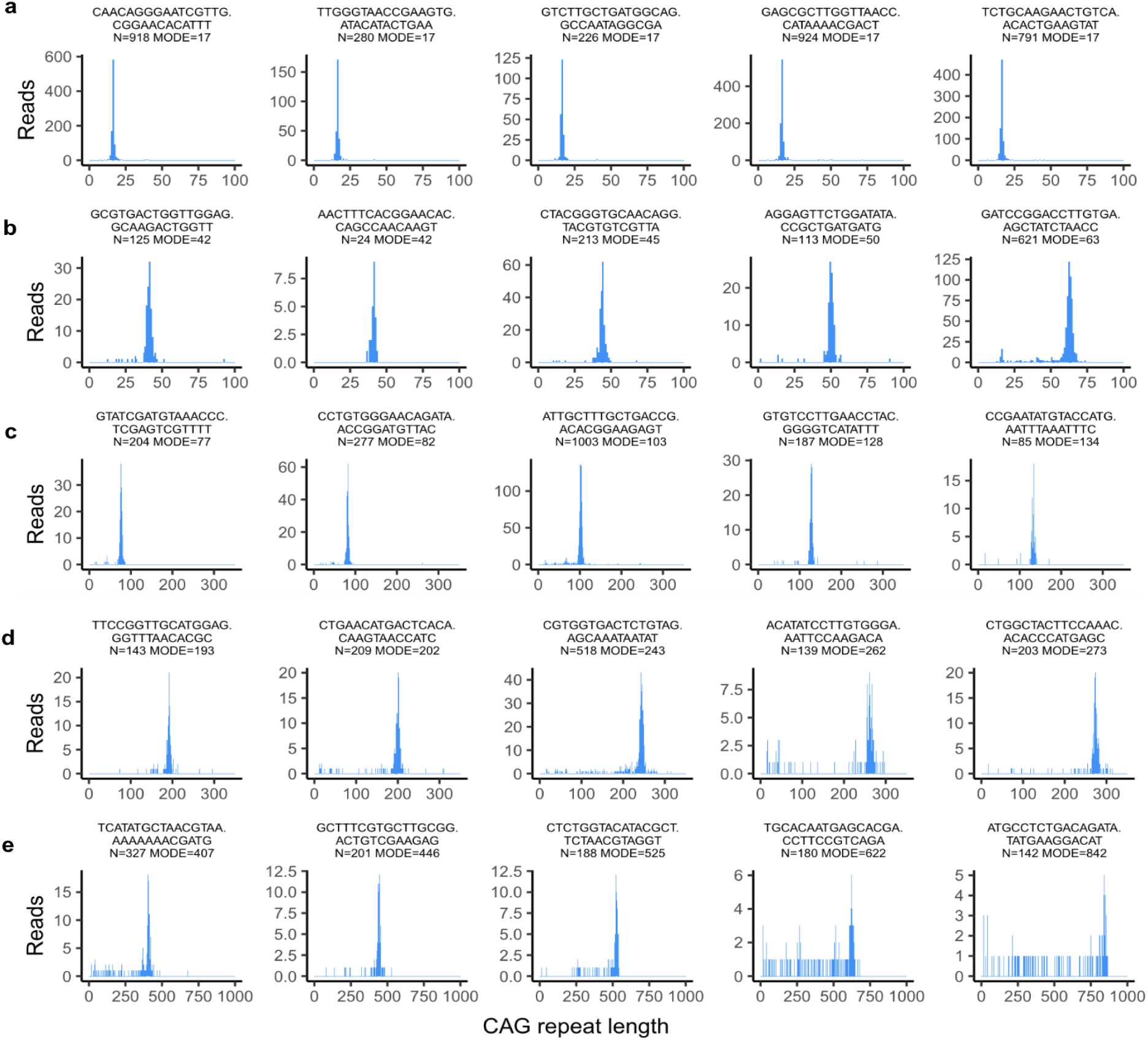
CAG-repeat length measurement from a set of sequence reads derived from individual *HTT* RNA transcripts. Sequence reads (PCR products) originating from the same underlying RNA molecule share a common unique molecular identifier (UMI) that we had applied during reverse transcription. The histograms show representative distributions of CAG-repeat lengths in the reads for individual UMIs (**a**) for short, common HD alleles (<35 repeats) that do not cause HD; (**b**) for HD-causing alleles with modest somatic expansion in the range of 40-60 CAGs (**c**) for longer somatic expansions in the range of 80-150 CAGs (**d**) for UMIs with somatic expansions in the range of 150-300 CAGs and (**e**) for UMIs showing very long somatic expansions beyond 300 CAGs. Note that each row uses a different x-axis scale. For transcripts with long somatically acquired CAG-repeat expansions, the PCR amplification during library preparation creates a left-tailed distribution, reflecting the way that PCR errors lead to the generation of molecules with shorter repeats and then favor these smaller molecules over longer ones. For each UMI, we use the mode (the Robertson-Cryer half-sample mode estimator, function hsm() from the R package modeest) as the consensus CAG-repeat length for that *HTT* transcript. The accuracy of these determinations is evaluated in Fig. 2b. Cell barcodes on the same sequence reads make it possible to then connect each such CAG-repeat length determination to the wider RNA-expression profile (cell type and cell state) of the nucleus from which it was sampled.

**Supplementary Figure 6.**
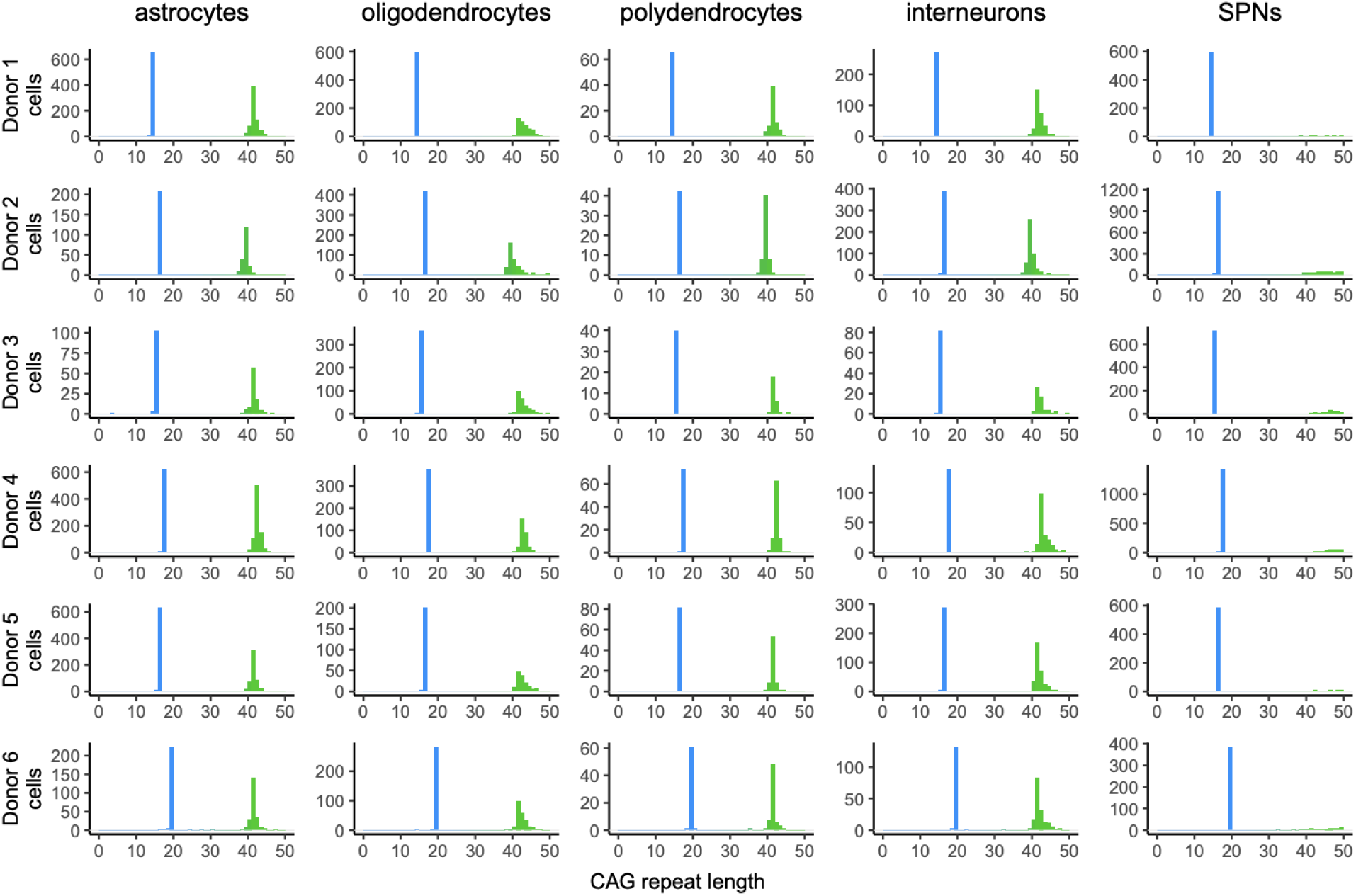
Allele specificity of somatic expansion. Distributions of CAG-repeat length showing both the short, non-HD-causing allele (< 35 repeats, blue) and the longer, HD-causing allele (>35 repeats, green), here zoomed in to the 0-50 range (beyond which most SPNs have already expanded their HD-causing allele). The shorter allele (blue) appears to be somatically stable across neuronal and glial cell types in all six persons with HD.

**Supplementary Figure 7.**
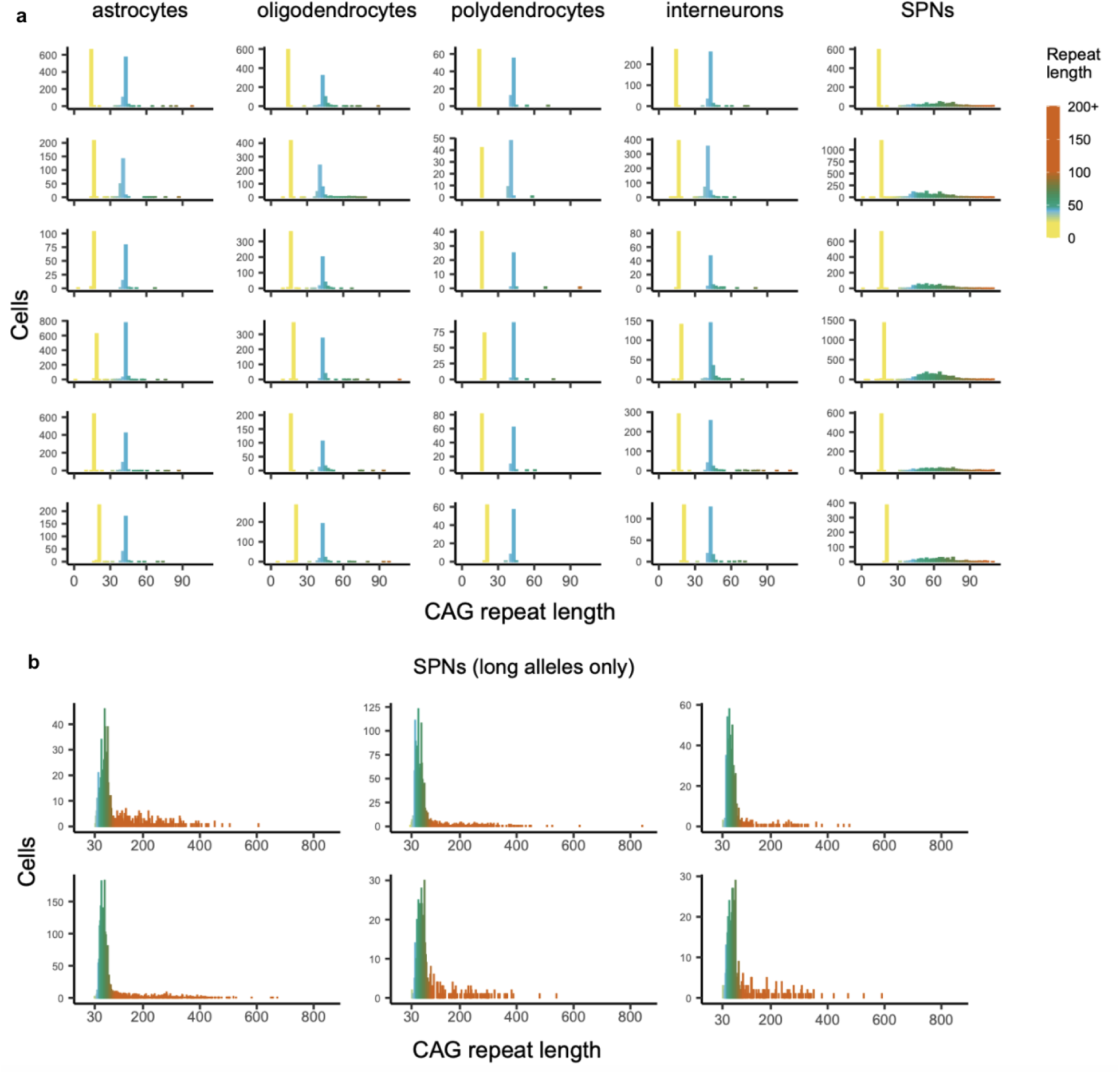
Alternative visualizations of the cell-type-specific CAG-repeat length data from Fig. 2c**,d**. Color scale is the same in both panels above.

**Supplementary Figure 8.**
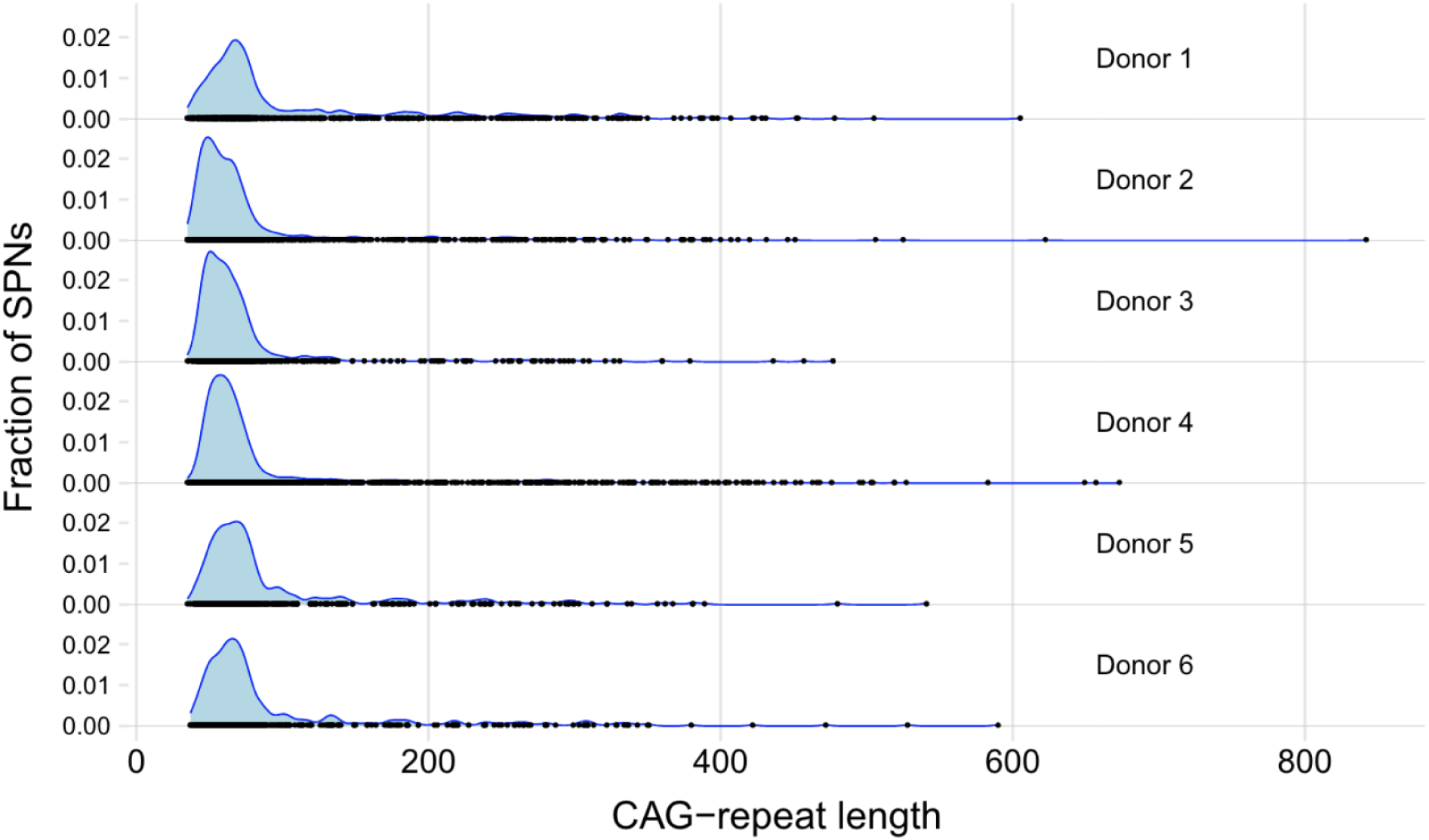
Length distributions for the HD-causing CAG repeat in SPNs, in six persons with HD that were deeply sampled (rows). Blue shaded areas are smoothed density estimates of the repeat-length distribution. Overplotted black points are a rug showing the measurements in individual SPNs. In all six donors, the repeat-length distribution exhibits an “armadillo-like” shape, where the majority of each donor’s SPNs have undergone modest expansion, up to 100 CAGs, but a small fraction of SPNs have undergone greater expansion with some reaching over 500 CAGs.

**Supplementary Figure 9.**
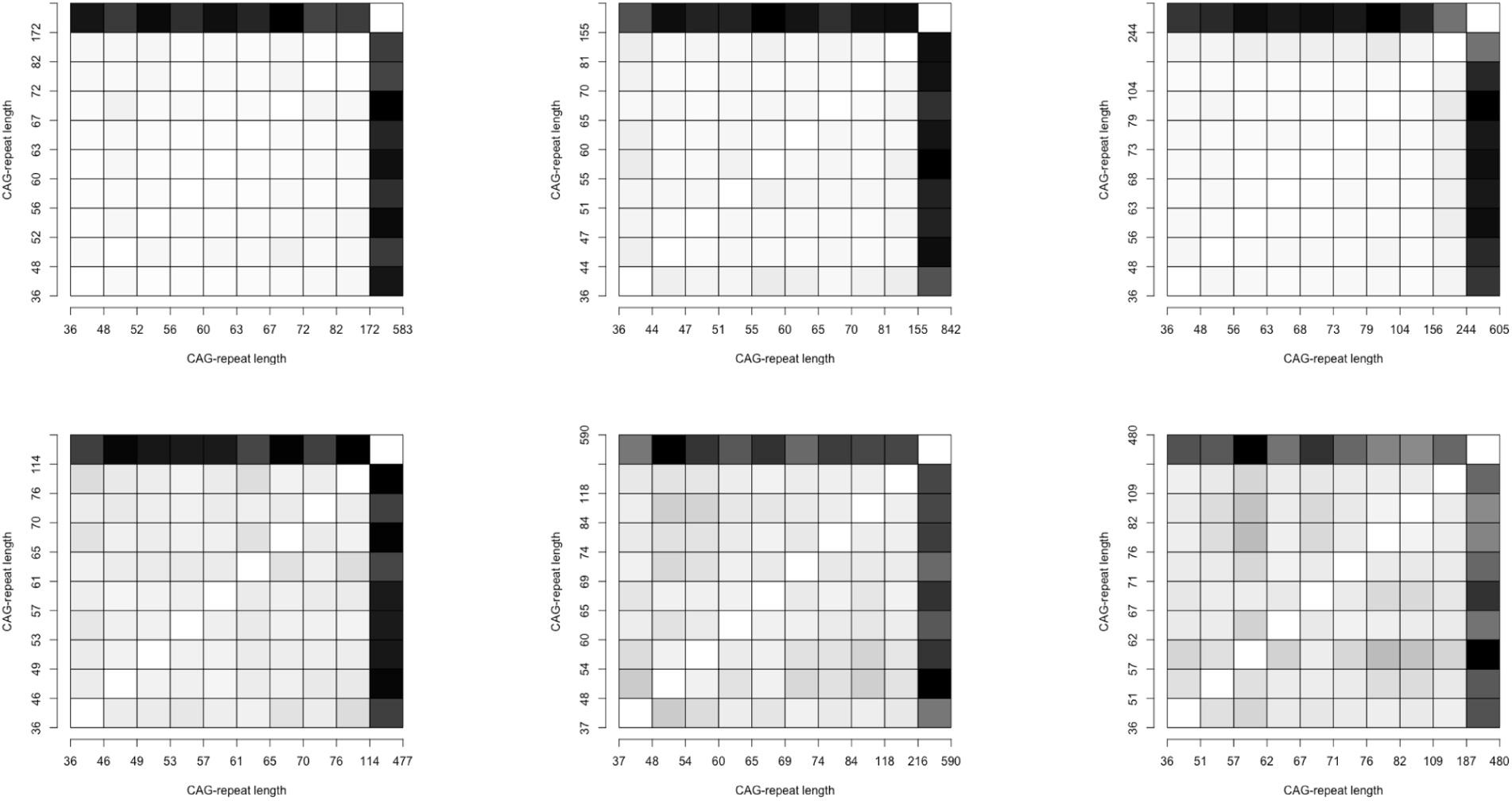
Appearance of gene-expression changes in SPNs with the longest CAG-repeat expansions. Correlation-squares show the magnitude of gene-expression differences (one minus the correlation coefficient) when comparing sets of SPNs (from the same donor) that have been grouped into deciles based on the CAG-repeat length of their HD-causing *HTT* allele. Gray scale: black indicates maximal difference observed in any comparison; white indicates no difference. Note that the CAG-repeat length thresholds for the deciles vary by donor, and that data for donors whose SPNs were less deeply sampled (e.g. lower right) exhibit more statistical noise. The donor in the upper left is the same donor shown in Fig. 3. Depth of sampling (row 1: 2337 SPNs; 1489 SPNs; 738 SPNs; row 2: 767 SPNs; 468 SPNs; 467 SPNs).

**Supplementary Figure 10.**
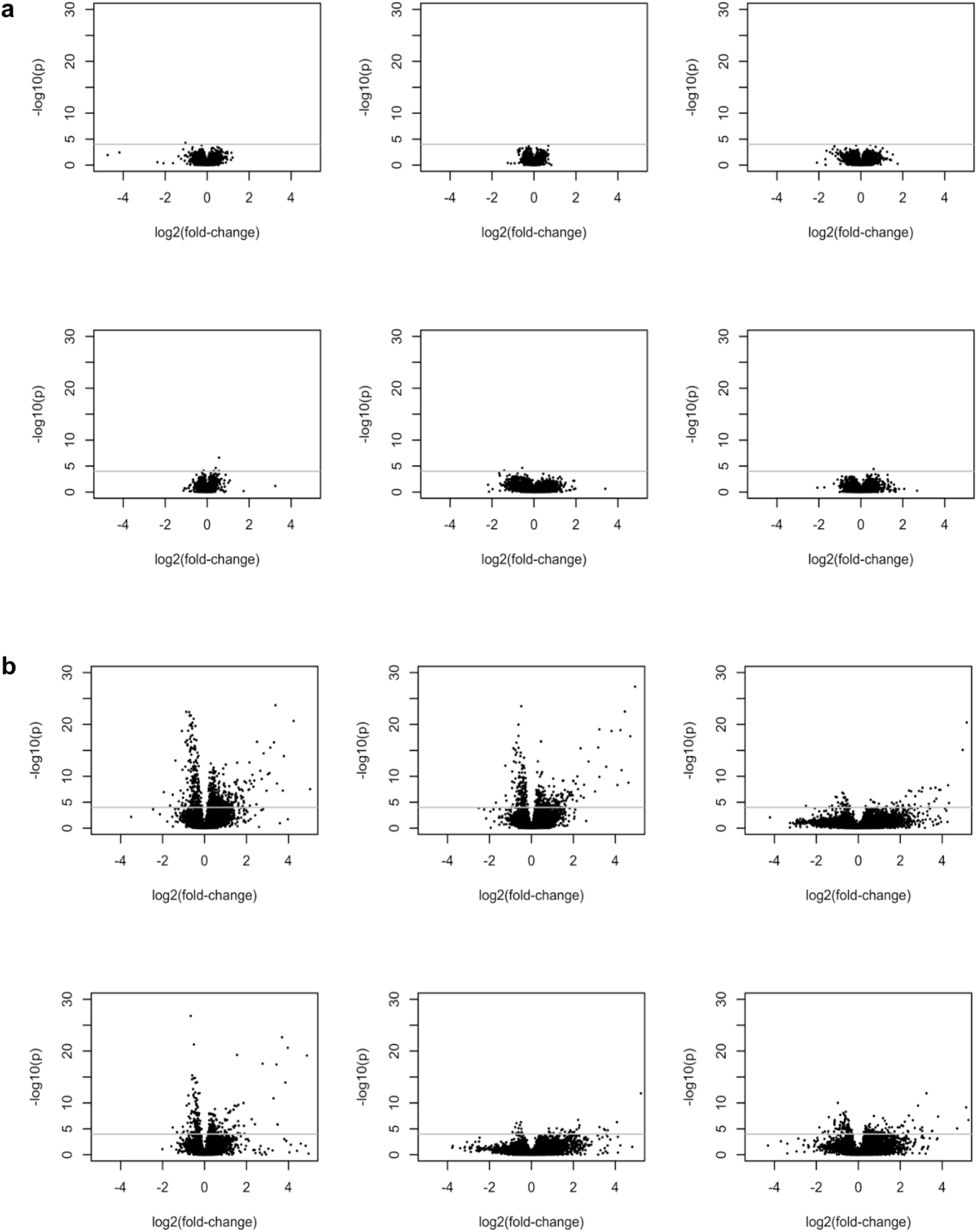
Changes in SPN gene expression with somatic CAG-repeat expansion (volcano plots). (**a**) Comparisons of gene expression (volcano plots) of SPNs with 35-65 to SPNs with 66-150 CAGs, in six persons with HD. Dashed lines show the thresholds for genome-wide significance. (**b**) Comparisons of gene expression (volcano plots) of SPNs with 35-150 to SPNs with>150 CAGs, in six persons with HD. Note that the statistical power of the analysis varies by donor in relation to the depth of sampling of SPNs with long (>150 CAGs) repeat expansion.

**Supplementary Figure 11.**
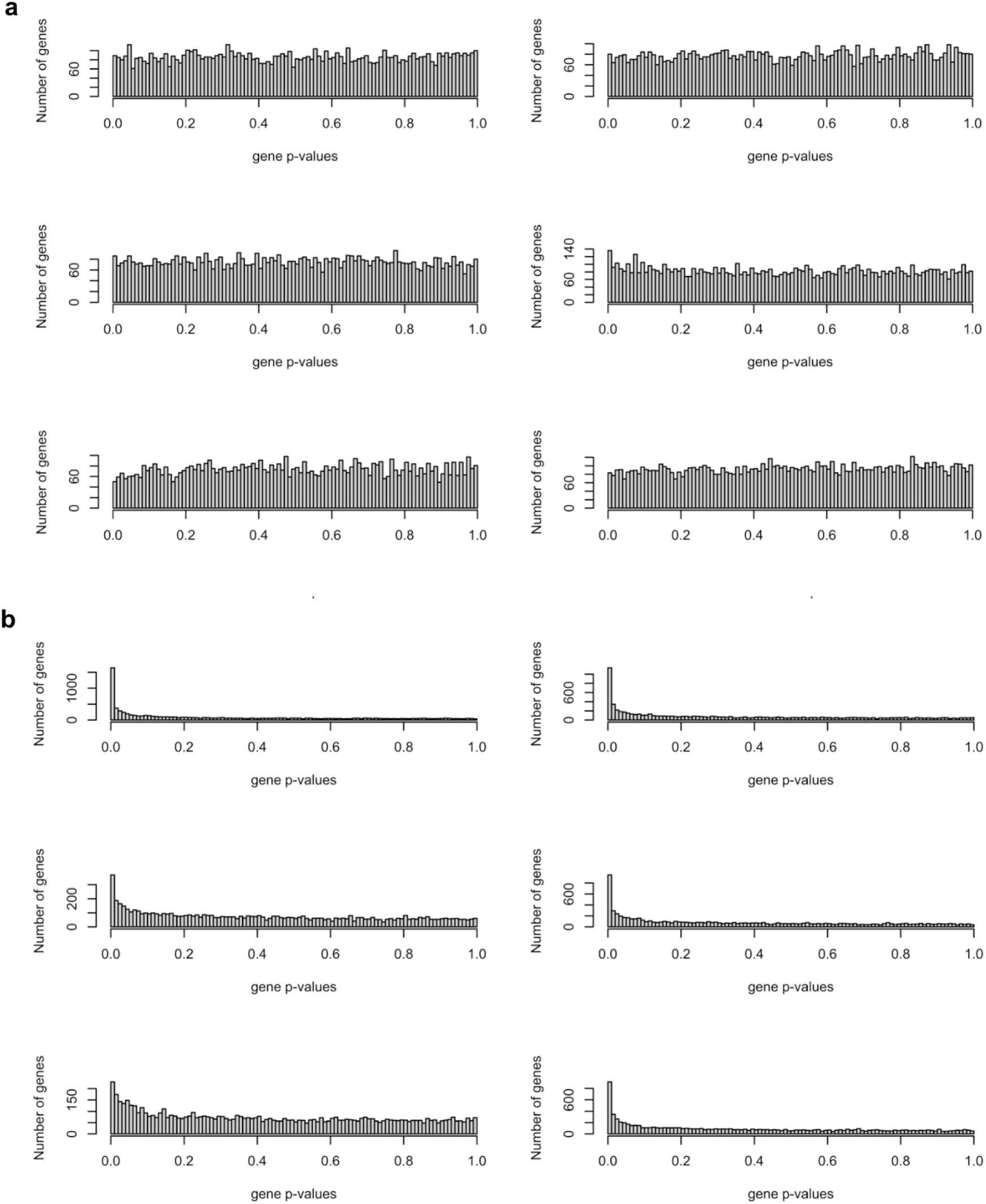
Changes in SPN gene expression with somatic CAG-repeat expansion (p-value distributions). (**a**) Comparisons of gene expression (p-value distributions) of SPNs with 35-65 to SPNs with 66-150 CAGs, in six persons with HD. (**b**) Comparisons of gene expression (p-value distributions) of SPNs with 35-150 to SPNs with>150 CAGs, in six persons with HD. Same donors as in Supplementary Fig. 2. Note that the statistical power of the analysis varies by donor in relation to the depth of sampling of SPNs with long (>150 CAGs) repeat expansion.

**Supplementary Figure 12.**
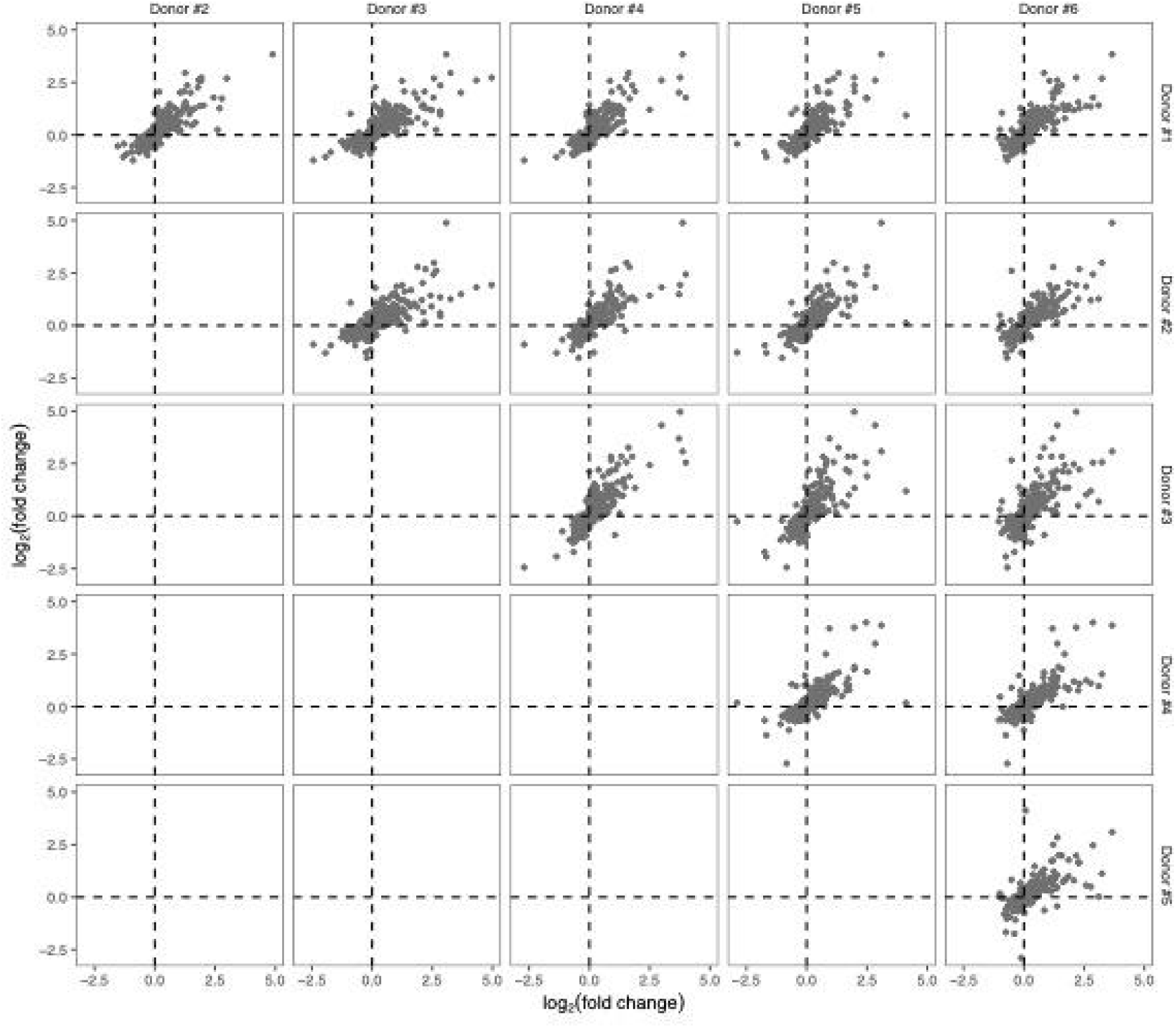
Similarity of long-repeat-expansion-associated gene-expression changes across persons with HD. Each panel is a pairwise comparison of SPN gene-expression data involving two persons with HD (x- and y-axes), in which the values on the two axes are plotted are the log2-fold-changes in gene expression when comparing (within-tissue) SPNs with >150 CAGs to SPNs with <150 CAGs. Genes whose expression levels change significantly with repeat expansion in at least one of the donors are shown.

**Supplementary Figure 13.**
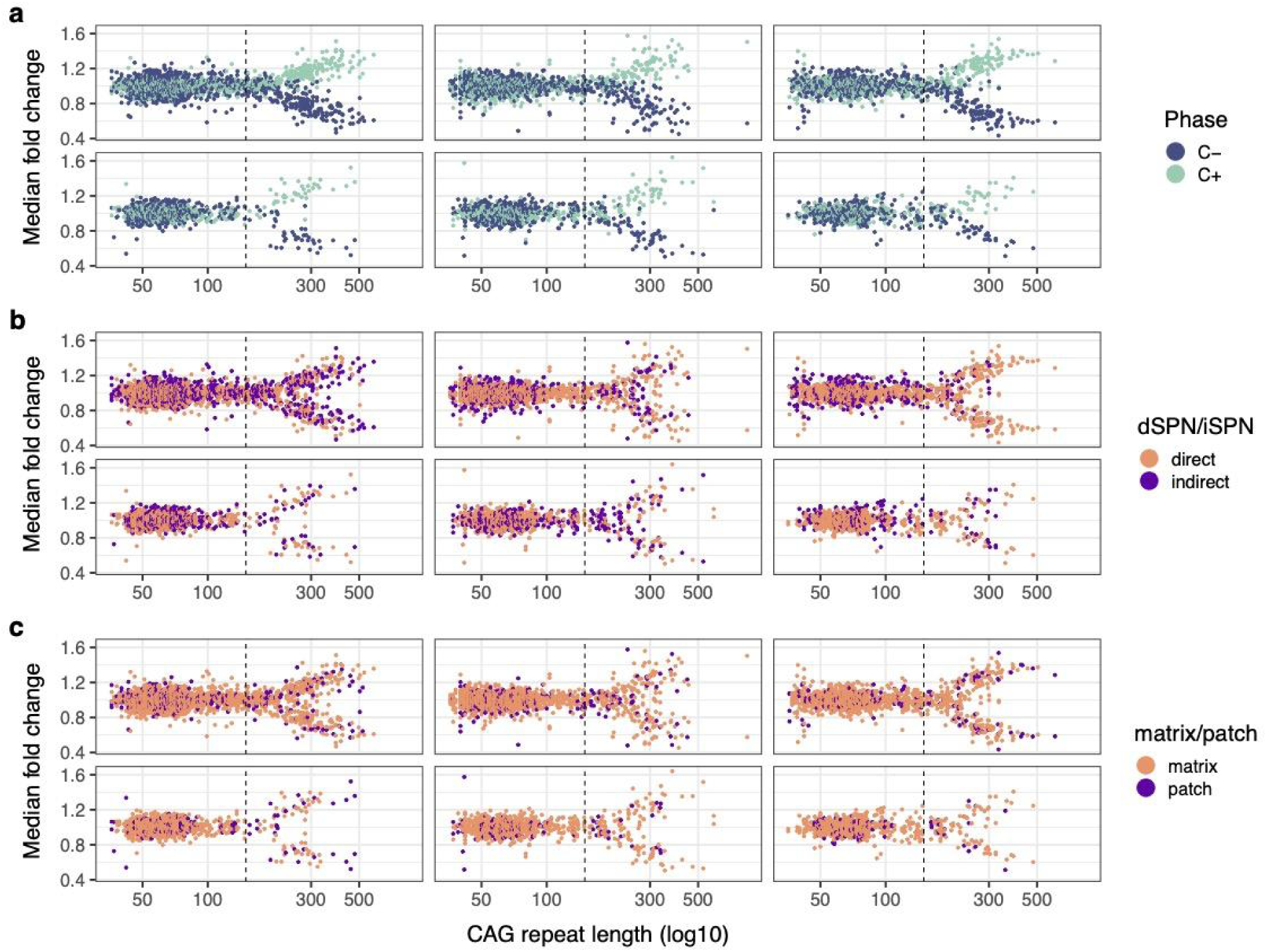
Shared thresholds for gene-expression changes across SPN subtypes and individual persons with HD. (**a**) Relationship between transcriptional changes and CAG-repeat length (same data as in Fig. 4b, but here with CAG-repeat length on the x-axis, using a logarithmic scale). Each SPN is represented by both a blue point and a green point: blue points show the median fold-change of a set of 203 genes that decrease in expression with repeat expansion (C-genes); green points show the average fold-change of a set of 369 genes that increase in expression with repeat expansion (C+ genes). (**b,c**) Same data in **a,** but here the same points are colored to show the subtype of SPN. Each SPN is represented by two points (here, in the same color) that show the expression fold-change for its C+ and C-genes. In b, “direct” and “indirect” refer to dSPNs (D1 SPNs) and iSPNs (D2 SPNs); in c, “patch” and “matrix” refer to striosomal SPNs and extra-striosomal SPNs respectively.

**Supplementary Figure 14.**
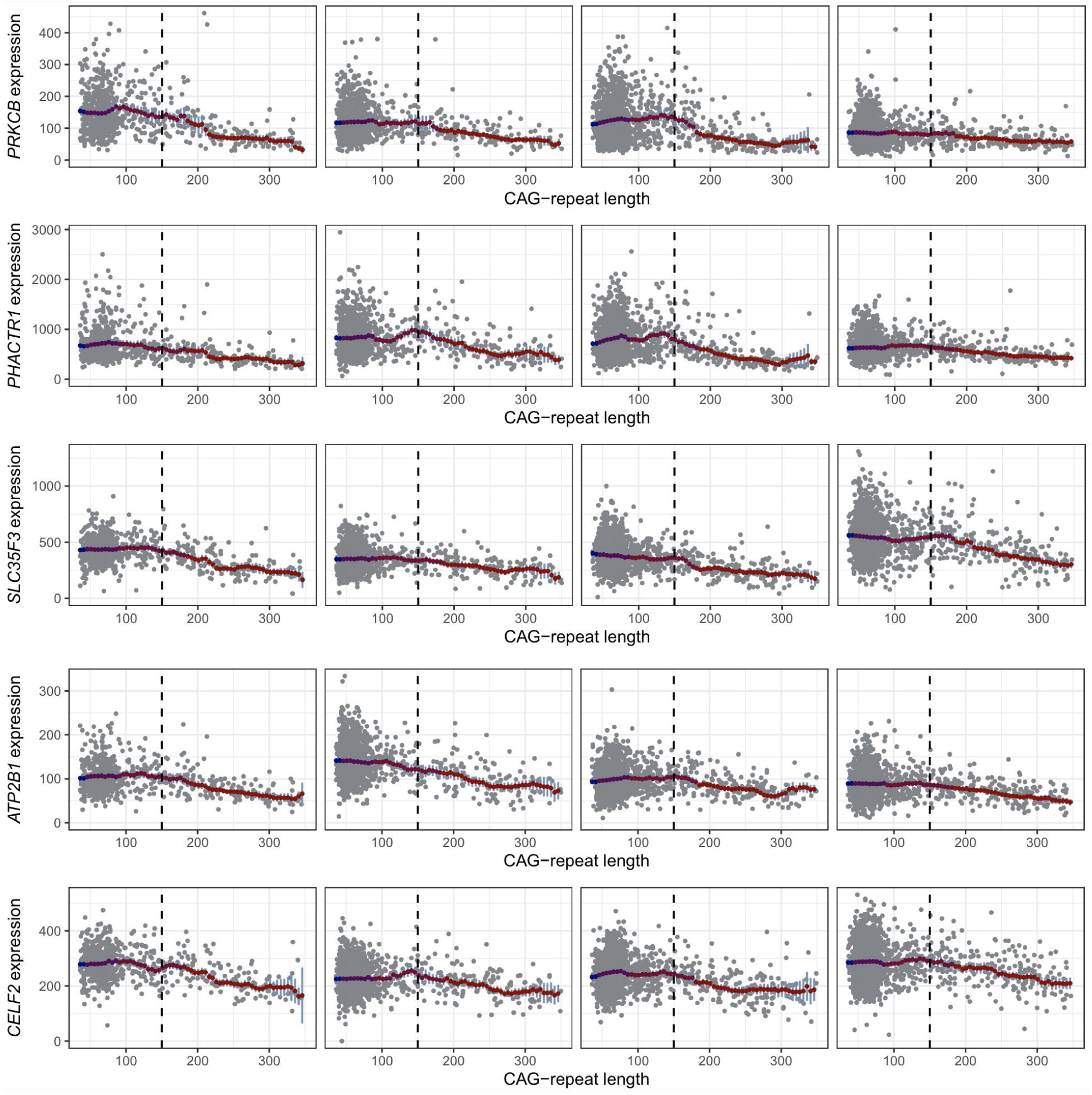
Expression levels of six example phase C-genes in the individual SPNs of four persons with HD. One gene per row, one donor per column. Gray points represent individual SPNs. Red bars and confidence intervals (blue) average the data in moving windows of width log_2_(CAG-repeat length) to reduce the measurement noise inherent in single-cell measurements. The genes shown are *PRKCB, PHACTR1, SLC35F3, ATP2B1*, and *CELF2*. All are genes that SPNs normally express more strongly than interneurons do. This analysis also helps explain why conventional descriptive-genomics analyses (to find “differentially expressed genes” in case-control comparisons) generally fail to recognize such effects in HD. First, these effects are present in just a small fraction of any donor’s SPNs at any one time (those SPNs with long CAG-repeat expansions), and are pronounced in a still-smaller fraction (those in which the CAG repeat has expanded even further) – so they appear as small and often insignificant changes in bulk and sorted-cell-type analyses. Second, most of these genes, like most human genes, exhibit normal inter-individual variation in expression levels (at baseline), further obscuring (in case-control comparisons) effects that are clear in within-donor, within-tissue comparisons of individual cells.

**Supplementary Figure 15.**
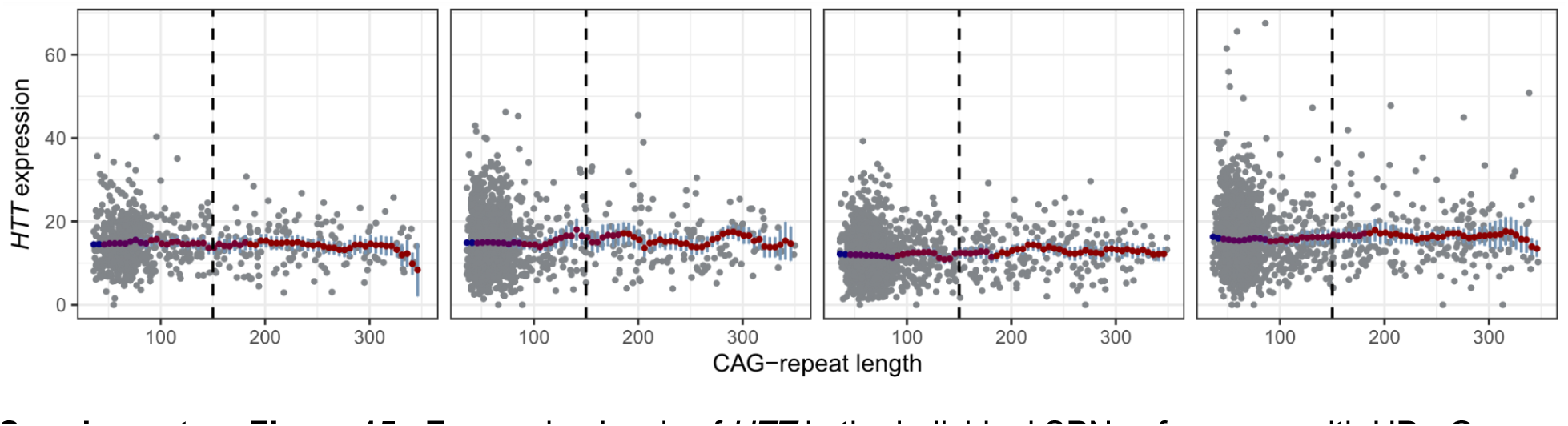
Expression levels of *HTT* in the individual SPNs of persons with HD. Gray points represent individual SPNs. Red bars and confidence intervals (blue) average the data in sliding windows to reduce the measurement noise inherent in single-cell measurements.

**Supplementary Figure 16.**
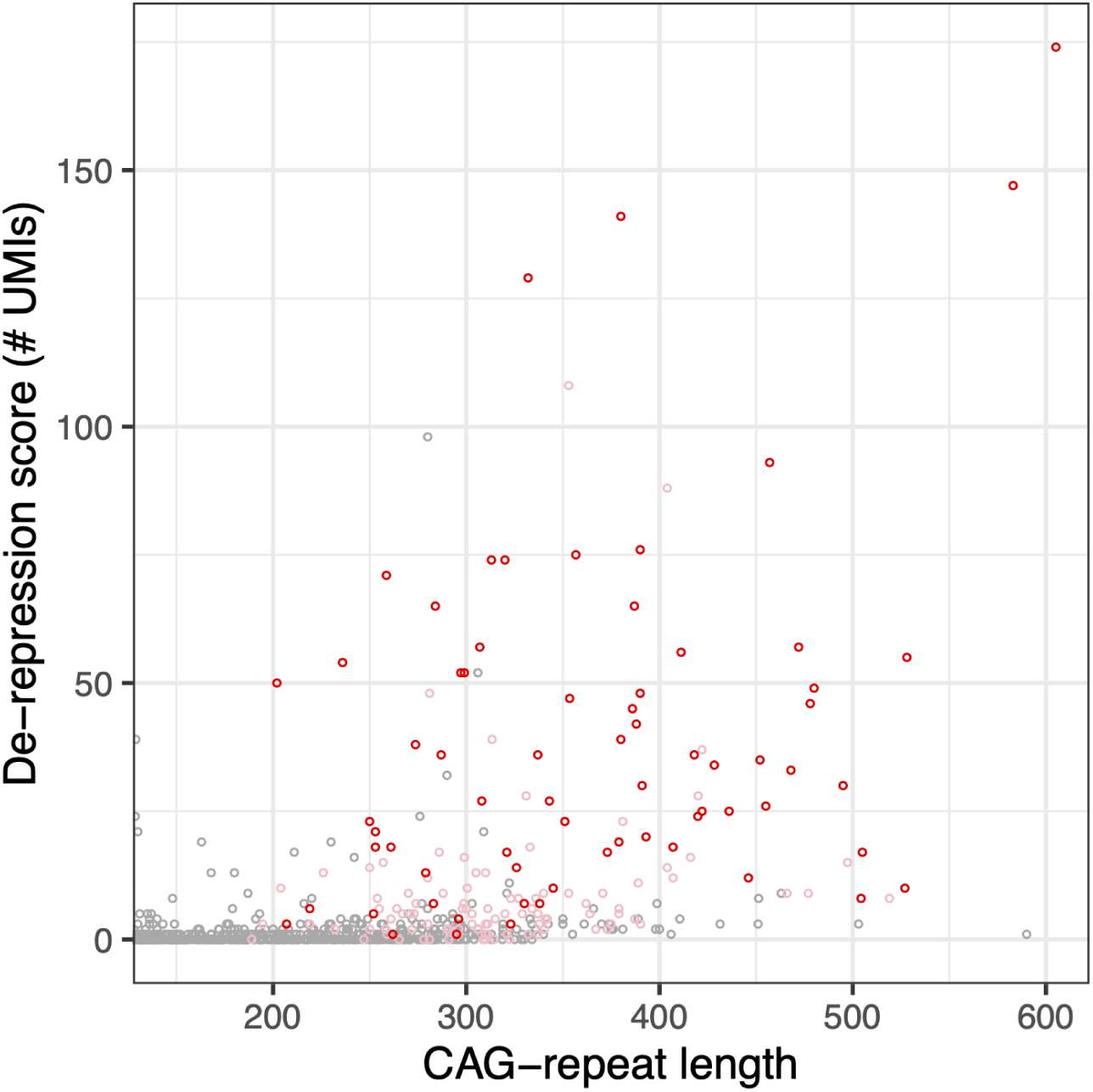
Relationship of de-repression (y-axis) to CAG-repeat length (x-axis) and progression of phase-C gene-expression changes (point colors). This is an alternative visualization of the relationship in Fig. 5a, to make more visible the CAG-repeat lengths of most of the SPNs in which these genes have become de-repressed.

**Supplementary Figure 17.**
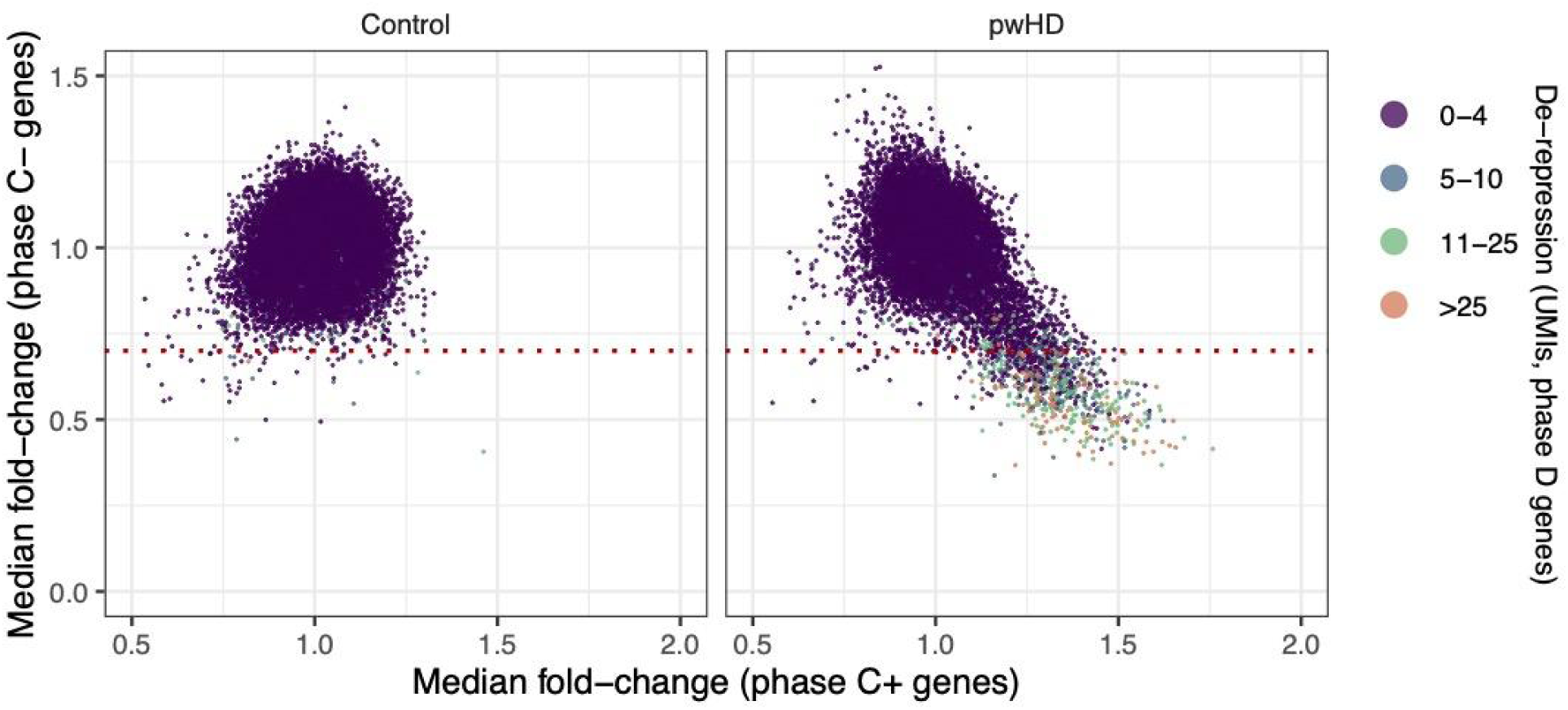
Relationships (across individual SPNs, points) among phase-D de-repression (indicated by point color, see figure legend) and progression of phase-C gene-expression changes (x-axis: phase C+ (increasing-expression) genes; y-axis: phase C- (decreasing-expression) genes). The left panel shows SPNs sampled from 56 unaffected control donors; the right panel shows SPNs sampled from 53 persons with HD. Dotted line shows the threshold used to define “altered transcription” for the analysis in **FIg. 5e**.

**Supplementary Figure 18.**
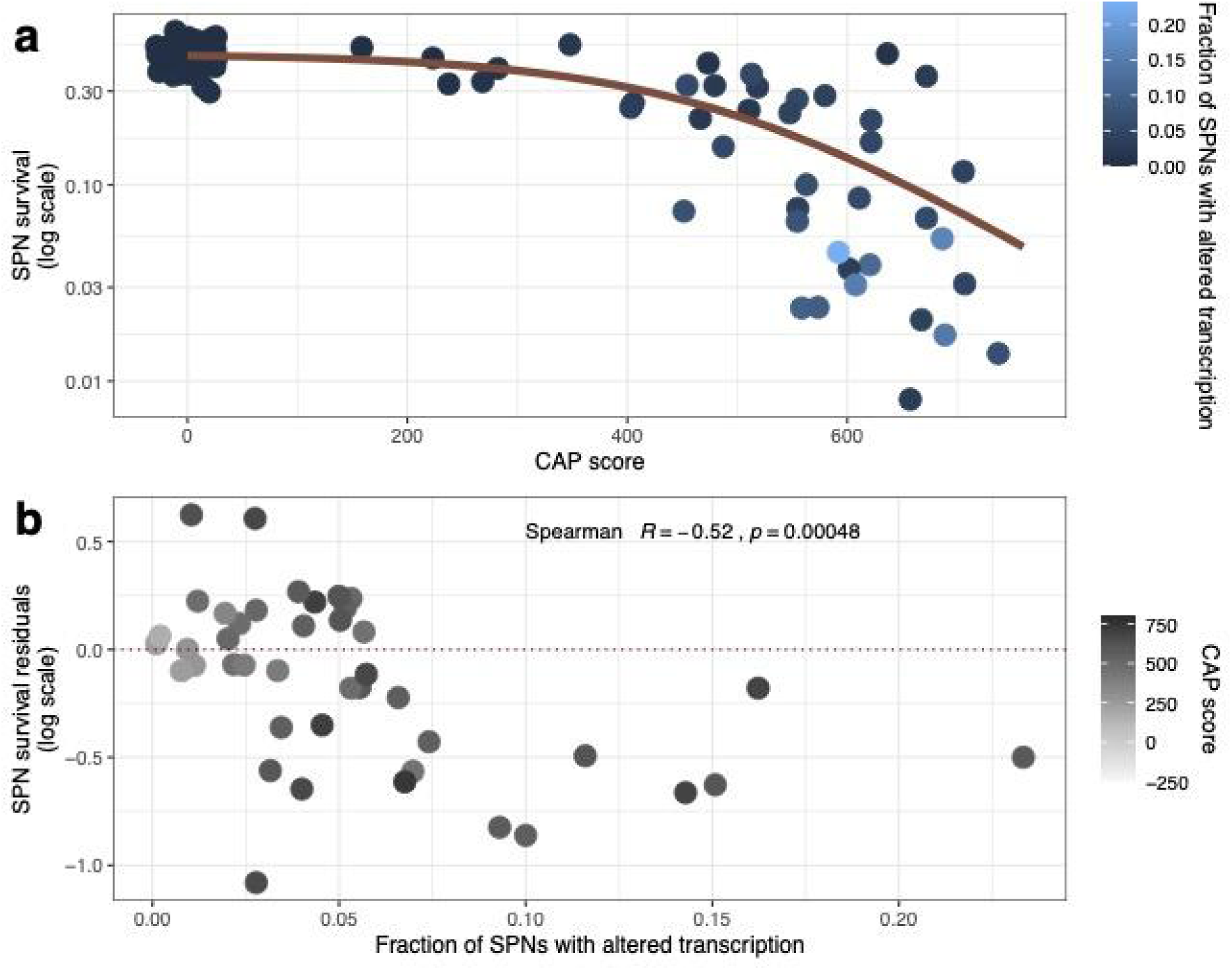
CAG-repeat-driven transcriptionopathy and rates of SPN loss. (**a**) SPN survival (on a logarithmic scale, y-axis) is plotted against CAP score, an estimate of age-expected HD progression. The red curve is from a fit of a logistic function to the SPN survival curve. Points are colored based on the fraction of each donor’s SPNs that have phase-C transcriptionopathy (which is plotted on the x-axis in **b**). (**b**) Residuals of the relationship in **a** (negative values represent excess SPN loss relative to age-expected loss) are plotted (y-axis) against the fraction of SPNs with transcriptionopathy (x-axis). Gray shading represents CAP score. Note in both panels that persons with excess SPN loss (relative to CAP score) tend also to be those for whom a larger fraction of SPNs exhibit phase-C transcriptionopathy.

**Supplementary Table 1.**
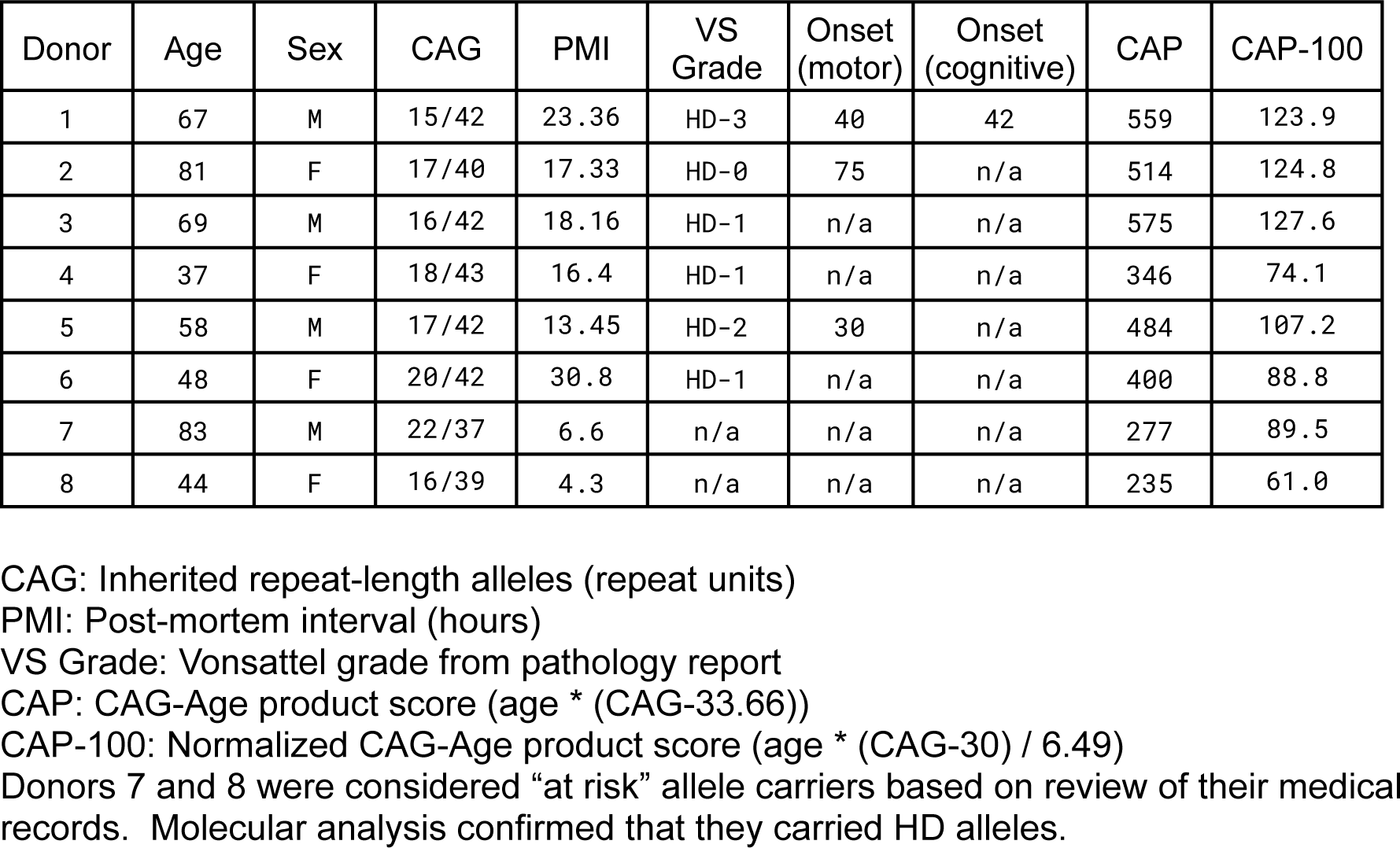
Donor characteristics for deeply sampled donors

**Supplementary Table 2.**
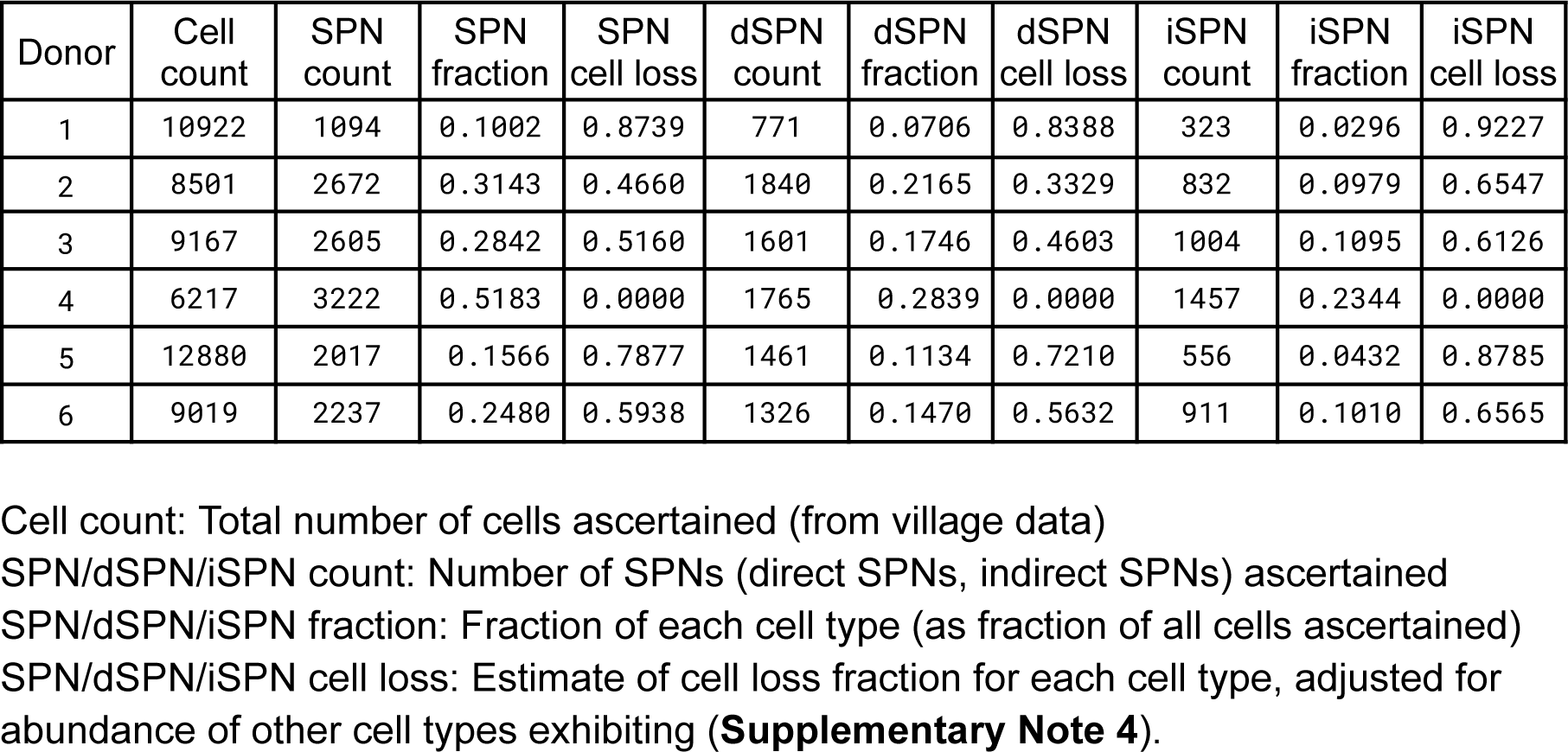
SPN loss estimates for deeply sampled donors

| Donor | Cell count | SPN count | SPN fraction | SPN cell loss | dSPN count | dSPN fraction | dSPN cell loss | iSPN count | iSPN fraction | iSPN cell loss |
| --- | --- | --- | --- | --- | --- | --- | --- | --- | --- | --- |
| 1 | 10922 | 1094 | 0.1002 | 0.8739 | 771 | 0.0706 | 0.8388 | 323 | 0.0296 | 0.9227 |
| 2 | 8501 | 2672 | 0.3143 | 0.4660 | 1840 | 0.2165 | 0.3329 | 832 | 0.0979 | 0.6547 |
| 3 | 9167 | 2605 | 0.2842 | 0.5160 | 1601 | 0.1746 | 0.4603 | 1004 | 0.1095 | 0.6126 |
| 4 | 6217 | 3222 | 0.5183 | 0.0000 | 1765 | 0.2839 | 0.0000 | 1457 | 0.2344 | 0.0000 |
| 5 | 12880 | 2017 | 0.1566 | 0.7877 | 1461 | 0.1134 | 0.7210 | 556 | 0.0432 | 0.8785 |
| 6 | 9019 | 2237 | 0.2480 | 0.5938 | 1326 | 0.1470 | 0.5632 | 911 | 0.1010 | 0.6565 |

Cell count: Total number of cells ascertained (from village data)
SPN/dSPN/iSPN count: Number of SPNs (direct SPNs, indirect SPNs) ascertained
SPN/dSPN/iSPN fraction: Fraction of each cell type (as fraction of all cells ascertained)
SPN/dSPN/iSPN cell loss: Estimate of cell loss fraction for each cell type, adjusted for abundance of other cell types exhibiting (**Supplementary Note 4**).

## Supplementary Note 1: Case-control gene-expression differences and their relationship to cell-autonomous and -non-autonomous effects of the CAG repeat in HD

A conventional approach to functional genomics in human disease involves comparing gene-expression data between cases and controls to arrive at a list of “differentially expressed genes” (DEGs). We found that, even when we applied a conservative statistical approach (a non-parametric Wilcox test comparing the 53 cases to the 56 controls) to identifying differentially expressed genes, every caudate cell type – including all types of neurons, glia, and vascular cells – exhibited thousands of DEGs whose expression levels differed (on average) between cases and controls (**Fig. SN1.1**). This broadly altered gene expression in every cell type potentially reflects the profound consequences of HD, which causes atrophy of the entire caudate (reduced in HD to a small fraction of its normal size), neuronal death, and devascularization. Such changes may affect the biology of every cell type.

**Figure SN1.1.**
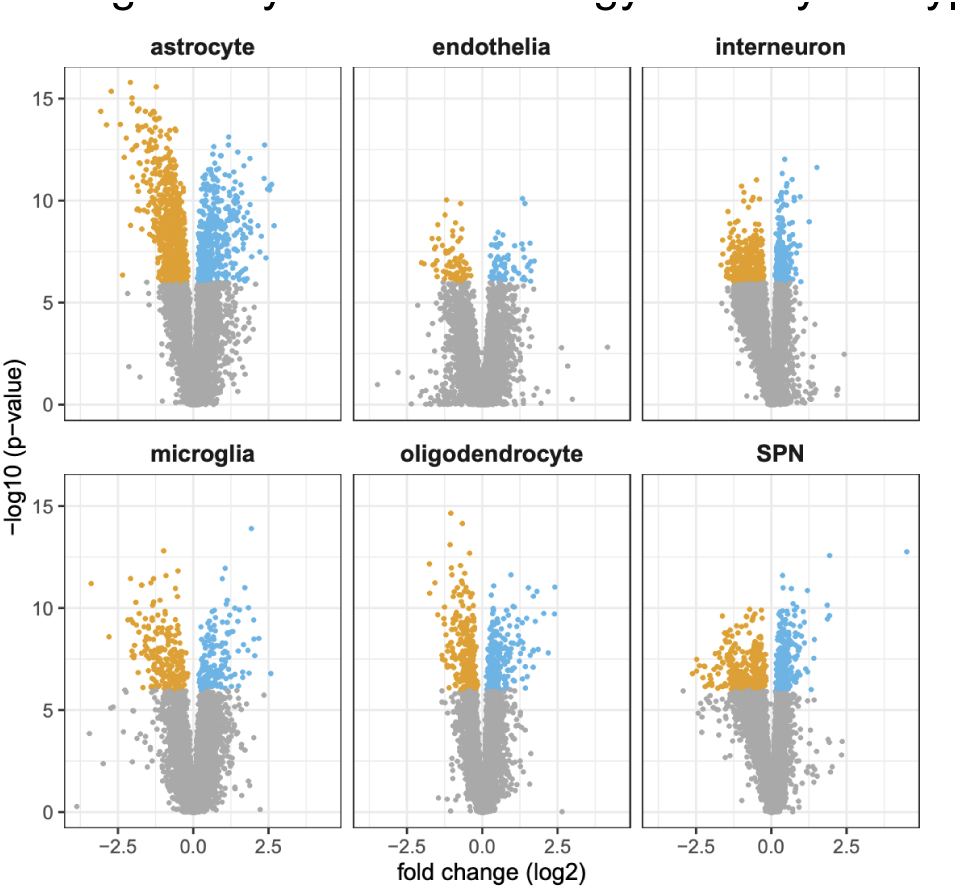
Case-control gene-expression differences. Volcano plots representing case-control differences in gene expression in each cell type. Median fold-changes (x-axis) and p-values (y-axis) are based on comparison of 53 persons with HD to 56 controls, using a Wilcox (non-parametric) test. Blue points: genes with higher average expression levels in HD; orange points: genes with reduced average expression levels in HD.

The expression levels of these same genes (case-control DEGs) correlated strongly, in an HD-only analysis, with donors’ earlier caudate atrophy (**Fig. SN1.2**) as measured by SPN survival (estimated as in Fig. 2). This was true of SPN gene expression as well as gene expression in other cell types (**Fig. SN1.2**).

**FIgure SN1.2.**
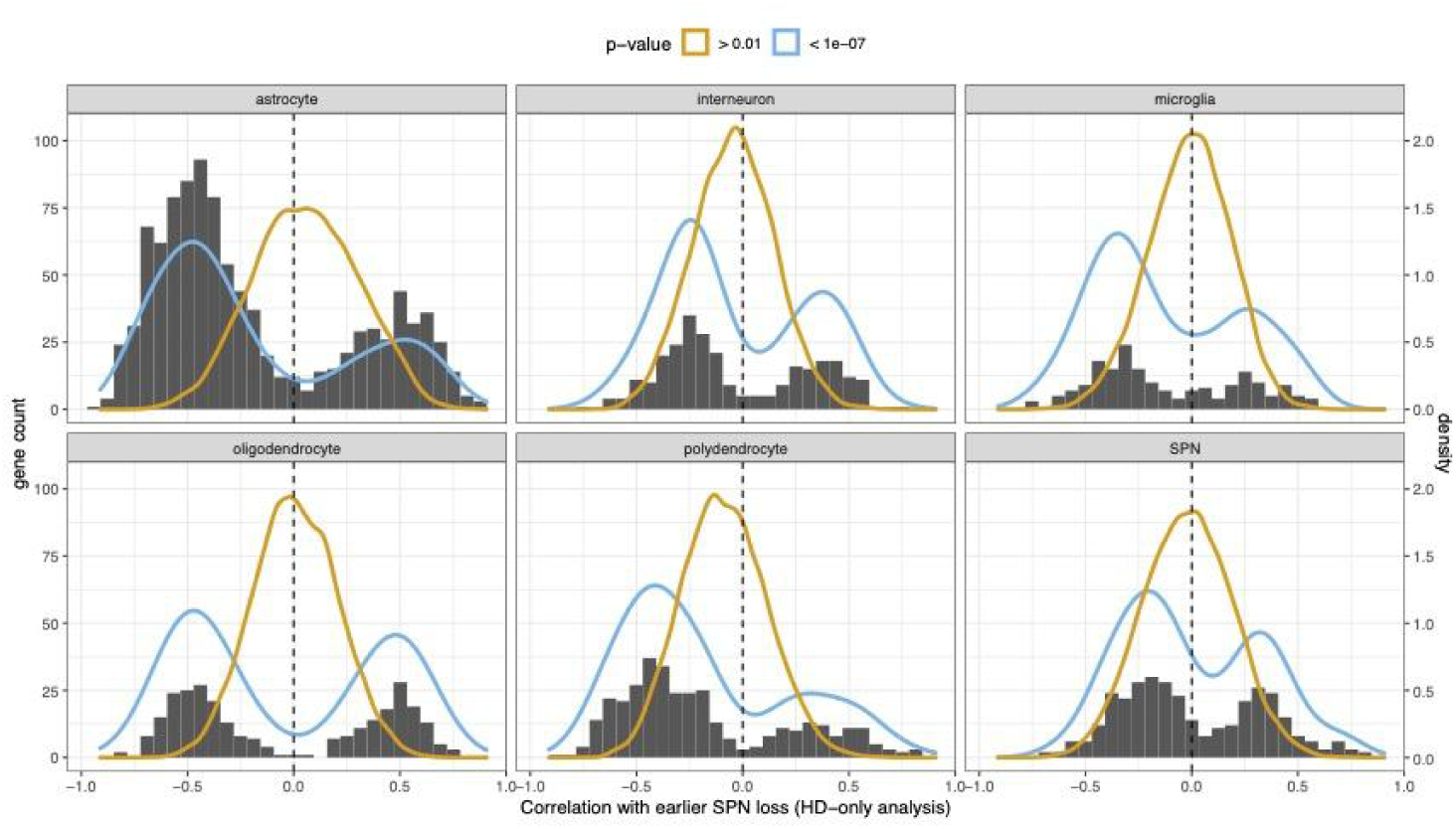
In each cell type, genes whose expression levels associated with case-control status (at genome-wide significance (at p < 10^-7^, blue density curves) show a systematic pattern of strong positive or negative correlation (x-axis), in an HD-cases-only analysis, with earlier caudate atrophy as estimated from the extent to which SPN loss has already occurred (bars, summarized by blue density curve). As a negative control, genes whose expression levels did not associate with case-control status (orange density curve) did not exhibit this within-cases relationship to earlier SPN loss. These results are consistent with a model in which case-control gene-expression differences in each cell type are dominated by responses to caudate atrophy and earlier neuronal loss.

Such analyses are useful for identifying biomarkers for HD progression and for understanding the responses of diverse cell types to neuronal death and circuitry changes. However, inferences of cause and effect are hard to make from case-control DEGs when so many genes change in expression levels in a highly correlated manner and in proportion to earlier atrophy.

### Detection of phase-C and phase-D genes in case-control analysis

What about the genes for which expression changes are detected in our (within-tissue) single-cell allelic-series analysis (Fig. 3**-5**)? Are these expression effects detectable in conventional case-control comparisons?

We found that the phase-D genes (that are repressed in SPNs with CAG repeats of 35-150 CAGs, but de-repressed in SPNs with particularly long somatic expansions, Fig. 5) exhibited clear case-control differences in the expected direction (greater expression in HD than in unaffected controls); in fact, almost all of the genes with large fold-changes in HD were phase D genes, and these were among the most significant results (**Fig. SN1.3a**). Why then has this pattern not generally been recognized by earlier case-control comparisons (whether by single cell or bulk/sorted-SPN approaches)? We found that because the expression levels of these genes are low, and there are only a small number of phase-D cells present at any moment in time, these genes have often fallen below the detection/filtering thresholds of earlier studies. (Notably, a 2015 study that used bulk RNA-seq with deep sequencing did detect expression of HOX and other homeobox genes in HD (Labadorf *et al*., 2015), and a study of microRNAs also recognized ectopic expression of microRNAs from the HOX gene clusters in HD (Hoss *et al*., 2014).)

**Figure SN1.3.**
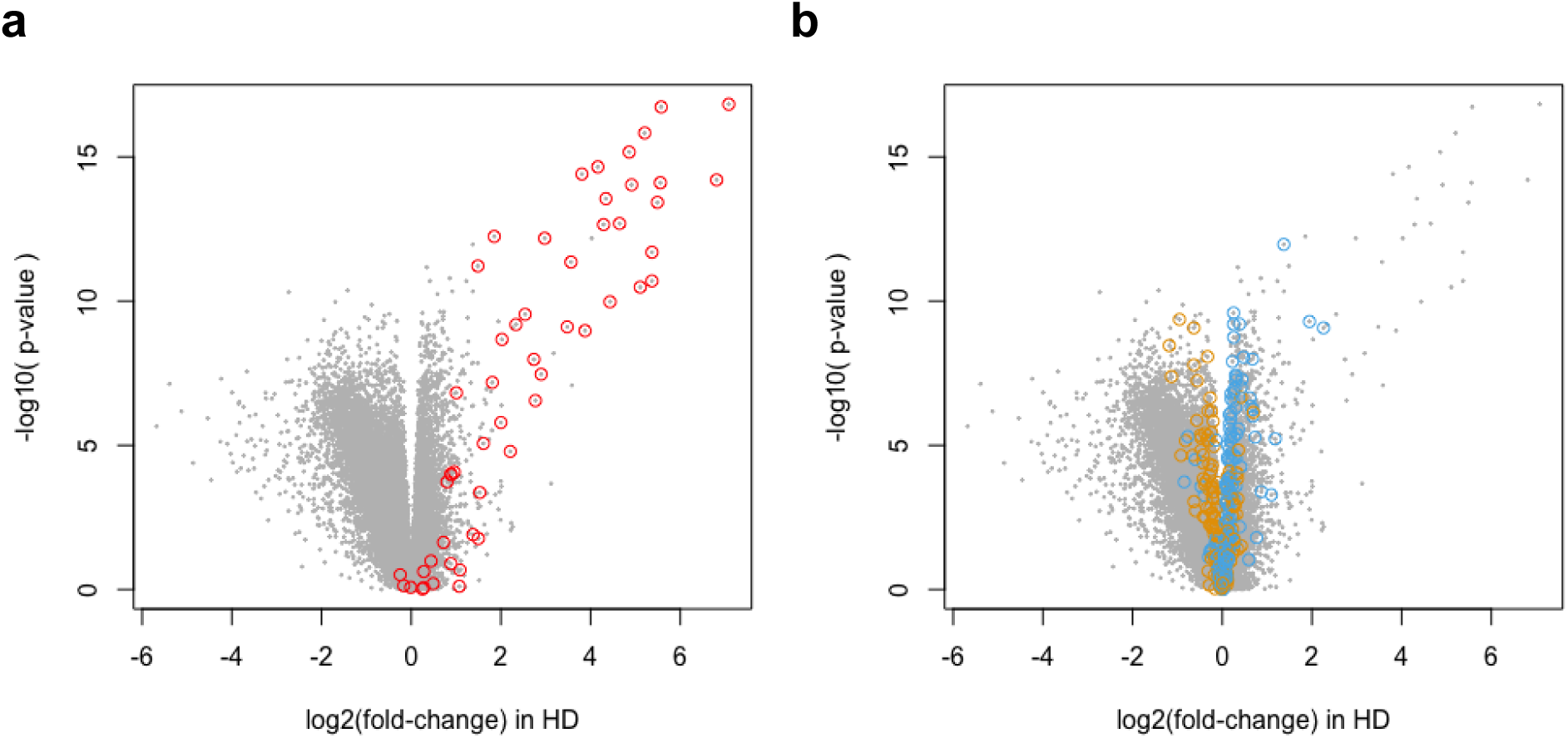
Volcano plots showing the results of an SPN HD-vs-control differentially-expressed-genes analysis comparing gene expression levels in 56 unaffected individuals to 53 persons with HD (Wilcox test). Each gene is shown as a gray point. Genes detected as de-repressed in a distinct analysis (the within-donor single-cell allelic-series analysis) are highlighted in color. (**a**) Phase-D genes, which are normally repressed in SPNs but are de-repressed in a small subset of SPNs with particularly long somatic CAG-repeat expansions, are highlighted in red. (**b**) Phase-C genes, which exhibit up-regulation (blue) or down-regulation (orange) to the extent that a cell’s own CAG repeat has expanded beyond 150 CAGs, are highlighted in blue and orange respectively.

What about phase C genes, which are expressed highly at baseline and exhibit continuous, quantitative changes in expression to the extent the CAG repeat has expanded beyond 150 CAGs (Fig. 4)? We find that, in case-control comparisons, their direction of effect analysis is in general the direction predicted by the single-cell allelic-series analysis (**Fig. SN1.3b**). However, these appear as small effects (median fold-change of 7%) and are in general dwarfed by the much larger apparent fold-changes detectable for genes whose expression levels have changed in all SPNs regardless of CAG-repeat length (**Fig. SN1.3b**).

Why are these changes so hard to recognize in a human case-control analysis? It appears to reflect two important effects. First, phase-C changes are present in just a small fraction of SPNs at any one time, and are initially extremely modest (until the CAG repeat has further expanded to 250+ CAGs in a still-smaller minority of cells, Fig. 3). Second, the phase-C genes, like most human genes, exhibit substantial natural inter-individual variation in expression levels. In the context of natural human variation, the effects of phase C changes (which are pronounced in just a tiny minority of SPNs at any moment in time) can be lost in the noise of inter-individual variation; of the fraction of these genes that do reach genome-wide significance, most do so with small apparent fold-changes (**Fig. SN1.3b**).

Note that in a comparison of the HD Q175 mouse model to wild-type mice, a set of gene-expression changes analogous to our phase-C changes are recognized strongly (Malaiya *et al*., 2021). We believe this reflects that, in the Q175 mouse, all SPNs are experiencing a toxic mutant HTT simultaneously, whereas in human HD they are doing so asynchronously.

Note in **Fig. SN1.3b** that there are a small number of phase-C-genes (orange circles) with larger effect sizes and more-significant case-control differences. These include *ANO3* and *PDE10A*. We find that these genes are affected by *both* a cell’s own CAG-repeat length (in the single-cell allelic-series analysis) and by cell-nonautonomous effects (that are experienced by all of a donor’s SPNs regardless of CAG-repeat length, but in proportion to earlier SPN loss). Their expression in all SPNs declines with HD progression (**Fig. SN1.4a**), and in addition, like other SPN-identity genes, *ANO3* and *PDE10A* are *also* genes that decline during phase C (**Fig. SN1.4b**).

**Figure SN1.4a.**
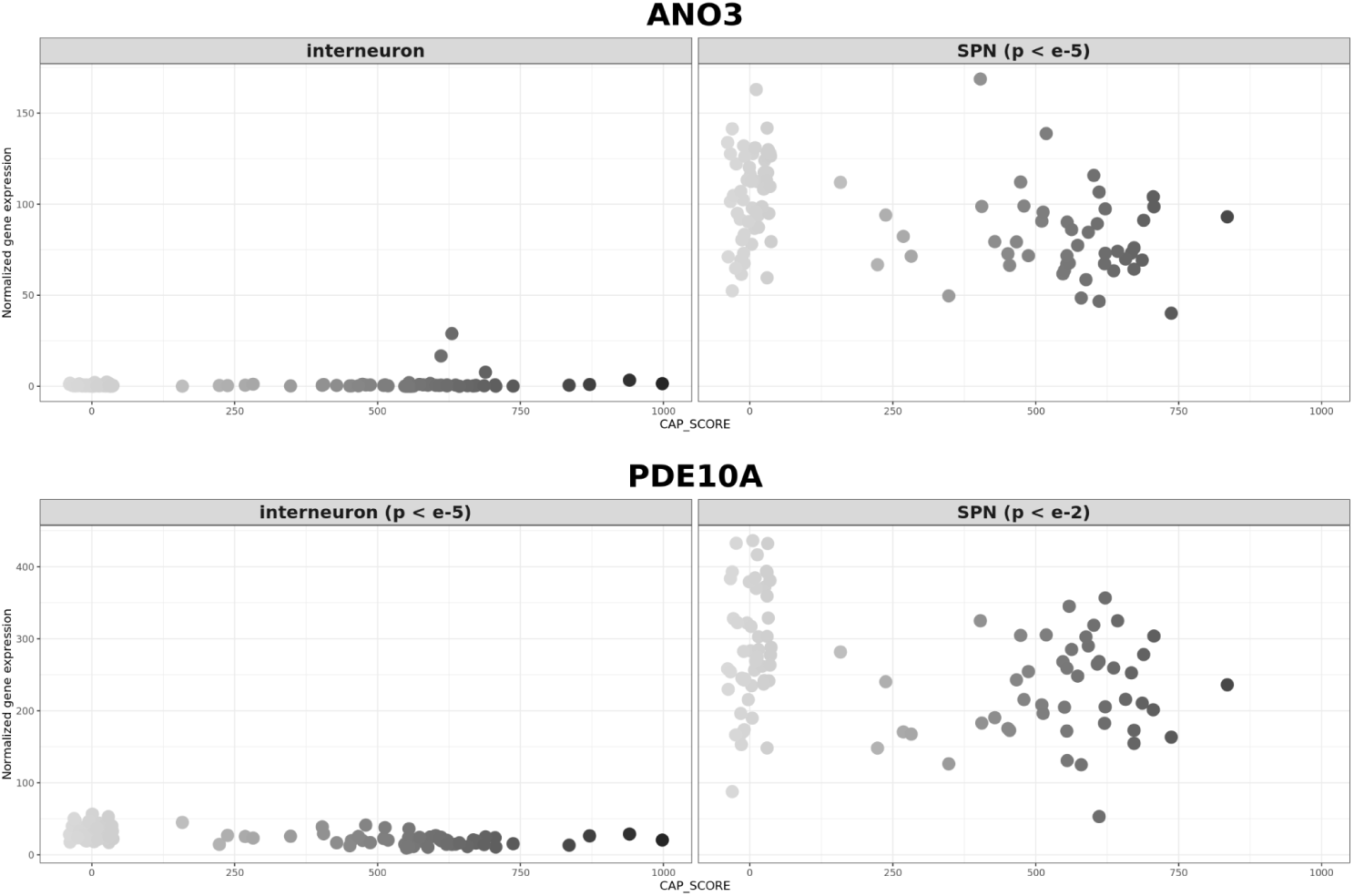
Expression of *ANO3* and *PDE10A* in SPNs across individual donors.

**Figure SN1.4b.**
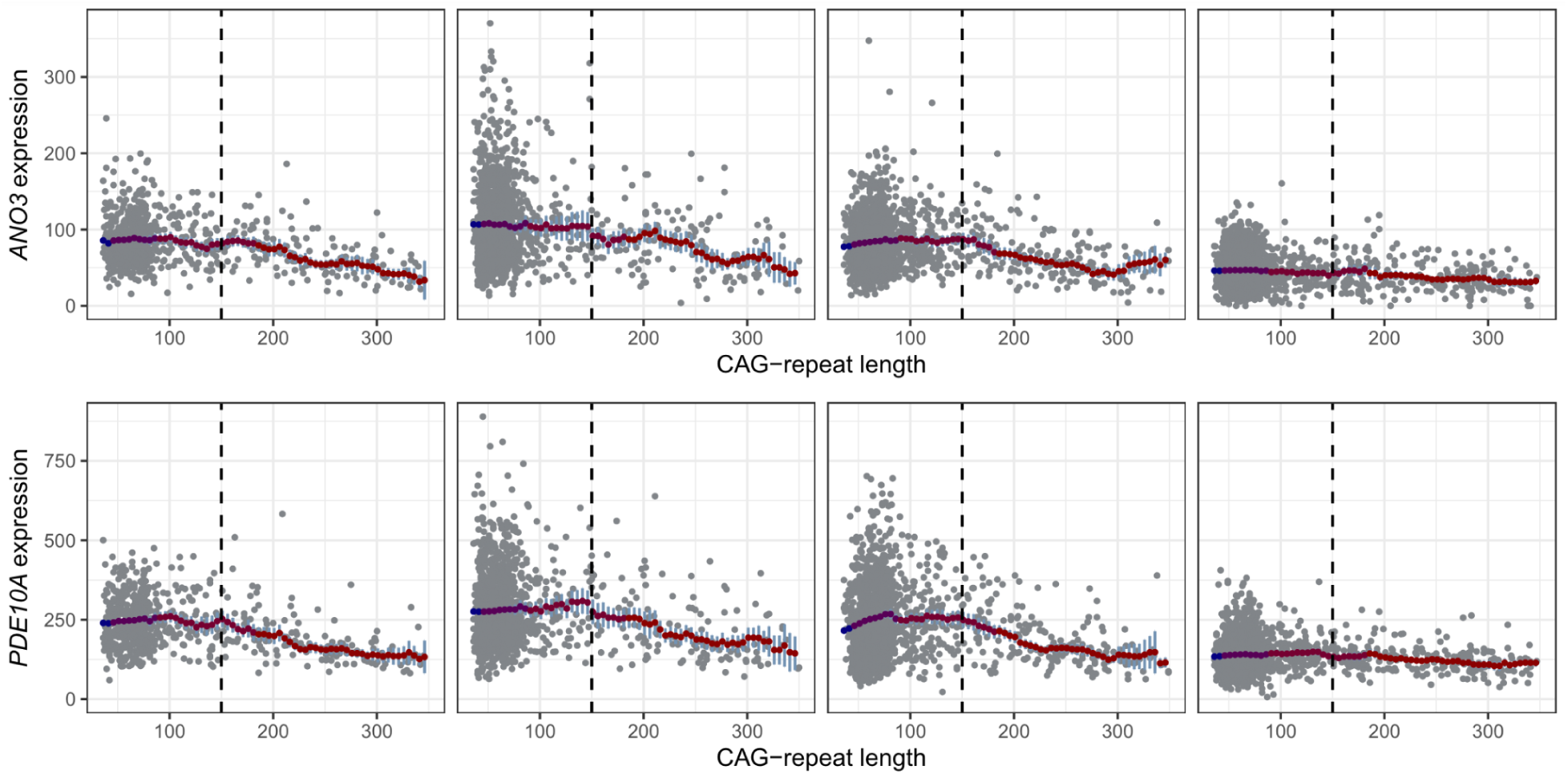
Expression of *ANO3* (top) and *PDE10A* (bottom) across individual SPNs from the same donor. Within any one donor, expression of these genes does not associate with CAG-repeat length across the (40,150) range, but it exhibits a further decline in those SPNs that have expanded beyond 150 CAGs.

Thus, *ANO3* and *PDE10A* are affected *both* by a neuron’s own CAG-repeat length (to the extent that is has expanded beyond about 150 CAGs) and also by disease sequelae (that are shared by all of a donor’s SPNs regardless of their CAG-repeat lengths). A recent study (Matlik et al., 2024) filtered a large set of HD case-control DEGs to identify genes that were found to be DEGs in SPNs but not interneurons, and highlighted *ANO3* and *PDE10A.* We note that such an approach can implicitly identify SPN-identity genes, and indeed *ANO3* and *PDE10A* are both expressed >10 times more strongly in SPNs than in any other striatal cell type (**Fig. SN1.3**).

### Summary

For these reasons, case-control “differentially-expressed gene” (DEG) analysis in human HD brain tissue tends to be dominated by a large number of disease effects that: scale with earlier caudate atrophy; involve large fold-changes because they are present in all cells regardless of their CAG-repeat length; and generally do not (except when also involving SPN-identity genes) correspond to the cell-autonomous effect of the expanding *HTT* CAG repeat in SPNs.

## Supplementary Note 2: Somatic expansion in caudate cell types other than SPNs

A recent study found that SPNs and cholinergic interneurons had similar amounts of somatic expansion and concluded that, therefore, somatic expansion is not sufficient to explain the relative vulnerability of SPNs in HD (Mätlik *et al*., 2024). Our own data suggest that SPNs have much more somatic expansion than cholinergic neurons do (**Fig. SN2.1**), and in addition that SPNs have far more expansions in the range (150+ CAG repeats) that we consider to be biologically consequential. Somatic expansion in cholinergic interneurons is also variable across individual persons with HD (**Fig. SN2.2, SN2.3**). Finally, we consider it possible that some cholinergic interneurons are indeed lost in at least some persons with HD; in our own data, in persons with HD, cholinergic neurons exhibited a modest decline in HD, though this result was only nominally significant (p = 0.02). Our conclusion is that SPNs have much greater somatic expansion and vulnerability than cholinergic neurons do, and that additional factors beyond somatic expansion into the meaningful range (150+ CAGs) are not needed to explain the relative vulnerability of SPNs.

**Figure SN2.1.**
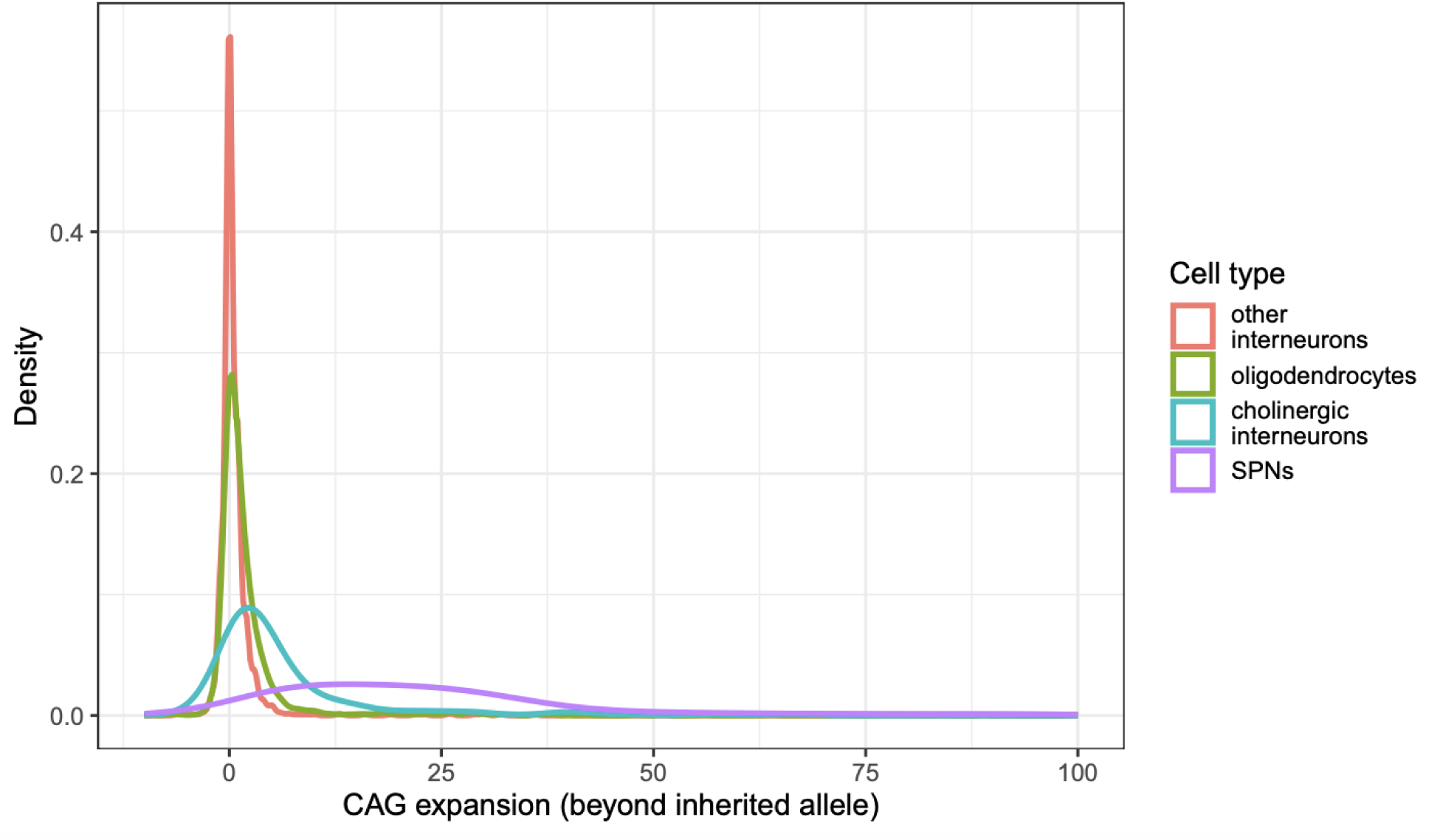
Relative amounts of somatic expansion (density plot) in selected caudate cell types. Smoothed density plot showing the degree of somatic expansion, combined across six donors, in SPNs, oligodendrocytes, cholinergic interneurons and other caudate interneurons.

**Figure SN2.2.**
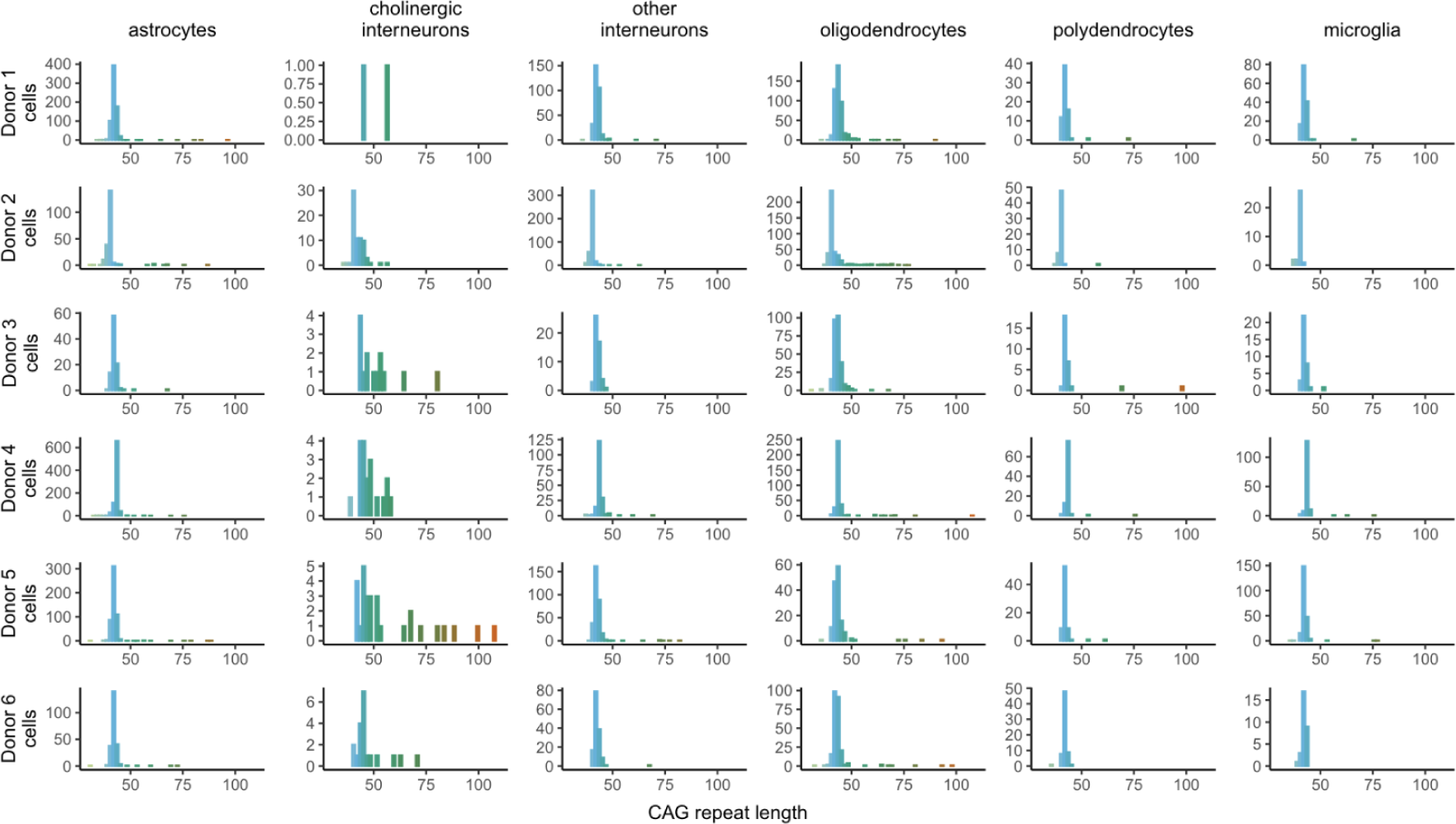
Somatic expansion in caudate cell types other than SPNs. In most cell types, the *HTT* CAG repeat exhibited only modest somatic instability. An exception was the cholinergic interneurons, a sparse caudate cell type (comprising a median of 1.4% of sampled nuclei) which exhibited more expansion than other non-SPN cell types did. Oligodendrocytes consistently exhibited more somatic expansion than other glial cell types, but less than cholinergic interneurons, which in turn exhibited far less expansion than SPNs did (see **Fig. SN2.3**). Data is shown only for long HD-risk alleles.

**Figure SN2.3.**
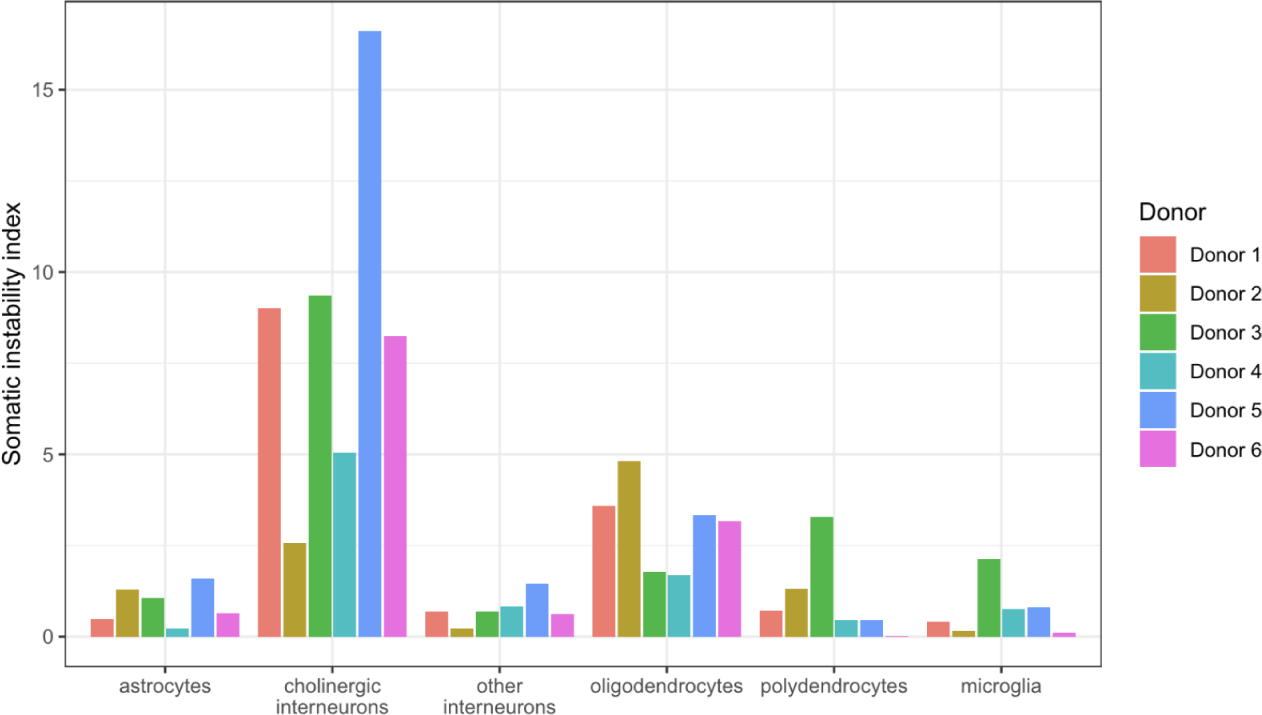
Somatic expansion in caudate cell types other than SPNs. Here the data for each of the six deeply sampled donors in Fig. SN2.2 (denoted here by the colors of the bars) are summarized with a somatic instability index, which was calculated as the mean difference between the long CAG repeat allele in each cell and the inherited repeat length in each donor (uncommon outliers beyond 100 CAG repeats have been Winsorized).

## Supplementary Note 3: Modeling the effect of CAG-repeat length on gene expression in SPNs

### Introduction

We sought to explore a variety of models and functional forms for the effect of CAG-repeat length upon gene expression in SPNs, and to identify the genes whose expression is affected by CAG-repeat expansion in a cell-autonomous manner.

Our finding (Fig. 3**-4**, **Supplementary Fig. 4-6**) that there were no apparent effects of CAG-repeat length in the 40-to-100 range was surprising, given that this is a range in which the CAG repeat has previously been assumed to have real and escalating effects upon human biology, and is the only range in which it is measured in most human brain studies. We thus sought to challenge our conclusion by exploring the possibility that there were more-modest cell-autonomous effects in this interval that a more-sensitive analysis would detect. We also sought to optimize the identification of genes whose expression is affected by CAG-repeat expansion.

### Approach

In order to identify genes whose expression changes as the HD-causing *HTT* allele’s CAG-repeat expands, we considered a number of Generalized Linear Regression Models (GLMs) containing CAG-repeat length (CAG_LENGTH) as one of the predictive variables which contribute to a gene’s expression level. We fitted these models using the single-cell-resolution data from the SPNs from all six (6) deeply sequenced individuals with clinically apparent HD. Some of the models were fitted on all the SPNs, while in others we considered only the SPNs with CAG-repeat length in a specific range (e.g. between 36 and 100), to better recognize any changes within this range and make sure the gene expression signal from the CAG_LENGTH term was not distorted by SPNs with far-longer CAG lengths.

Taking into consideration that single-cell-resolution gene expression data are consistently overdispersed (Choudhary and Satija, 2022), we decided to utilize Negative Binomial Regression (NBR) models, which model the errors in GLMs using the Negative binomial distribution. Unlike the Poisson distribution, in which the variance 𝒱 is equal to the mean μ, in the Negative binomial distribution 𝒱 > μ, making it a more suitable distribution for modeling overdispersed data.

In more formal terms, for each gene *g*, we modeled the levels of its expression *E_g_(c)* (number of UMIs), in a cell *c* with a mutant CAG allele length measurement CAG_LENGTH*_c_* by fitting the following generic negative binomial regression (NBR) model:

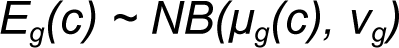

where the log mean of this negative binomial model, log(*μ_g_*), is a linear function *L* of the following cell covariates:

*N_c_* is the total count of UMIs (RNA transcripts from any gene) in cell *c f_m,n_(CAG_LENGTH_c_)* is a function family (parameterized by positive integers *m* and *n such that m < n*) of the CAG-repeat length of the HD allele in cell *c*. *f* can be a simple function of CAG_LENGTH (for example, it can be CAG_LENGTH itself), or can be defined for example as a “hinge function” (in which CAG_LENGTH has no effect until a threshold m is reached, then has a continuously escalating effect with further increase in CAG_LENGTH):

*SPN_DI_c_* indicates whether cell *c* is a direct or indirect SPN

*SPN_MP_c_* indicates whether cell *c* is a matrix or patch (striosome) SPN

*DONOR_c_* is the donor from whom cell *c* was sampled. Donor-level effects implicitly include the effects of age, genetic background, disease stage, and earlier caudate atrophy.

We fitted and compared a number of such negative binomial regression models with *μ_g_(c) ∼ L(…)(c)* that differed in (i) the choice of which CAG repeat-length ranges (i.e. which SPNs) were used to fit the model, (ii) the function *f(CAG_LENGTH),* and (iii) the presence or absence of a binary term *isPhaseD* which, when included, can distinguish between the continuously changing (phase C) and the more-discrete (phase D) effects. (This is described further below.) Since the terms “log(UMI) + SPN_DI + SPN_MP + DONOR’’ were present in all the models we fitted and compared, in the subsequent model definitions we denote them for brevity as AMT (additional model terms).

### Over what range does an SPN’s CAG-repeat length affect its gene expression?

To explore this, we fit a series of single-cell-resolution SPN gene-expression models with the same covariate terms “CAG_LENGTH + AMT” on the data with the CAG-repeat length in the intervals (35, 100], (100, 150] and (150, 300]:

Model structure: *CAG_LENGTH + AMT*

Analysis A: fitted using (n=5,109) SPNs with 35 < CAG_LENGTH <= 100 Analysis B: fitted using (n=372) SPNs with 100 < CAG_LENGTH <= 150 Analysis C: fitted using (n=551) SPNs with 150 < CAG_LENGTH <= 300

**Fig. SN3.1.**
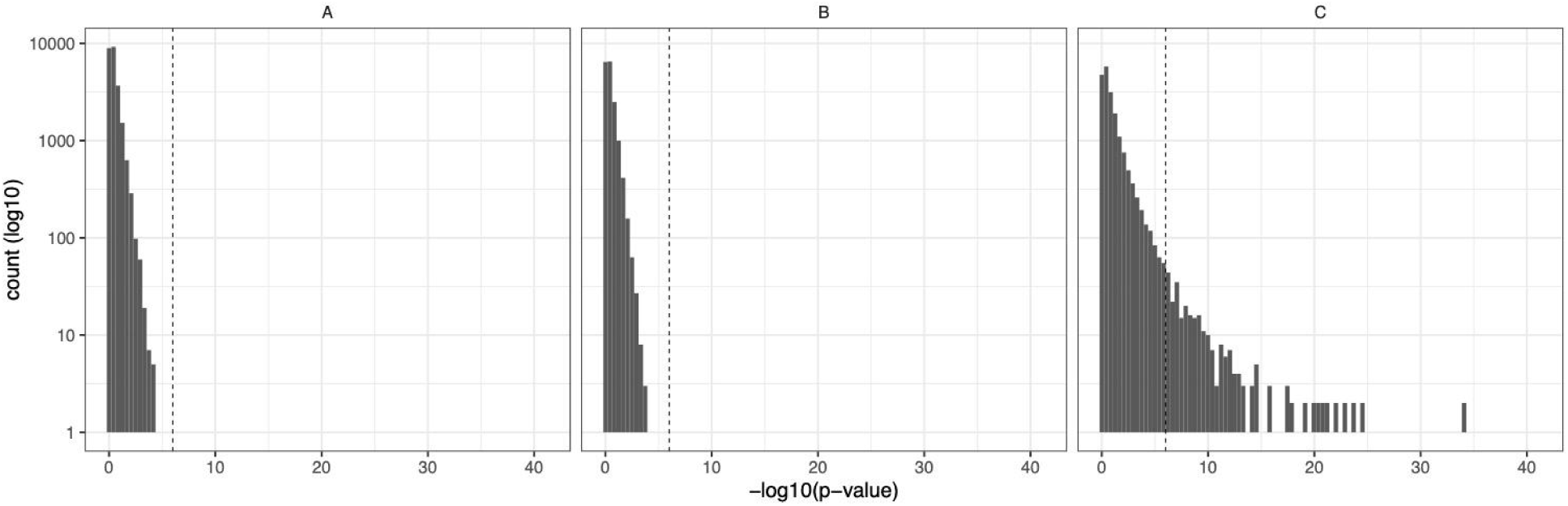
Histograms of the CAG_LENGTH terms’ -log_10_(p-values) (for each gene) in the NBR analyses A, B and C.

Analyses A and B, which were run on the SPNs in the repeat-length intervals (35-100] and (100-150] CAGs respectively, showed generally null distributions of p-values for the CAG_LENGTH term, with only a handful of genes for which this term was highly significant (p < 10^-6^). However, Analysis C, which was run on just the 551 SPNs with 150 to 300 CAGs, identified 296 genes for which the CAG_LENGTH term was highly significant (p < 10^-6^). This provides additional evidence (beyond the comparisons of groups of SPNs binned by CAG-length ranges in Fig. 3 and **Supplementary Fig. 3-6**) that little if any detectable gene-expression change results from somatic expansion from 40 to 150 CAGs.

On many plots of the SPN gene-expression data, changes become visually apparent at about 180 CAGs (**Supplementary Fig. 14, Fig. 4a**). We sought to understand whether a more-sensitive mathematical analysis could confidently place these changes as commencing at an earlier (shorter) CAG-repeat length than 180. To do that, we next fitted the gene-expression data for all SPNs with a repeat-length measurement for the HD-causing allele, using either the CAG-repeat length itself as the CAG_LENGTH term, or a “hinge function” of the CAG-repeat length in which the CAG_LENGTH term is assumed to have no effect until some repeat length *l*, and then has a linearly escalating effect to the extent it has expanded beyond length *l*.

(D) “Naive linear model” CAG_LENGTH + AMT fitted using 6,236 SPNs with CAG_LENGTH > 35
(E) “Hinge-at-150 model” f_150,500_(CAG_LENGTH) + AMT fitted using 6,236 SPNs with CAG_LENGTH > 35
(F) “Hinge-at-180 model” f_180,500_(CAG_LENGTH) + AMT fitted using 6,236 SPNs with CAG_LENGTH > 35

**Fig. SN3.2.**
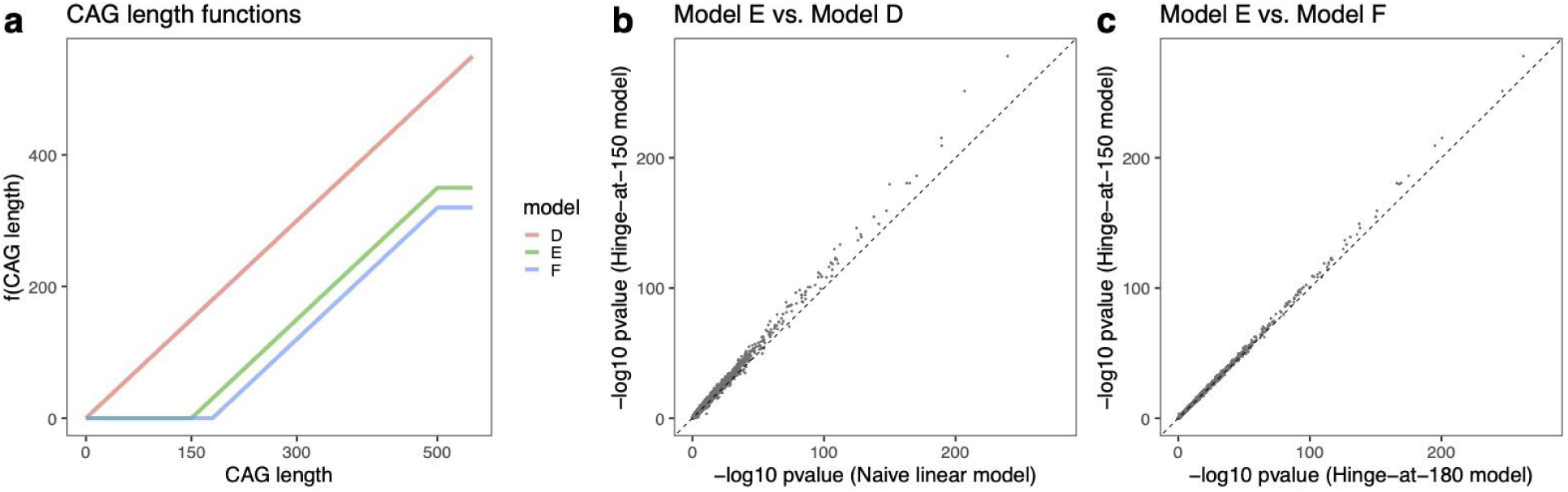
Evaluating potential thresholds for the effect of CAG-repeat length on gene expression in SPNs. (**a**) Functions of CAG-repeat length -- the identity function and two “hinge” functions in which effects commence at 150 and 180 CAGs -- used in the models D, E and F respectively as the f(CAG_LENGTH) term. (**b**) and (**c**) Comparisons of gene-level -log_10_(p-values) produced by Model E (“hinge-at-150”) (y-axis) vs. those produced by Models D and F respectively. The consistent tendency of genes with significant p-values to be located above the y = x diagonal – implies a systematically better fit by the hinge-at-150 model.

Note that these models are highly correlated with each other, and we must rely on a small number of SPNs with intermediate CAG-repeat lengths (150 to 180 CAGs) to provide the information with which to try to arbitrate between model E (hinge at 150 CAGs) and model F (hinge at 180 CAGs). We nonetheless see a clear pattern in which, for genes whose expression is well-predicted by the models, expression tends to be better predicted by the hinge-at-150 model than by the hinge-at-180 model, and substantially better than by the naive linear-effect model. Importantly, there is no set of “dissenting” genes that favor either the naive linear-effect model (in which repeat-expansion is consequential across its entire range) or the hinge-at-180 model.

This analysis provided evidence that gene-expression changes had commenced (however modestly) prior to 180 CAGs. We did not find similar evidence establishing commencement of changes prior to 150 CAGs, though we cannot formally exclude the possibility of there existing genes with very small expression changes or genes that are expressed in very few cells as such expression changes might fall below the sensitivity level of this analysis.

### Are there multiple phases of pathological gene-expression change?

From the hinge-at-150 model (which appeared to have the most power to identify effects, **Fig. SN3.2**), we identified 1,839 genes whose expression levels associated significantly (p < 10^-10^) with CAG-repeat length change in this way. Among these genes, we noticed two large subsets with somewhat different patterns of change: one subset (generally genes that were highly expressed at baseline) appeared to change expression in a continuous, graded manner to the extent the CAG repeat had expanded beyond 150 CAGs; while another set of genes (generally genes that were repressed at baseline) appeared to become expressed together in a specific subset of SPNs – generally a subset of SPNs with still-longer expansions (>250 CAGs). We sought to understand whether CAG-length-driven gene-expression changes in SPNs could be better modeled by incorporating an additional pathological phase of gene-expression change into the model – i.e. by models that included terms for both a continuous phase (“phase C”) and a subsequent, discrete phase (“phase D”) of gene-expression change.

We identified a set of 173 candidate phase D genes that are normally repressed in SPNs but had become de-repressed in a subset of SPNs with CAG-repeat length > 150 (Fig. 5a). From these genes we identified SPNs (across the six donors) in which our experiments had detected 5-174 UMIs (instead of the typical 0-1 UMIs) from these genes together. We call these “phase D SPNs”.

In order to test whether adding a covariate term for the probability of a neuron being in the phase D state would increase explanatory power of the gene expression NBR models, we defined a cell-level covariate isPhaseD as the following logistic function of the total count of the phase D gene transcripts (#phaseD UMIs):

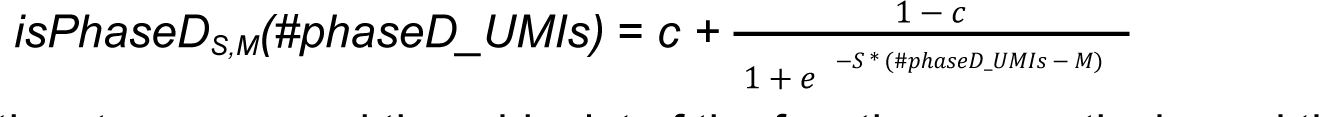

where S and M are the steepness and the midpoint of the function respectively, and the constant c is calculated so that *isPhaseD_S,M_(0) = 0*

We then defined a family of new models parameterized by these positive numbers *S* and *M*, by adding the term *isPhaseD_S,M_*to the hinge-at-150 model:

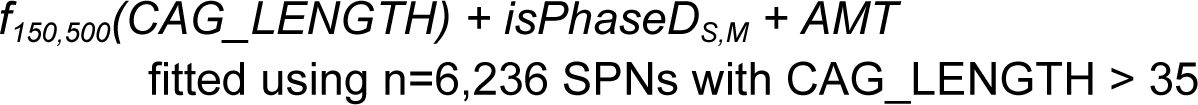

We fitted models with this structure for all the pairs of (S, M) in the direct product of {0.1, 0.25, 0.5, 1} x {0, 1, 2, 3, 4, 5, 10}, then performed the Akaike information criterion (AIC) comparisons for each pair of such models. While the AIC values for a given gene produced by these models are quite similar, slightly better overall results are produced for S = 0.5 and M = 0. Thus we selected model G to be of the following form:

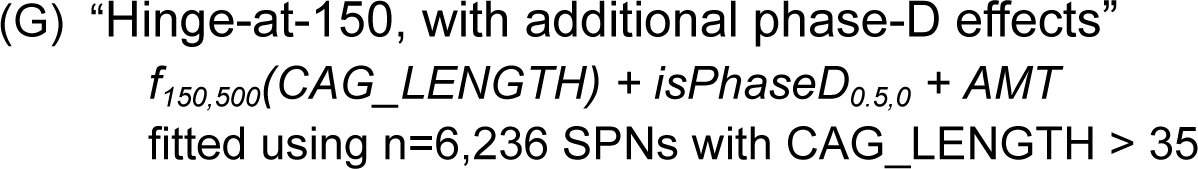

Since Model G includes an additional term (isPhaseD), we cannot compare it to the earlier models using the p-values for any one term, nor the overall fit. Therefore, to compare Model G to Model E, we also used the AIC values produced by each of these models:

**Fig. SN3.3.**
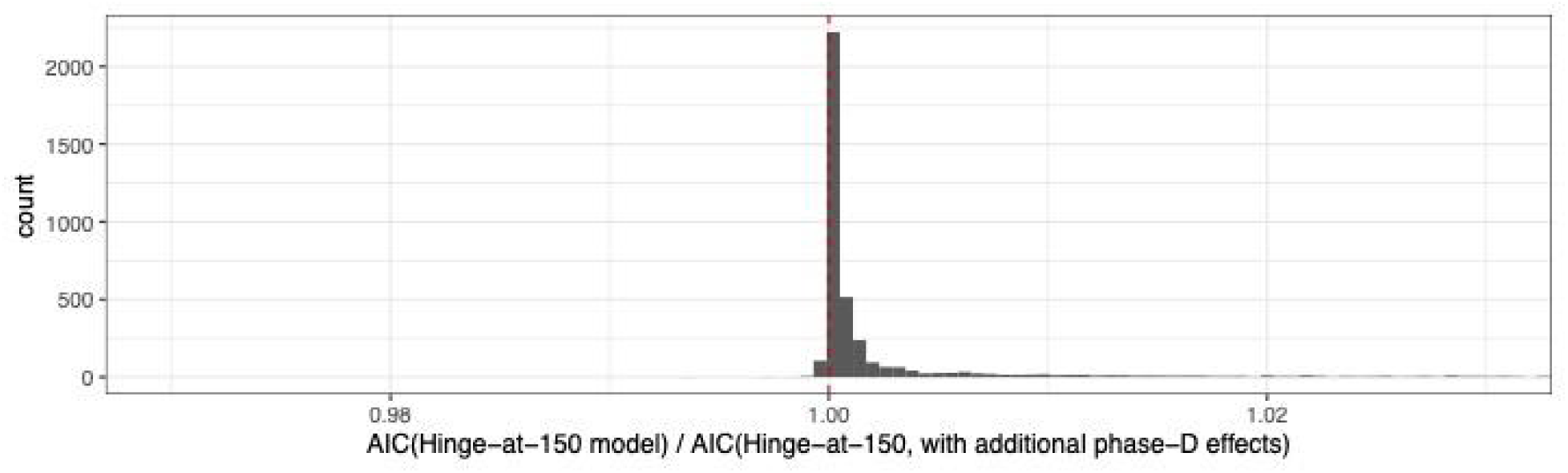
Histogram of the ratios of the AIC values from Model E (hinge-at-150) to the AIC values from Model G (hinge-at-150, with additional phase-D effects). The consistently lower AIC values for Model G (AIC ratio > 1) implies that Model G fits the data better than Model E does.

Most of these ratios are greater than 1, demonstrating that for the majority of genes whose expression is significantly affected by the CAG repeat length, adding the isPhaseD term to the hinge-at-150 model improved its explanatory power.

### Phase C and phase D genes

Having determined that the model with both continuous phase-C effects (commencing at 150 CAGs) and an additional phase D effect provides the best overall fit to the data, we can now ask which genes change in expression in phase C, and which genes change in expression in phase D.

We plotted here the log_10_-p-values (signed by direction of effect) of the terms *CAG_LENGTH* and *isPhaseD* for each gene (**Fig. SN3.4**):

**Fig. SN3.4.**
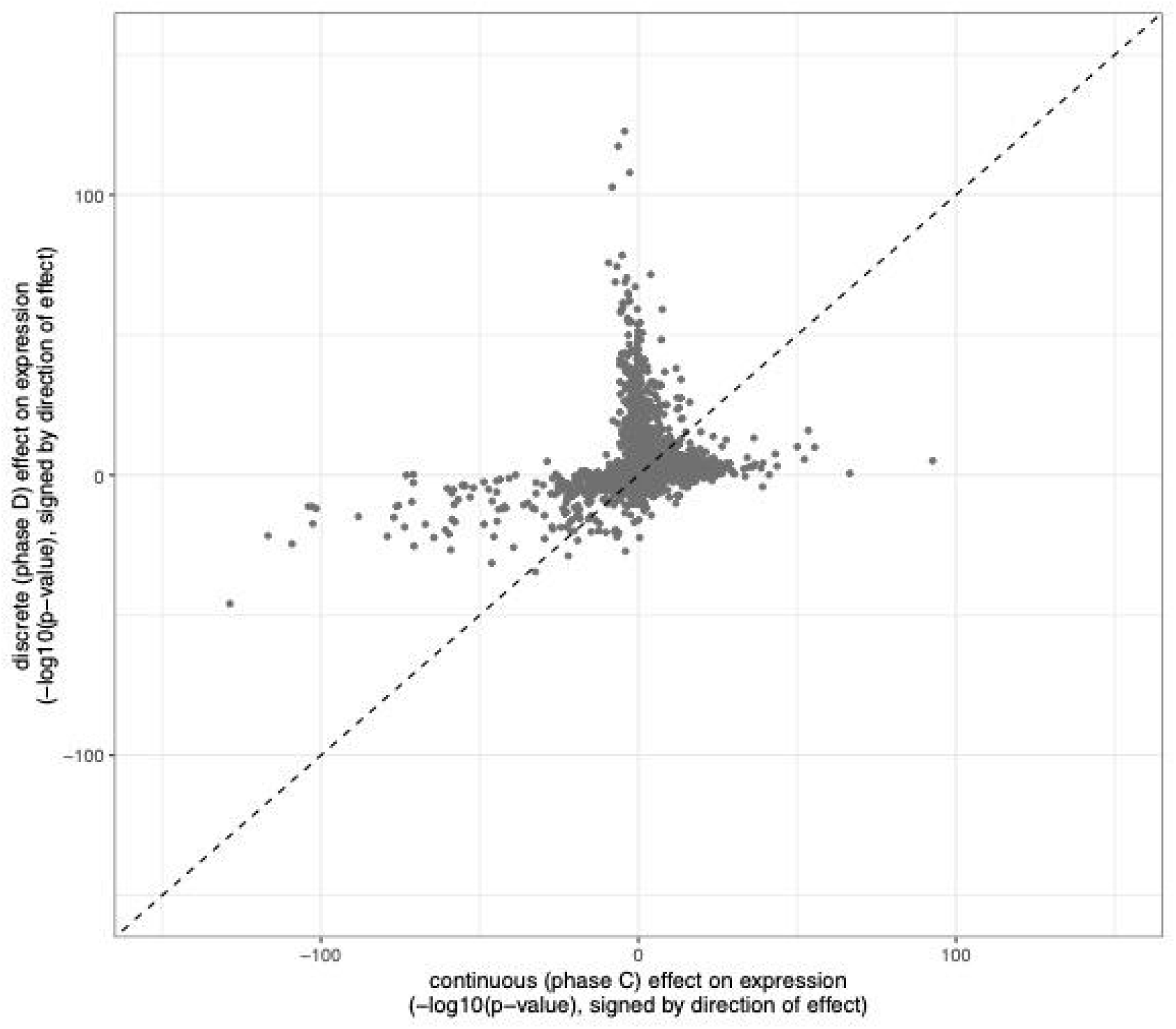
Scatter plot of the strength and direction of continuous effect on gene expression of the CAG_LENGTH term of model D (x-axis) and the strength and direction of discrete effect on gene expression of the isPhaseD term of model D (y-axis). For each gene, the magnitude of the value in each dimension is the -log10(p-value) of the term and the sign indicates the direction of effect. The lack of correlation indicates that most genes change either in a continuous fashion (as a function of repeat length) or in a discrete event in concert with other phase D genes.

Note how individual genes tend to be regulated in either the phase-C (continuous change) or phase-D (discrete genome-wide crisis in specific cells) pattern. This is seen in the tendency of points to fall along the x- or y-axes. (Note, though, that our model imperfectly parses these two effects, in substantial part because of a population of “borderline phase D” SPNs; this gives rise to the tendency of points with positive changes in phase C (points along the right side of the x-axis) to also have slightly positive (though generally insignificant) associations with phase D.)

Phase C appears to involve a mixture of positive and negative effects on gene expression (as also illustrated in Fig. 3**-4**). Phase D appears to consist almost entirely of positive effects (increasing expression), generally on genes that were repressed at baseline.

### Conclusion

These results support a model in which direct, cell-autonomous effects of the CAG repeat upon an SPN’s own gene expression commence at a high CAG-repeat-length threshold of approximately 150 CAGs and traverse multiple stages characterized initially by continuous, quantitative gene-expression changes concurrent with further CAG-repeat expansion (phase C) and then by a rapid-onset de-repression crisis (phase D).

## Supplementary Note 4: Modeling and simulation of SPN CAG-repeat expansion dynamics

The distributions of observed CAG repeat lengths in caudate SPNs of donors with HD exhibited an unusual yet consistent shape, with a pronounced right tail consistently commencing at about 80 CAGs across many donors (Fig 2c). To better understand how these distributions might arise and evolve from the kind of simple, stepwise expansion-and-contraction process that biological studies have suggested is their primary mode of somatic mutation (reviewed in (Iyer and Pluciennik, 2021)), we developed stochastic models and simulations for the dynamics of the somatic expansion process. We aimed to see whether such models could explain these observed distributions and the typical trajectories of SPN loss in HD – particularly if, as our experiments suggested, the *HTT* CAG repeat becomes pathogenic only when it is quite long, a length attained by few SPNs at any moment in time.

### Continuous-time Markov chains

Inspiration for the classes of models we used came from prior work modeling repeat-expansion dynamics in both HD and other disorders (Higham *et al*., 2012). In particular, we developed stochastic models in which the mutation process in the HD-causing CAG repeat is a biased random walk. In each SPN, the probability of the CAG repeat either expanding or contracting, within a given time interval ***t***, is a function of its own current CAG-repeat length. Such memoryless models can be expressed as continuous-time Markov chains (CTMCs) in which the state space corresponds to the range of modeled repeat lengths and the transitions correspond to mutations that change the repeat length.

In our models, we use a state space ranging from CAG=0 to CAG=max_cag where in practice we set max_cag to 500. In our models, the state corresponding to CAG=max_cag is treated as a “sink” state where once a cell reaches this repeat length, it will stay in this state indefinitely. In practice, we pick this state to be beyond the range of repeat lengths where we would like to model the dynamics of the somatic expansion process. We use this sink state to represent all cells that have reached very long CAG lengths and, as discussed below, in most of our models we assume that the majority of such cells have been lost.

Every CTMC process can be described by a rate matrix ***Q[i,j]*** that describes the probability of transitioning from state ***i*** to state ***j*** in unit time. In our models, the time unit is years. For each somatic expansion model ***M***, we define ***Q[i,j]*** for ***M*** in terms of a smaller set of model parameters ***v*** for model ***M(v)*** and then fit these model parameters as described below. For any CTMC with rate matrix ***Q***, there exists a unique matrix ***P(t)*** where the entries ***p[i,j]*** specify the probability that a cell with repeat length ***i*** at time zero has a repeat length ***j*** at time ***t***. ***P(t)*** can be expressed as the matrix exponential

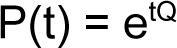

which can be efficiently computed using the expm package in R. For a specific donor, if we set ***t*** equal to the age of brain donation and assume that all cells started with a repeat length equal to the donor’s inherited repeat length (***inh_cag***), then the function

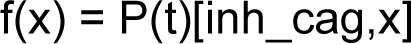

gives the probability distribution of repeat lengths for this donor we would expect to observe under the model ***M(v)***. We then fit ***f(v)***, which is a function of the model parameters for ***M***, to the observed repeat length distribution for each donor to determine the best-fit estimate the model parameters for that donor.

### Generating a rate matrix Q for a specific model

As a concrete example, consider a simple model with three parameters (***p***, ***r*** and ***T***) where the rate of mutation is a linear function of the repeat length above a threshold ***T*** and each mutation results in either a one-repeat-unit CAG expansion with probability ***p*** or a one-repeat-unit CAG contraction with probability ***1-p***. We model the mutation rate per unit time as

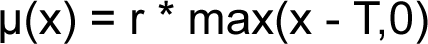

where ***x*** is the current repeat length and ***T*** is the threshold below which no expansion or contraction occurs. We can then form the rate matrix ***Q[i,j]*** where ***0 <= i,j <= max_cag*** as

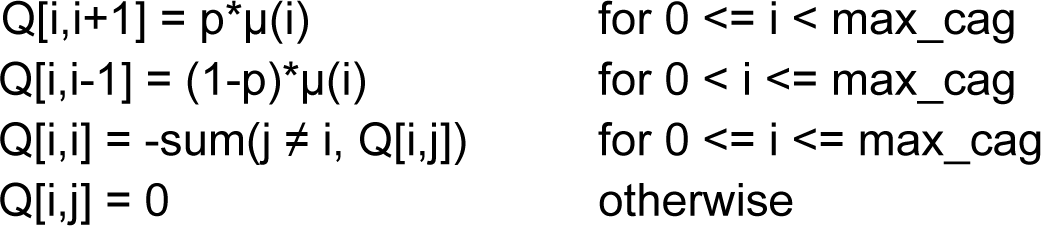

The rate matrix ***Q*** is a function of the model parameters (***p***, ***r*** and ***T***). Each of these model parameters can either be fixed or be fitted from the data.

### Estimating the amount of SPN loss for each donor

Caudate SPNs are lost during the course of Huntington’s disease (Fig. 1). From the transcriptome data, we estimated the proportion of different cell types in the tissue samples from the anterior caudate in both HD donors and controls. We found that in unaffected control donors 46% (+/- 0.06%) of the nuclei we sampled were derived from SPNs, whereas the fraction in donors with HD ranged from less than 1% to 51%, with an observed dependence on disease progression as measured by CAP score (Fig. 1) or Vonsattel grade.

We observed, based on the transcriptome data, that across persons with HD the abundance of of observed polydendrocytes (oligodendrocyte progenitor cells) tended to decline alongside that of SPNs, but that other cell types had stable proportions relative to one another, mainly expanding (as a fraction of the total) to the extent that SPNs and polydendrocytes were lost. As a result, we treated both the SPNs and polydendrocytes as cell types undergoing cell loss associated with disease progression.

To estimate the degree of SPN loss in each donor, we compared the ratio of SPNs to cells of unaffected cell types (all cell types excluding SPNs and polydendrocytes) and compared this ratio to the median ratio observed in 52 control donors. This yielded estimates of SPN loss ranging from 0% to almost 100% (**Supplementary Table 2**).

Uncertainty (noise) in these cell-loss estimates is introduced by sampling of the inherently non-uniform tissue in the caudate (e.g. sampling of different amounts of white matter vs. gray matter). For many donors we had two estimates of SPN fraction: one from the many-donor (“cell village”) experiments (Fig. 1) and another from deep resampling of individual donors alongside single-cell CAG-repeat-length measurement (Fig. 2**-4**). We used the cell-loss estimates from the village experiments (Fig. 1), whenever available, as these measurements were from a consistent anatomical site within the anterior caudate and they exhibited stronger relationships with each donor’s CAP score (Fig. 1) and with neuropathological determination of disease stage (Vonsattel grade).

### Incorporating cell loss estimates in repeat expansion models

As HD progresses, loss of SPNs is profound. Importantly, experimental measurements of CAG-repeat length distributions are measurements only of the SPNs that are still present at the end of a brain donor’s life – which will generally be a small fraction of the SPNs with which a donor began life. Any model must take into account this profound effect of attrition on the observed CAG-repeat length distributions.

We make a simple assumption, rooted in our single-cell gene-expression data – that cell loss occurs only in the right tail of the CAG-length distribution, beyond the threshold (∼150 CAGs) at which gene-expression changes begin to become detectable. We do not try to model the details of this cell-death process nor its precise probabilistic relationship to CAG-repeat length. In fact, as described in more detail below, the details of this relationship have little impact on the results, as somatic expansion beyond 150 CAGs is extremely fast compared to somatic expansion at earlier stages, e.g. 40-50 CAGs.

### Fitting a repeat expansion model for an individual donor

For each donor and each model ***M(v)***, we want to find the values of the model parameters ***v*** that best predict the observed repeat-length distribution and cell loss estimate for that donor. These model parameters define the dynamics of the repeat mutation process in that donor.

We implemented model fitting as an optimization problem. For each individual donor, we use the inherited repeat length (inh_cag), the donor’s age at brain donation (d) and a vector of observed repeat lengths (x) from N SPN cells. Given a model M with a vector of fixed parameters and a vector of parameters to be fitted theta, we used the optim package in R to find optimal values for theta. We used both the Nelder-Mead algorithm and the L-BFGS-B methods implemented in the optim package in R - both of which achieved similar results. In the final reported analyses, we used the L-BFGS-B method in the optim package along with empirically derived parameter ranges to aid with rapid convergence of the model fitting.

The objective function over which we optimized was the log-likelihood of the experimentally measured SPN CAG-repeat-length distribution under the parameterized model, with two modifications. First, all observed cells with a repeat length greater than max_cag were assigned to the last repeat-length bin (corresponding to max_cag). Second, we adjusted the likelihood to account for cell loss, based on the donor-specific estimate of the fraction of dead (and thus unobserved) SPNs, estimated as described above. We incorporated this cell-loss estimate in the following way: From the number of observed cells and the loss estimate, we calculated the number of missing/unobserved cells and added these as pseudo-counts to the last bin (corresponding to max_cag). To prevent large estimates of cell loss from dominating the model fit, any estimate for cell loss greater than 90% was capped at 90%

### Single-jump vs multiple-jump models

Within the space of CTMC models, we explored and evaluated a variety of models for somatic expansion. Our analyses ended up focusing on a set of models with a specific form based on two quantities:

μ(x) A function giving the mutation rate as a function of the current repeat length

p_exp_ The probability of an expansion (vs. a contraction), when a mutation occurs

This approach formally models processes where the repeat length mutates by exactly one CAG unit at a time (either an expansion or contraction) with the probability of an expansion being p_exp_ and of a contraction being 1 - p_exp_. These are so-called “single-jump” models, in contrast to multiple-jump models that incorporate larger mutation events by assuming some probability distribution over the change in repeat length, conditional on the occurrence of a mutation.

We evaluated several multiple-jump models, in particular models where the change in the repeat length was drawn from a geometric distribution with a maximum jump size ranging from two to five CAG units. Compared to otherwise-equivalent models with single-repeat jumps, these models generate distributions with higher variance. We found that although the multiple-jump models fit the observed data well, the fits were not substantially better than the equivalent single-jump models. As a result we concluded that we lacked sufficient power to reject single-jump models in favor of multiple-jump models with the currently available data and focused on single-jump models instead.

We reasoned that as long as *most* mutation events are short, single-jump models should have good predictive power, even if some mutations are longer than one unit. Our results indicate that single-jump models are sufficient to explain the observed data, which is consistent with a model where most jumps are, in fact, short. There are also other lines of evidence suggesting that most repeat-length mutations are likely to be short. *In vitro* studies suggest that mismatch repair is more efficient at recognizing small extrusions compared to larger ones, including in systems that are specifically based on the genes that are modifiers of HD onset (reviewed in (Iyer and Pluciennik, 2021)). In addition, the smooth, unimodal shape of the distributions observed in our data, as well as other studies in humans and mice, are consistent with large jumps being rare, at least in the early phases of repeat expansion.

In the later, more rapid phases of somatic expansion, the increased overall rate could in principle be driven either by a higher rate of mutation events, a larger average jump size per mutation, or some combination of these effects. The available data do not strongly distinguish among these possibilities, as their effects upon the repeat-length distributions and dynamics are so similar.

### Sensitivity of model-fitting to the net expansion rate

Importantly, model-fitting for both the single-jump and multiple-jump models was strongly driven by the goodness-of-fit to the net expansion rate, which is the product of the mutation rate and the mean jump length. For multiple-jump models, we found that an increase in mutation rate could be offset by a reduction in the expected jump size and vice versa to create roughly equivalent model fits.

For single-jump models, the net expansion rate is

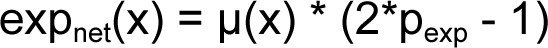

A formal assumption in our models is that although the expansion probability p_exp_ can vary between donors, it is fixed in each donor and is independent of the current repeat length.

Because the model fit is driven by the net expansion rate, it is difficult to distinguish between models where the mutation rate varies as a function of the repeat length, models where the expansion bias (p_exp_) varies as a function of repeat length, or some combination. For simplicity, we chose to model p_exp_ as a constant in each donor and allow the mutation rate μ(x) to vary as a function of repeat length.

Whether the expansion bias (or jump size) changes with repeat length will be an interesting area for future study, but will likely need the benefit of a more complete mechanistic understanding of the somatic expansion process.

### Single-phase vs. two-phase models of somatic expansion

We implemented and investigated a number of different models for somatic expansion. Broadly, we divided these models into two categories: single-phase and two-phase. The investigation of the two-phase models was driven by the observation that the empirical cumulative distributions of the repeat lengths, across donors, tended to have a sharp bend (or “kink”) around 70-90 repeats, suggestive of a rapid change in the dynamic expansion behavior in this range (Fig 6c).

Though such a kink could in principle be generated by a complex model assuming substantial SPN death at 70-90 CAGs (and further assuming that a subset of SPNs were immune to this for unknown reasons), the lack of apparent gene-expression changes across the 40-150 CAG range guided us to focus on simpler models in which this kink was entirely generated by repeat-expansion dynamics. Moreover, even if many SPNs were lost at 70-90 CAGs, the surviving SPNs would still need to have repeats that continue to expand rapidly to reach the long repeat lengths (up to 800+ CAGs) observed in our data.

It has been well established that CAG repeats of 35 or fewer CAGs exhibit very little instability and are not known to lead to Huntington’s disease. Indeed, we observed no significant instability in the short-alleles of vulnerable cell types in HD donors. Accordingly, all of the models we explored assumed that somatic expansion begins at a threshold (T1) of around 35 CAGs.

In practice, the models we used set the mutation rate to zero below a fixed threshold. Since such models create a “sink” state at the instability threshold, we found it practical to set the lower threshold, T1, to 33.5 to reduce the number of cells that would become trapped in this lower sink state.

We categorized models that used a single threshold for somatic instability (T1), whether this threshold is fixed or fitted, as single-phase models. We also explored models in which there was a second threshold (T2) at which the dynamic properties of the model were allowed to change, in particular to allow faster acceleration of the expansion rate beyond what could be achieved with common smooth functions and a single instability threshold. We categorized any model with this type of second threshold as a two-phase model. We chose to apply minimal smoothing at the transition between the two phases, requiring only continuity of the mutation rate function (and in some models its first derivative), as this was sufficient to explain the observed data. The underlying biological process quite likely does not exhibit as abrupt a transition as represented in these models.

### Models of repeat expansion

The main analysis presented here are based on two models, which we refer to as the TwoPhaseLinearModel and the TwoPhasePowerModel.

In the TwoPhaseLinearModel, the mutation rate varies as a piecewise linear function of the repeat length with three regimes: There is a threshold T1 below which the mutation rate is zero and a second threshold T2 separating the other two regimes. The mutation rate function for the TwoPhaseLinearModel is given by

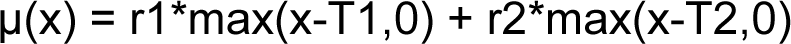

The parameters r1 and r2 are rate constants for the two non-zero regimes. The effective rate (slope) of the second of these regimes is r1+r2. In practice, we fixed T1 at 33.5 and fit T2 from the data for each donor, as well as r1 and r2.

In the TwoPhasePowerModel, the mutation rate varies as a power function of the repeat length over three regimes, similar to the TwoPhaseLinearModel. The mutation rate function for the TwoPhasePowerModel is given by

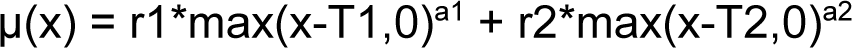

where r1 and r2 are rate constants similar to the previous model and a1 and a2 are (fitted) exponents for the two non-zero regimes. In practice, we fixed T1 at 33.5 and fit T2 from the data for each donor (along with r1, r2, a1 and a2).

We utilized these two models in different ways. The TwoPhaseLinearModel was the simplest model that gave a good fit to the observed data. We used this model to estimate and compare parameter values between donors. The TwoPhasePowerModel was potentially over-parameterized, but had the property of fitting the observed data well (better than the TwoPhaseLinearModel), at the cost of some over-fitting. We found that using these over-fitted models allowed us to compute more reliable stochastic trajectories of the cells, which were useful for further analysis. Since there is a small degree of overfitting in the TwoPhasePowerModel, we avoided comparing the specific parameter values that generate the best fits for this model, but instead compared the predicted trajectories as described further below.

As points of comparison, we also evaluated a number of “one-phase” models. The one-phase models described here include the following, listed with their corresponding mutation rate functions:

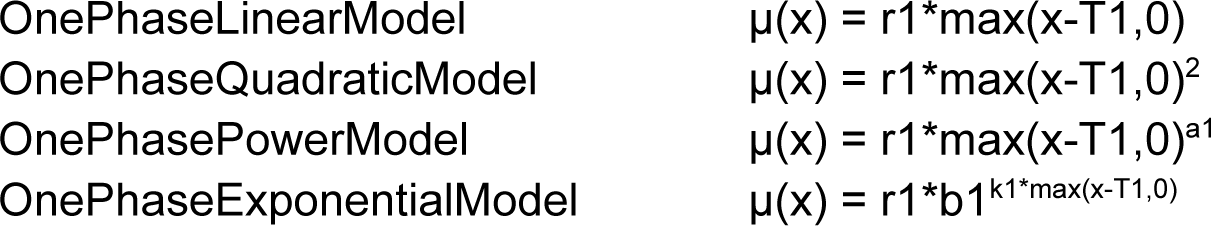

In each of these models, we fixed T1=33.5 and fit the other parameters from the data.

When we refer to a specific model by name, we generally include the repeat-length threshold used for modeling cell loss after the name of the model, separated by slash. For example TwoPhasePowerModel/150 would refer to the two-phase power model fitted using a cell loss threshold of 150 CAGs. The cell loss threshold is the minimum repeat length at which the cell loss is assumed to begin to occur, as described previously.

### Comparison of different models of somatic expansion

Across all the donors whose SPNs we had deeply sampled with CAG-repeat length measurements, the two-phase model provided better fits than the one-phase models (**Fig. SN4.1**).

We observed two important qualities of the models that were able to best fit the observed data: First, that the models allowed the rate of somatic expansion (in CAGs/year) to increase as a super-linear function of the current repeat length. Second, the best models were two-phase models that allowed the dynamics of the expansion process to further accelerate around 70-90 CAGs. We found that the exact functional forms used in the two-phase models were not as critical (given the available data) as allowing the dynamics to accelerate at this repeat length threshold. For simplicity, we utilized two-phased models that used the same functional form for both phases.

We found it useful to evaluate these models based on their predicted net mutation rate curves (in CAGs/year) as a function of repeat length (**Fig. SN4.2**). We observed that in this framework both the one-phase and two-phase models appeared to converge to a common net mutation rate curve, best represented by the two-phase power models which adhere most closely to the shape of the observed data. As a consequence, we based most of our analyses on two models: the two-phase power model, which had the best overall fit of the models we tried and the two-phase linear model. Both of these models generated good fits across donors (**Fig. SN4.3**). We used the two-phase power model, which provided the best fits overall, when comparing the expansion trajectories across donors. We used the two-phase linear model in contexts where we wanted to directly compare the fitted parameter values between donors to avoid potential over-fitting of the two-phase power model.

Importantly, our use of two-phase models was not intended to suggest that there is some additional, previously unknown, biological process contributing to somatic expansion. The introduction of two phases appeared to better model the apparent shift in the observed net rate of expansion we observed in the data. In principle, many different phenomena could account for this increase in expansion rate, including interactions with local chromatin structure or changes in the jump size per mutation event as the repeat length increases. Understanding the mechanism that underlies this aspect of the somatic expansion dynamics will be an important area for future research.

**Supplementary Figure SN4.1.**
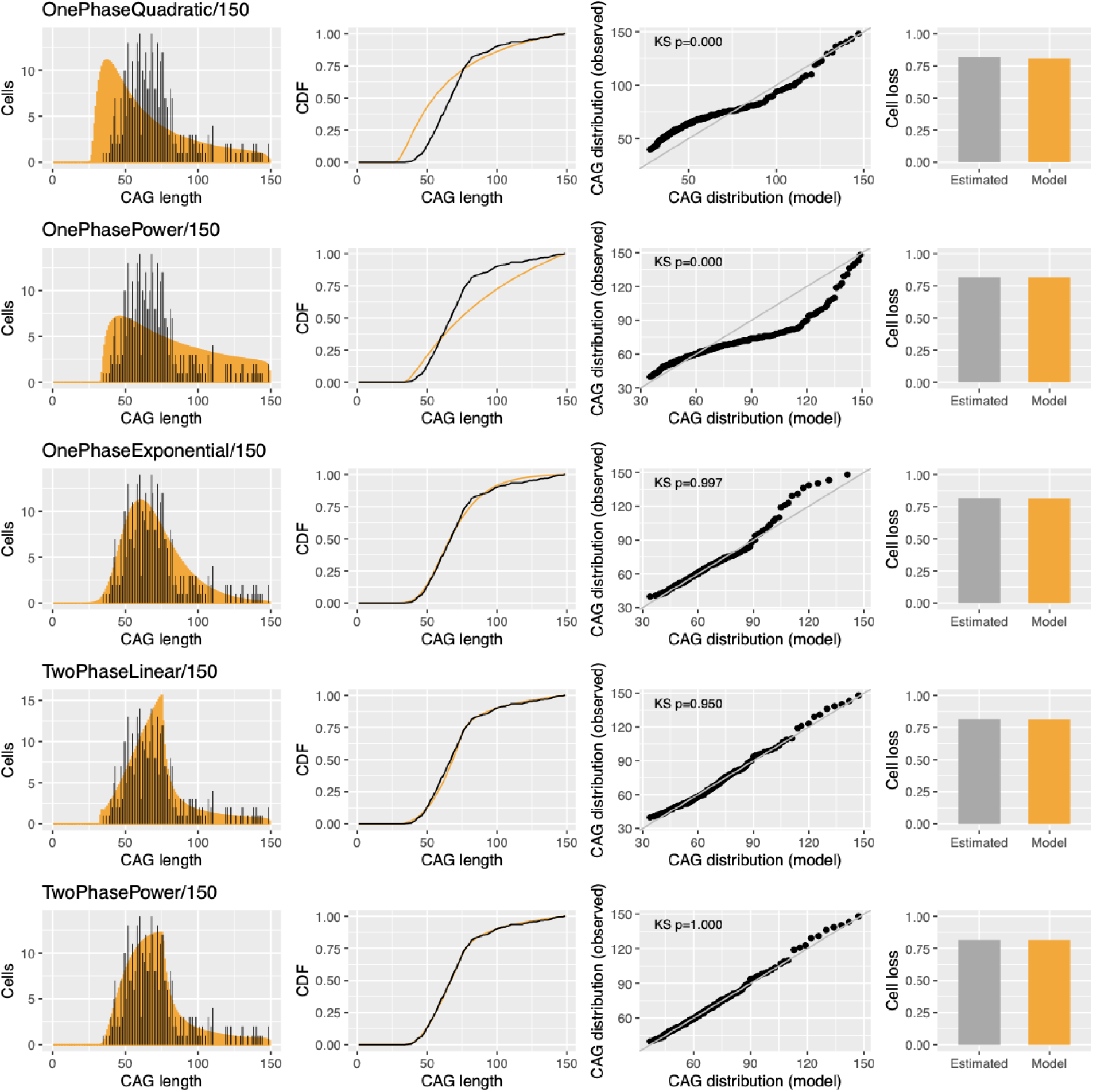
Comparison of model fits for different somatic expansion models. Comparison of the best fits for five different models to the data for one representative donor (Donor 5), including both one-phase (rows 1-3) and two-phase (rows 4-5) models. All models shown were fitted using a minimum cell-loss threshold of 150 CAGs. The two-phase models produce better fits to the observed data. Although the one-phase exponential model provides the best fit among the one-phase models we investigated, across donors it fails to replicate the shape of the long tail (overshooting the observed data in the range of 70-90 CAGs and undershooting the observed ata at longer repeat lengths). The two-phase power model provides the best fits overall. The two-phase linear model is nearly as good and we use this model in some analyses, for example to compare the overall expansion rates in the slow and fast phases of somatic expansion.

**Supplementary Figure SN4.2.**
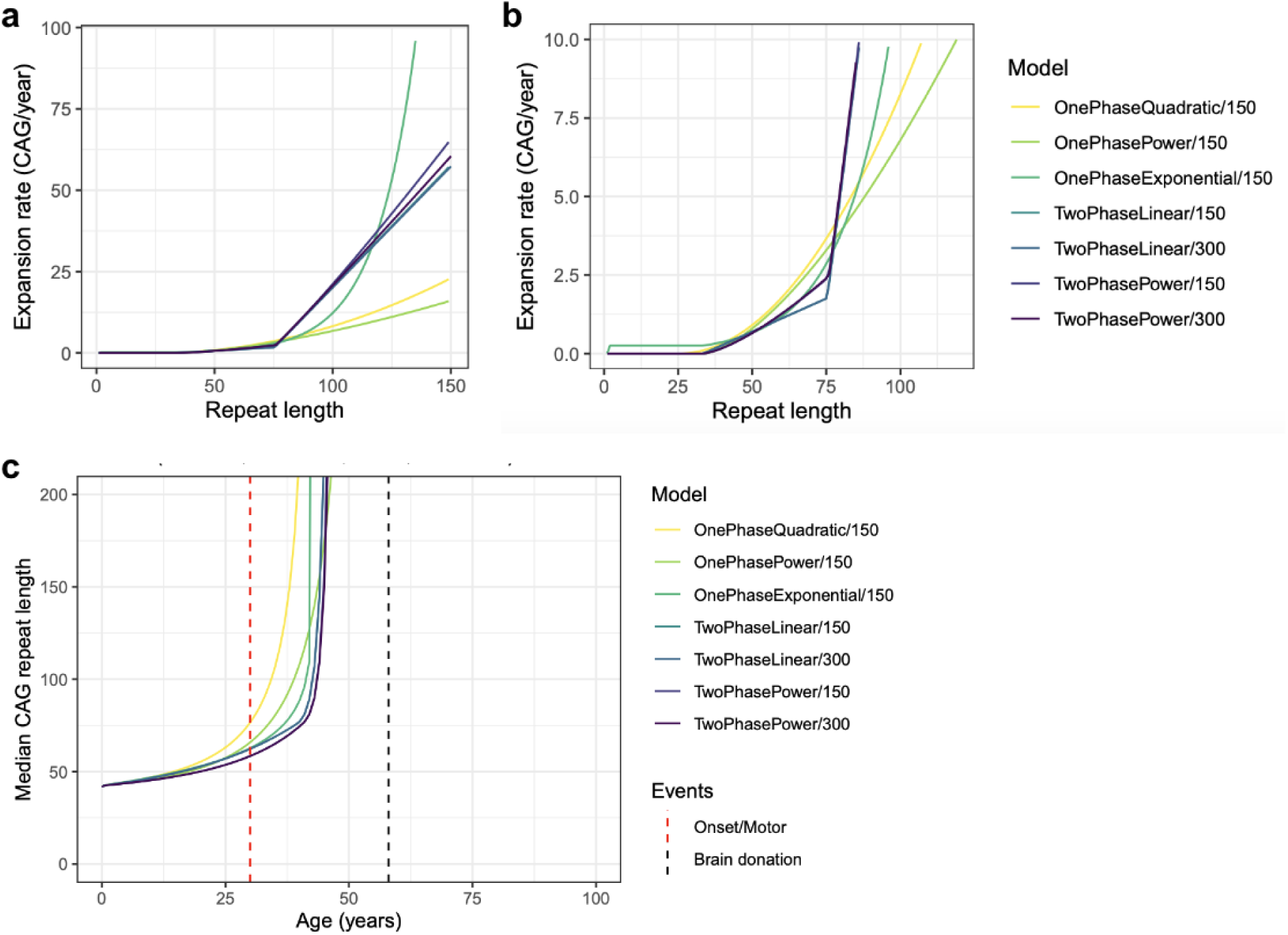
Multiple somatic expansion models converge to the same expansion trajectory. **(a)** Net expansion rate curves inferred from several different models, all fitted to the data for the same representative donor (Donor 5). The two-phase models (darker colors), which provide better fits to the data overall, model similar trajectories. The one-phase models (lighter colors), which have less free parameters, attempt to provide the best possible fits to this same trajectory. **(b)** An enlarged view of the expansion rate curves from (a) highlighting the critical transition region around 70-90 CAGs. **(c)** The fitted models for the same donor converge to the same trajectory (darkest lines), here represented as the median of the repeat length distribution as a function of the donor’s age. The repeat length distribution plotted here is not censored for cell loss and thus represents the distribution that would be observable if none of the donor’s SPNs were lost due to disease.

**Supplementary Figure SN4.3.**
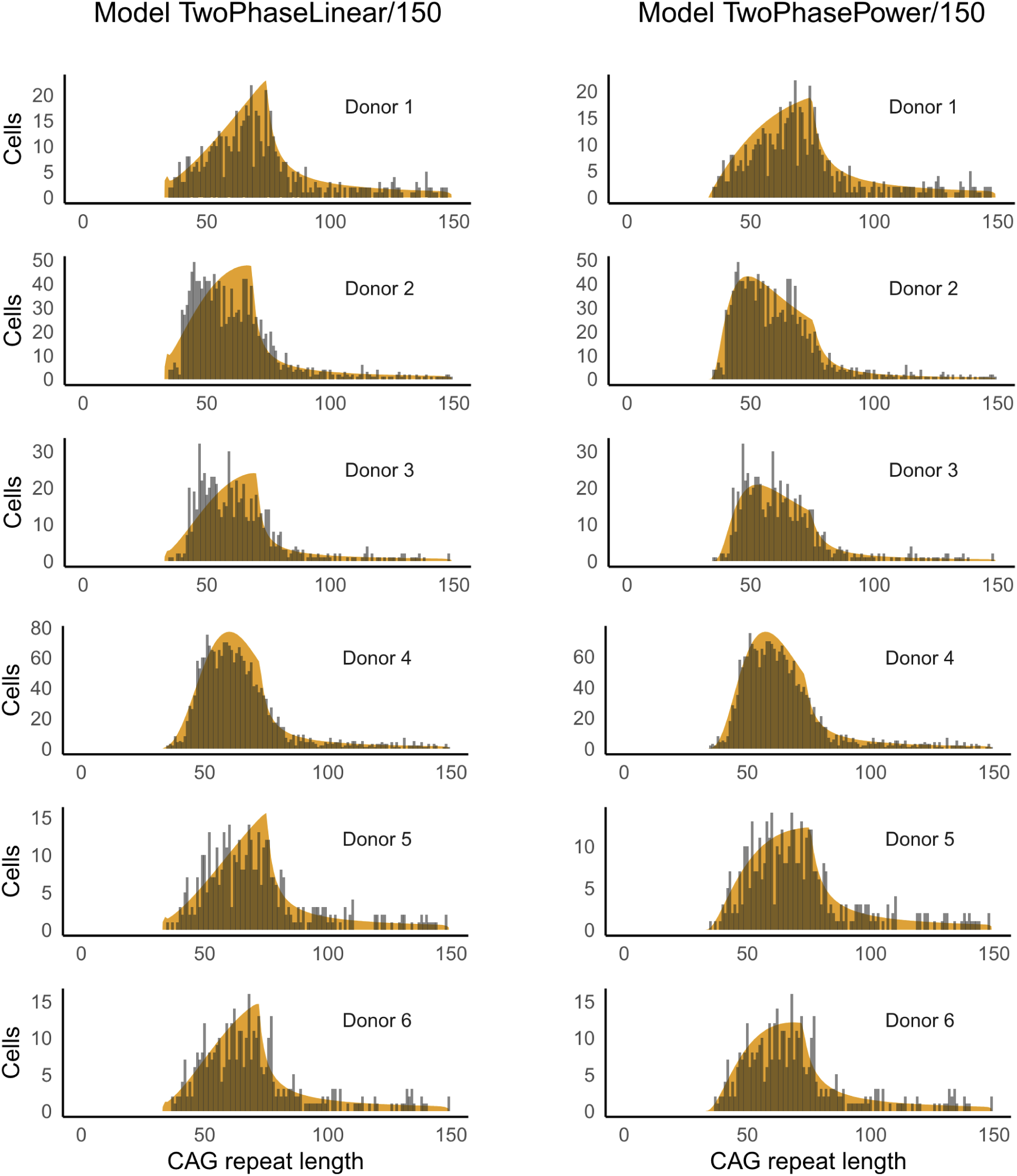
Comparison of two-phase linear and two-phase power models. Plots of the modeled distributions (orange) for caudate SPNs for the six donors with the most data after fitting to the observed distributions of repeat lengths (gray histograms). The two-phase power models (right column) provide better fits to the observed data than the two-phase linear models (left column).

### Characterizing the changes in somatic expansion between phase A and phase B

In our model for HD neuropathology at the single-cell level, we partition the initial stages, prior to the emergence of gene dysregulation into two phases: Phase A corresponds to the slow expansion phase predicted by our two-phase models of somatic expansion and phase B corresponds to the faster expansion phase prior to the beginning of transcriptomic dysregulation around 150 CAGs.

To quantify the transition between and the properties of these two phases, we relied on the two-phase linear model. While the two-phase power model provides a better fit overall and appears to better capture the overall trajectory, the two-phase linear model produces fits to the data that are similar has some advantages for parameter estimation: First, the parameters are easier to compare between the phases, as each phase is a simple linear function representing a kind of average behavior across the phase. Second, because the linear model has fewer parameters, it is easier to interpret and less vulnerable to over-fitting.

The parameters of the two-phase linear model fitted to each donor are shown in **Fig. SN4.4**. The parameters for the rate of expansion (pexp = 0.676 +/- 0.011) and for the threshold to transition between phase A and phase B (threshold2 = 72.2 +/- 2.53) are quite stable across donors. In different models, the value for threshold2 across donors tends to be similar, but the actual value inferred by different models can vary depending on the functional form used for mu(x) in each model and whether a given model allows a greater or lesser degree of smoothing around the transition. As a result, although the models use a discrete value to fit the transition, we recommend that the transition between phase A and phase B should be interpreted as a range. We use a consensus threshold of 80 CAGs to represent the midpoint of this transition range in downstream analyses.

The effective rate of expansion in each phase is rate1 * (2*pexp - 1) for phase A and (rate1 + rate2) * (2*pexp - 1) for phase B. The phase A expansion rate in the six deeply-sequenced donors is 3.51% (+/- 0.83%) CAGs/year and the phase B expansion rate is 57.6% (+/- 12.0%). We used the ratio of these two mean expansion rates (phase B / phase A = 16.4) as our consensus estimate of the change in expansion rates above and below the transition between phases A and B (**Fig. SN4.4**).

**Supplementary Figure SN4.4.**
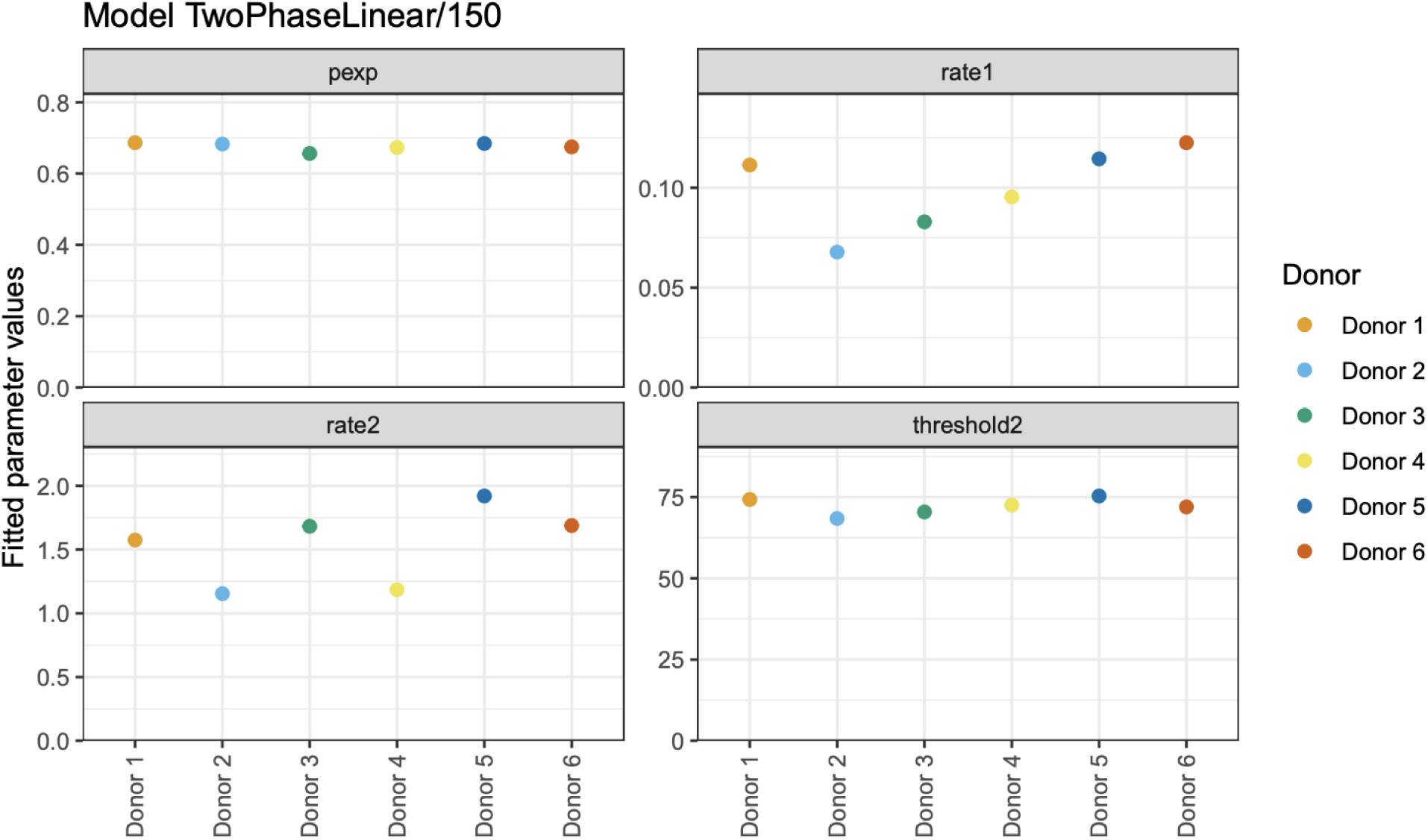
Parameter estimates for two-phase linear model. Best-fit parameter estimates for the four fitted parameters of the two-phase linear model (with cell loss threshold of 150 CAGs). The parameters for pexp (probability of expansion) and threshold2 (T2) the threshold for phase 2, are highly concordant across donors (pexp=0.676 +/- 0.011, threshold2=72.2 +/- 2.53). The effective rate of expansion in phase one (in CAGs/year as a function of the current repeat length) is rate1*(2*pexp - 1) = 3.51% +/- 0.83%. The effective rate of expansion in phase two is (rate1+rate2)*(2*pexp-1) = 57.6% +/- 12%.

### Results have limited sensitivity to the CAG-repeat threshold for SPN death

As described above, we incorporated donor-specific estimates of SPN cell loss in our models of somatic expansion dynamics by assuming a minimum repeat-length threshold for cell loss. Importantly, we avoided assuming any relationship between repeat length and cell loss above this threshold, only that cell loss should be at (or near) zero below this threshold.

To evaluate the sensitivity of our models to mis-estimation of this cell loss threshold, we ran the models assuming a cell loss repeat-length threshold of 150 CAGs and a higher repeat-length threshold of 300 CAGs (**Fig. SN4.5**). The models are relatively insensitive to the exact repeat-length threshold for cell loss. Since the models predict that the rate of somatic expansion is fast when the repeat length exceeds 100 CAGs, changing the assumed minimum repeat-length threshold for cell loss tends to have a minimal effect on the overall trajectory and rate of cell loss.

**Supplementary Figure SN4.5.**
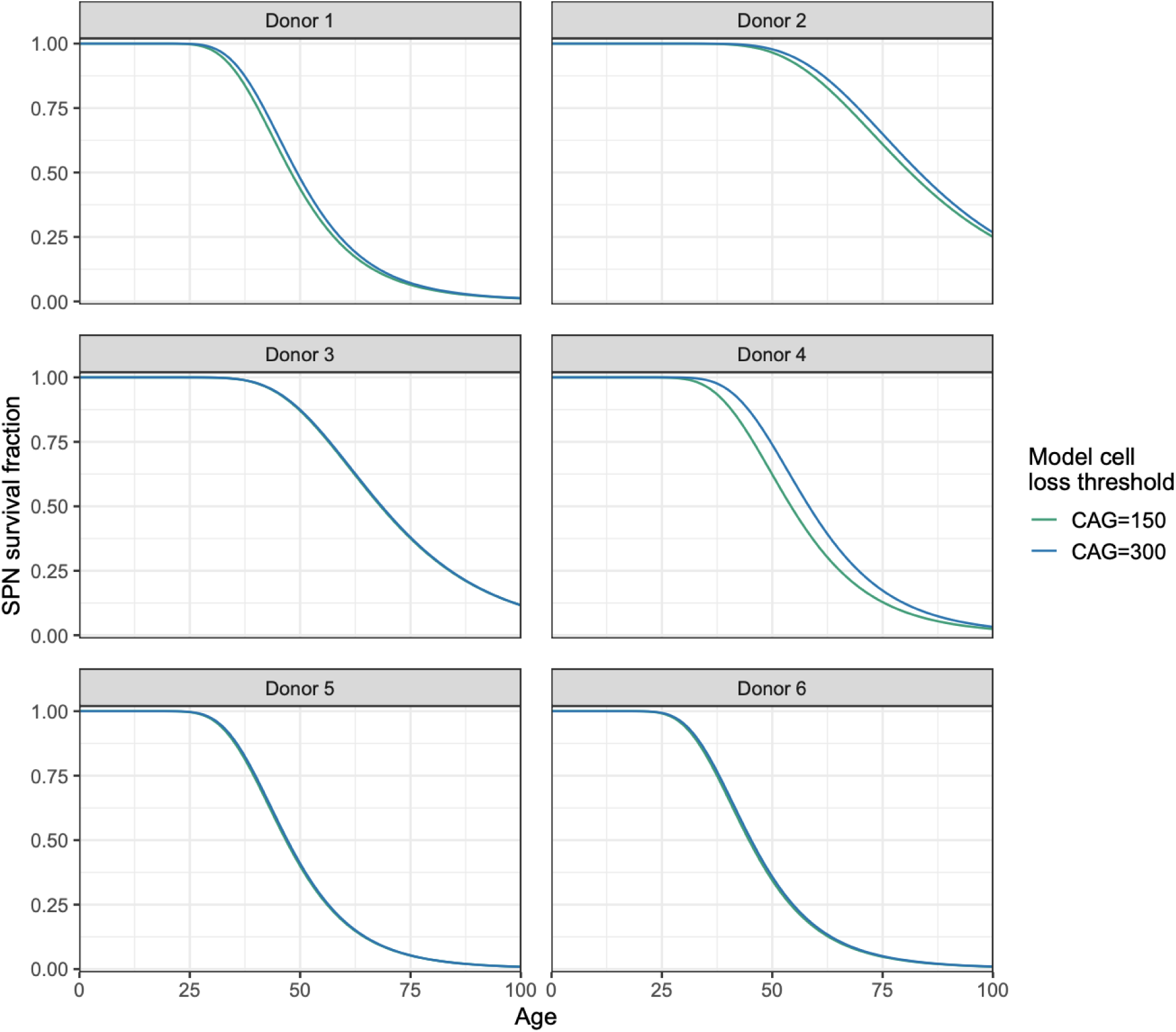
The models have low sensitivity to the minimum repeat-length threshold for cell loss. Plots show two predicted SPN survival curves for each donor, based on a (minimum) threshold of either CAG=150 or CAG=300 for cell death under the two-phase power model. The trajectories predicted under the two models are similar in all donors. This is due to the models predicting high rates of somatic expansion for cells where the repeat has exceeded 150 CAGs.

### Predicting the trajectory of somatic expansion across time

The somatic expansion models fit to the data from each donor allowed us to project the distribution of repeat lengths in each donor both backwards in time and forwards in time from the date of brain donation. **Fig. SN4.6** shows the change in the repeat length distribution over time as a 3D surface model for each donor, based on fitting the TwoPhasePower/150 model.

**Supplementary Figure SN4.6.**
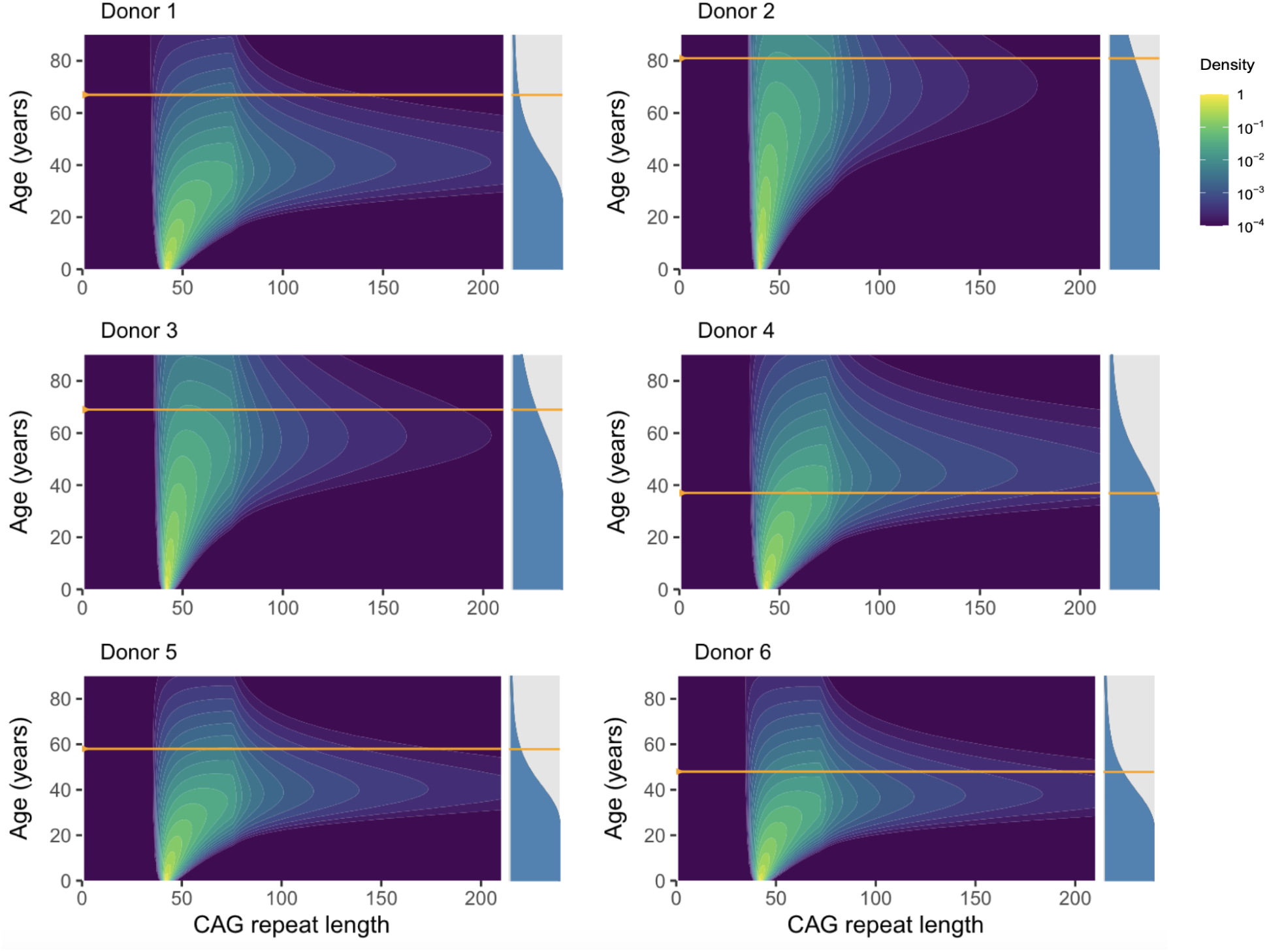
The trajectory of somatic expansion over time in each donor. The models of somatic expansion fit to the data from each donor were used to predict the distributions of the SPN repeat length over time, from birth to time points beyond the date of brain donation (horizontal orange line). The colors on the heat map on the left of each panel indicate the density of SPNs, normalized to the donor’s initial population of SPNs. The bar on the right of each panel indicates the predicted proportion of surviving SPNs (in blue) vs. lost SPNs (in gray) at each age.

**Supplementary Figure SN4.7.**
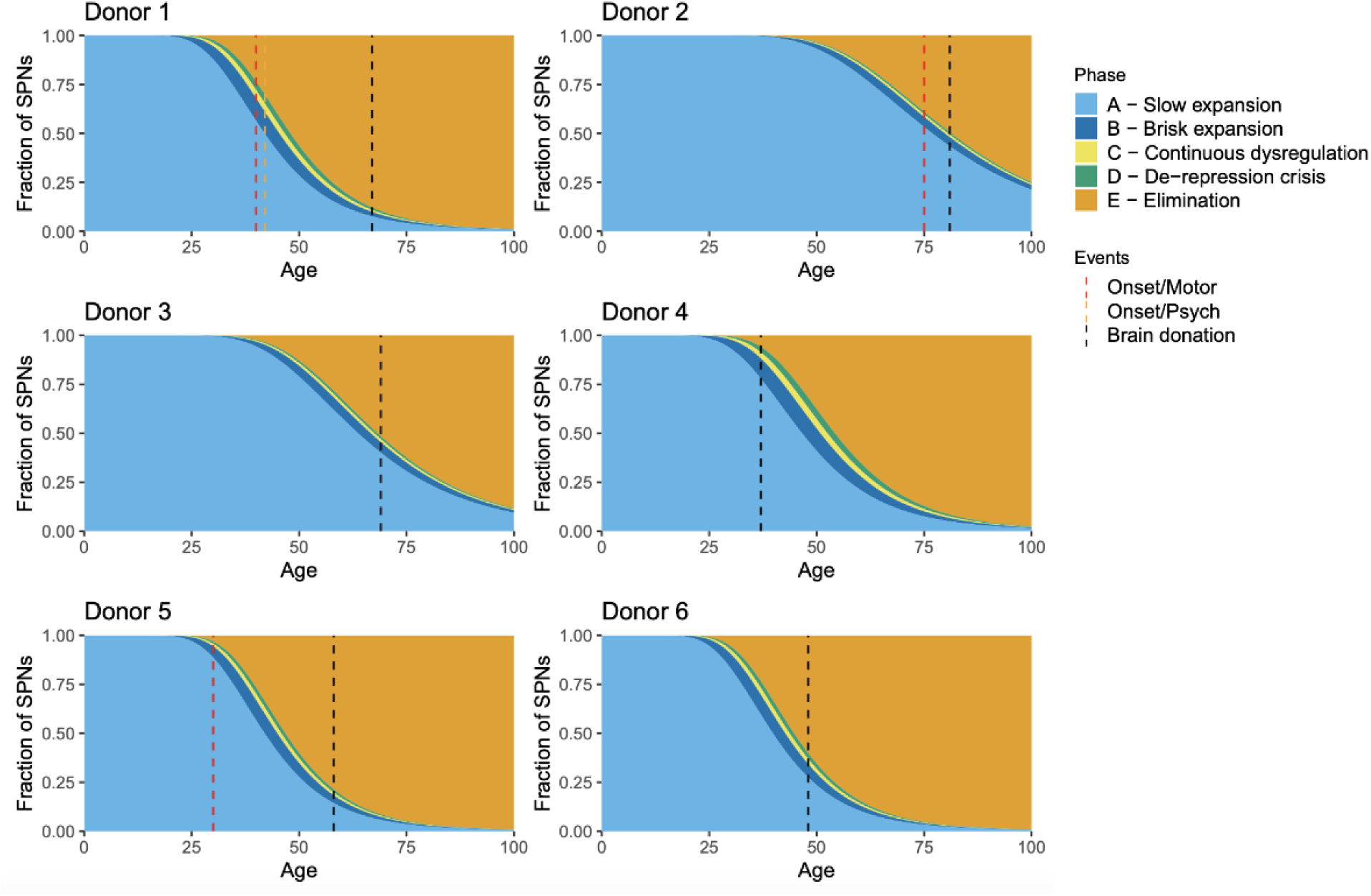
Phases of neuropathology in individual donors. The somatic expansion models fit to the data from each donor were used to predict the distributions of repeat lengths in each donor over time. Discrete thresholds were used to assign a predicted fraction of the donor’s total SPNs (at birth) to one of the five phases of pathology described in Figure 6c. Dashed vertical lines indicate the age at the time of death and brain donation (black) and, where known, the age of symptom onset (red and/or orange).

### Cell loss trajectories for caudate SPNs in individual donors

Our model for neuronal pathology consists of five phases (Figure 7c). Although the repeat-length thresholds at which an individual SPN transitions between these phases are not precise (for example, the transition from slow expansion to fast expansion happens within a range from about 70-90 CAGs), we estimated the fraction of SPNs in each of these phases of pathology over time based on the following repeat-length thresholds: Transition from phase A to B (80 CAGs), transition from phase B to C (150 CAGs), transition from phase C to D (250 CAGs), transition to phase E (500 CAGs) (**Fig. SN4.7**). Because the rate of expansion is rapid when the repeat is highly expanded (> 100 CAGs), these visualizations are not very sensitive to the precise thresholds used; different thresholds would produce qualitatively similar trajectories.

### Simulating the predicted effect of different inherited alleles on age-at-onset

To understand and validate our models for somatic repeat expansion, we experimented with predicting the effects of inheriting different repeat lengths. To do this, we first fitted a model (e.g. TwoPhasePowerModel/150) to the observed data from the individual donors. Then, keeping all other model parameters fixed, we changed only the donor’s inherited repeat length and used the resulting model to predict the distribution of SPN repeat lengths over time. This allowed us to explore hypothetical scenarios relative to the best-fit model for an individual donor.

We further sought to estimate, using our somatic expansion models, a molecular proxy for age-at-onset. In keeping with published reports from pathology studies, at consistent with data from the donors in our study when age-at-onset is available in their medical records, we selected as an age-at-onset proxy the age at which 25% of a donor’s original population of SPNs would be predicted to reach an expanded repeat length of 300 or more CAGs.

We plotted this age-of-onset proxy as a function of different (hypothetical) inherited repeat lengths for each donor (Fig 6e). The resulting curves were similar in each donor and are broadly consistent with the well-documented relationship between inherited repeat length and age-at-onset observed in large cohorts (Genetic Modifiers of Huntington’s Disease (GeM-HD) Consortium., 2019) .

### Estimating somatic expansion prior to future toxicity at age of onset

A question of great interest for patients and for the design of clinical trials is to estimate, at any point in disease, how many of a patient’s SPNs have not yet reached the toxic phase and thus will undergo further somatic expansion to reach this phase. To estimate this from our somatic expansion models, we used the following methodology.

Since we did not have actual age-at-onset data for many of our donors, we first estimated approximate age-at-onset as the predicted age at which 30% of the donor’s SPNs would have reached a repeat length of at least 250 CAGs (using somatic expansion model TwoPhasePowerModel/150). We then estimated, for this age, the fraction of the donor’s SPNs predicted to have a repeat length below 150 CAGs (not yet exhibiting toxicity) compared to the fraction predicted to have a repeat length under 500 CAGs (a conservative threshold for SPNs that would be alive/observable). Across the six donors with sufficient data for modeling, the mean of this quantity was 91.5% (+/- 3.8%).

This estimate was largely insensitive to the threshold we used for estimated age-at-onset, with nearly identical results when using 20% or 40% of SPNs with repeats longer than 150 CAGs. The estimate was also largely insensitive to the age at which the estimate is made. We estimated this quantity (fraction of a donor’s SPNs predicted to have repeat length below 150 CAGs compared to the fraction predicted to have repeat length under 500 CAGs) across all ages (up to 100 years old), covering effectively all disease stages, and computed the minimum for each donor. The mean across donors was 86.5% (+/- 6.2%).

### Estimating mean somatic expansion rates

To estimate the mean number of years for a typical cell to expand from 40 to 60 to 80 CAGs, we used the TwoPhasePowerModel/150 and performed the following analysis: First, we computed adjusted models for each donor as if they had inherited a repeat of length 40. Then we estimated the age at which the median CAG in each donor would reach 60 or 80 CAGs. The mean age of reaching 60 CAGs was 50.7 (+/- 13.5) years. The additional time to reach 80 CAGs was 11.7 (+/- 1.5) years. Reaching 150 CAGs was an additional 3.4 (+/- 0.5) years.

These results highlight that in our models, donors vary more in their predicted overall rate of progression than in their predicted future progression once a milestone has been reached.

For comparison, we also did the same analysis, but modeling each donor as starting with an inherited repeat length of 42 CAGs. In this case, the mean age of reaching 60 CAGs was 37.8 (+/- 9.0) years, the additional time to reach 80 CAGs was 11.6 (+/- 1.5 years) and to reach 150 CAGs was 3.4 (+/- 0.6) years.

### Simulations and animations of the dynamics of somatic expansion

To create animated visualizations of repeat expansion dynamics, we used a fitted repeat expansion model to simulate the dynamics for a specific donor and scenario (e.g. an arbitrary inherited repeat length) simulating the random walk of a large number of cells (typically n=3000). We initialized a vector of repeat lengths to the desired inherited allele length, then at each age from birth to the desired final length we chose the next repeat length based on the transition probability matrix underlying the CTMC model (P = exp(Q)) using a time step of 1 year. To generate the animations, we plotted the results of the simulation at each time step as a png file (using R) and then combined the individual png files as single movie frames into an animated gif using the ffmpeg software.

### Caveats and potential limitations of the somatic repeat expansion models

While the models we have developed are consistent with many features of HD progression, any attempts to generalize from these models to the general population of persons with HD should be done cautiously, for two reasons: First, these models are based on a modest number of donors with the most common HD-causing alleles (40-43 CAGs), and there is substantial inter-donor variability. Second, an important input to each individual’s predicted trajectory is our estimate of the amount of earlier SPN loss that the donor has experienced during their lifetime; we estimate this from the fraction of nuclei sampled that are SPNs (see Fig. 1), but the precision of these estimates is uncertain.

### Software availability

An R library implementing the repeat expansion modeling is available at https://github.com/broadinstitute/HD-CAG-Modeling.

## Methods

### Brain donations, neuropathological diagnosis and ethical compliance

Brain donors were recruited by the Harvard Brain Tissue Resource Center/NIH NeuroBioBank (HBTRC/NBB), in a community-based manner, across the USA. Human brain tissue was obtained from the HBTRC/NBB. The HBTRC procedures for informed consent by the donor’s legal next-of-kin and distribution of de-identified post-mortem tissue samples and demographic and clinical data for research purposes are approved by the Mass General Brigham Institutional Review Board. Post-mortem tissue collection followed the provisions of the United States Uniform Anatomical Gift Act of 2006 described in the California Health and Safety Code section 7150 and other applicable state and federal laws and regulations. Federal regulation 45 CFR 46 and associated guidance indicates that the generation of data from de-identified post-mortem specimens does not constitute human participant research that requires institutional review board review.

The HBTRC/NBB confirmed HD diagnosis and excluded clinical comorbidity and presence of unrelated pathological findings by reviewing medical records and by formal neuropathological assessment. The 1985 Vonsattel et al. grading of neostriatal pathology (Vonsattel *et al*., 1985) was used for diagnosis. Diagnosis on early cases is done using histological stainings and polyglutamine immunohistochemistry (Mattsson *et al*., 1974; Vonsattel *et al*., 1985; Hedreen *et al*., 1991). Positivity in pontine gray neurons rules out HD like-2 neuropathology (Greenstein *et al*., 2007), and cerebellar dentate neurons are mildly positive even in very early cases, while Purkinje cells are negative (unlike in cerebellar ataxia CAG expansion cases).

Affected individuals were selected for this study so as to represent a range of HD stages – from “at-risk” gene-expansion carriers who passed away before symptom onset, to affected persons with advanced caudate neurodegeneration. Experiments utilized fresh frozen brain tissue from each donor.

We sequenced the CAG repeat within the *HTT* gene in each donor’s genomic DNA (isolated from Brodmann Area 17) using the MiSeq assay developed by Darren Monckton’s lab.

### Calculation of CAG-Age-Product score (CAP score)

We used a well-established approach to calculate CAP score (Zhang *et al*., 2011)

*age* * (*inheritedCAGlength* - 33.66)

We also considered a newer formula (CAP-100) (Warner *et al*., 2022) that weights the contribution of age and inherited CAG differently and is standardized such that CAP = 100 at the expected age of diagnosis, though we found that the Zhang formula exhibited a (modestly) stronger relationship to SPN loss across the brain donors in our analyses.

### Extraction and analysis of nuclei in 20-donor “villages” for snRNA-seq

For analyses comparing *across* donors (Fig. 1**-3**, **11**), to make rigorous comparisons of nuclei from many brain donors – while controlling for technical influences from extraction of nuclei, single-cell library construction, and sequencing – we processed sets of 20 brain specimens (each consisting of affected and control donors) at once as a single pooled sample, an approach we have previously described (Wells *et al*., 2023; Ling *et al*., 2024) in which we make preparations of nuclei from sets (or “villages” (Wells *et al*., 2023)) of 20 donors at once. ESpecimens were allocated into batches of 20 specimens per batch. Each set of 20 tissue samples was processed as a single sample through nuclei extraction, encapsulation in droplets, library creation, and sequencing (**Supplementary Fig. 1.1**). We used combinations of hundreds of transcribed SNPs in each cell’s sequence reads to assign each nucleus to its donor-of-origin, using the computational approach we have described previously (Wells *et al*., 2023; Ling *et al*., 2024). This experimental approach allowed the data to be highly comparable donor-to-donor (Fig. 2).

Nuclei were isolated from frozen brain tissue using approaches we have described (Wells *et al*., 2023; Ling *et al*., 2024) and deposited in protocols.io (https://www.protocols.io/view/village-nuclei-isolation-with-optiprep-36wgq3bmxlk5/v1). Briefly, in Ling et al., frozen brain tissues (20 specimens including 10 controls and 10 HD patients were pooled in a village, otherwise each specimen was processed individually for deep-dive experiment) on the glass slide was shaved off, minced and transferred to a 6-well plate containing nuclei extraction buffer {NEB: 1% Triton X-100, 5% Kollidon VA64 in dissociation buffer (DB: 81.67 mM Na2SO4, 30 mM K2SO4, 10 mM glucose, 10 mM HEPES, 5 mM MgCl2 [pH 7.4])}. Tissues were disrupted by pipetting and syringing, and filtered through a 20-micron filter and 5-micron filter serially. The filtered nuclei were resuspended in 50 mL of DB and spun down at 500g in 4C for 10 min. After removing the supernatant, the pellet was resuspended in 1 mL of DB. Nuclei were visualized and counted by staining them with DAPI. For the density gradient-based nuclei isolation, the frozen brain tissue was transferred to dounce homogenizers filled with Nuclei EZ lysis buffer (MilliporeSigma, #NUC101) supplemented with 1 U/uL of NxGen® RNase Inhibitor (Biosearch technologies, #30281). After the tissues were homogenized by douncing, the lysates were filtered with 70 micron cell strainers and spun down at 4C with 500g for 5 min. The supernatant was discarded and the pellets were resuspended in 300 ul of G30 (30% iodixanol, 3.4% sucrose, 20 mM tricine, 25mM KCl, 5 mM MgCl2, [pH 7.8]). The resuspended tissue pellets were layered with 1 mL of G30 and spun down at 4C with 8000g for 10 min. The supernatant was carefully removed and the nuclei pellet was washed twice with 1 mL of wash buffer (1% BSA in PBS supplemented with 1 U/uL NxGen® RNase Inhibitor). The nuclei were resuspended in 50 ul of the wash buffer and counted by using LUNA-FL™ Dual Fluorescence Cell Counter (Logos Biosystems).

### Single-nucleus RNA-seq (transcriptome libraries)

The isolated nuclei were encapsulated into droplets and the snRNA-seq library was prepared by using Chromium Next GEM Single Cell 3’ GEM, Library & Gel Bead Kit v3.1 (10X Genomics, PN-1000121) according to the manufacturer’s protocol with only minor modifications. The libraries were sequenced on Illumina NovaSeq 6000 systems platform.

### Processing snRNA-seq data

Raw sequencing reads were aligned to the hg38 reference genome with the standard Drop-seq (v2.4.1) workflow. Reads were assigned to annotated genes if they mapped to exons or introns of those genes. Ambient / background RNA were removed from digital gene expression (DGE) matrices with CellBender (v0.1.0) remove-background.

All classification models for cell assignments were trained using scPred (v1.9.2). DGE matrices were processed using the following R and python packages: Seurat (v3.2.2), SeuratDisk (v0.0.0.9010), anndata (v0.8.0)(Virshup *et al*., 2021), numpy (v1.17.5), pandas (v1.0.5), and Scanpy (v1.9.1).

### Measurements of CAG-repeat length in individual cells

We also developed a novel approach for sequencing the CAG repeat of *HTT* transcripts in snRNA-seq experiments, and assigning these sequences to the cell from which the HTT transcript was derived. Our approach creates two molecular libraries from each set of nuclei: one library samples genome-wide RNA expression (“transcriptome library”), and another library specifically captures the 5’ region of *HTT* transcripts (“*HTT*-CAG library”) (**Fig. M1**). The presence of cell barcodes, shared between the two libraries, allows each CAG-length measurement to be matched to the gene-expression profile of the cell from which it is derived, and thus to the identity and biological state of that cell.

**Figure M1.**
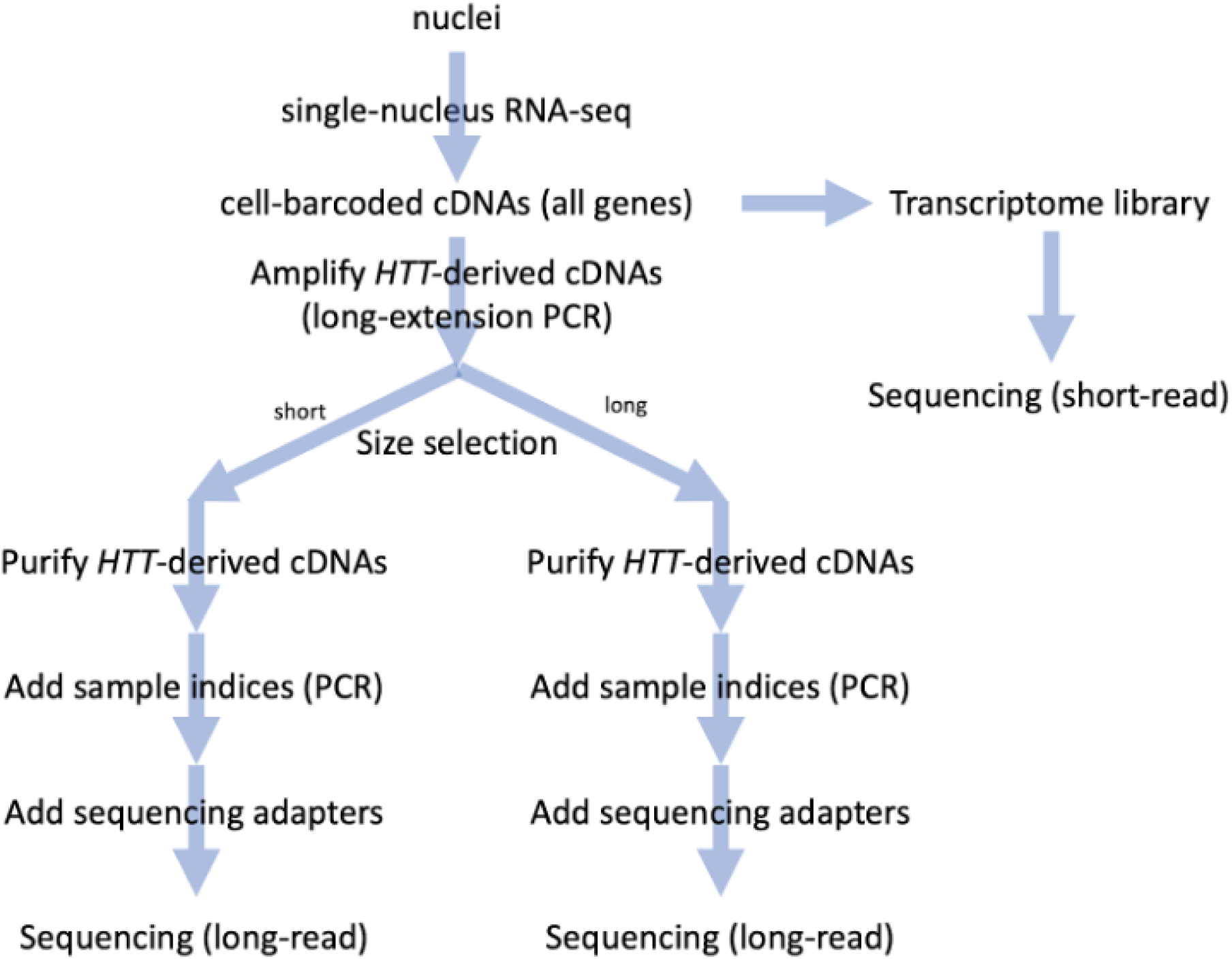
Workflow for creating transcriptome and CAG-repeat libraries from the same nuclei, with a shared set of cell barcodes.

Key aspects in creating these *HTT*-CAG libraries include the use of *HTT*-targeting primers at multiple steps; *HTT*-targeted amplification and purification; steps to preserve long molecules throughout library preparation; careful calibration of PCR conditions to prevent the emergence of chimeric molecules during PCR; and analysis by long-read sequencing. An elaborated, step-by-step protocol, with helpful tips, tricks, modifications and pausing points, is being prepared for release on protcols.io with the publication of this work. We include a summary here.

We begin by isolating and encapsulating nuclei in droplets as described above. To enable the capture and amplification of 5’ HTT sequences, we spike in primers that target *HTT*-specific gene sequences.

We then prepare two libraries from each sample: a standard transcriptome sequencing library, and an *HTT*-CAG library. To generate the *HTT*-CAG library, 4 uL of the cDNA generated from each reaction of the encapsulated nuclei is used for PCR amplification with a biotinylated forward primer that targets *HTT* 5’ of the CAG repeat and a partial read1 primer appended to Nextera adapter sequence. Purification of the resulting PCR product on streptavidin beads (Dynabeads™ MyOne™ Streptavidin C1 (ThermoScientific, #65002)) enriches the library for *HTT* CAG sequences.

We separate the resulting product into long (“L”) and short (“S”) molecular libraries from the same PCR reactions by using SPRIselect beads (Beckman Coulter, #B23319). The next step involves further preparing libraries for long-read sequencing. Each purified “L” and “S” library from the same PCR reactions was indexed with the same Illumina Nextera indices (N701-712 and N501-508), and pooled separately at an equimolar ratio for generating Pacific Biosciences (PacBio) circular-consensus-sequencing libraries. The PacBio libraries were generated by using SMRTbell® express template prep kit 2.0 (Pacific Biosciences, #100-938-900). The “L” and “S” libraries were sequenced on different flow cells on the SEQUEL IIe platform (Pacific Biosciences). We used the adaptive loading feature of the sequencing platform to target a concentration of 100 pM to improve sequencing yield.

### Computational analysis of *HTT*-CAG library data

After sequencing, we processed the reads from each PacBio flowcell using a standard workflow for circular-consensus-sequencing (CCS) base calling and alignment using the following programs: (1) “ccs” (Pacific Biosciences) either version 6.0.0 or version 6.3.0 with standard options (2) “extracthifi” (Pacific Biosciences) to extract only reads with QV >= 20 and (3) “pbmm2” either version 1.4.0 or version 1.10.0 with —preset ISOSEQ and using the GRCh38 “no alt” reference genome (ftp://ftp.ncbi.nlm.nih.gov/genomes/all/GCA/000/001/405/GCA _000001405.15_GRCh38/seqs_for_alignment_pipelines.ucsc_ids/GCA_000001405.15_GRCh3 8_no_alt_analysis_set.fna.gz). Some of the data generated early in this study used one version of the above referenced software while data generated later in the study used updated versions of the same software, but we did not observe any functional differences between the different software versions.

After base calling and alignment, we further processed the data from each flowcell using a custom analysis pipeline we developed (https://github.com/broadinstitute/HTT-CAG-Software). This pipeline consisted of the following steps.

Each read was analyzed (“decoded”) based on the expected layout based on the library construction protocol (**Fig. M2**). The decoding algorithm searched each read for a particular set of landmarks, identifying the landmark using a sensitive Smith-Waterman alignment algorithm designed to accommodate base-level errors and base insertions or deletions in the input reads. Based on the recognition of these landmarks, the read was divided into segments as illustrated.

**Figure M2.**
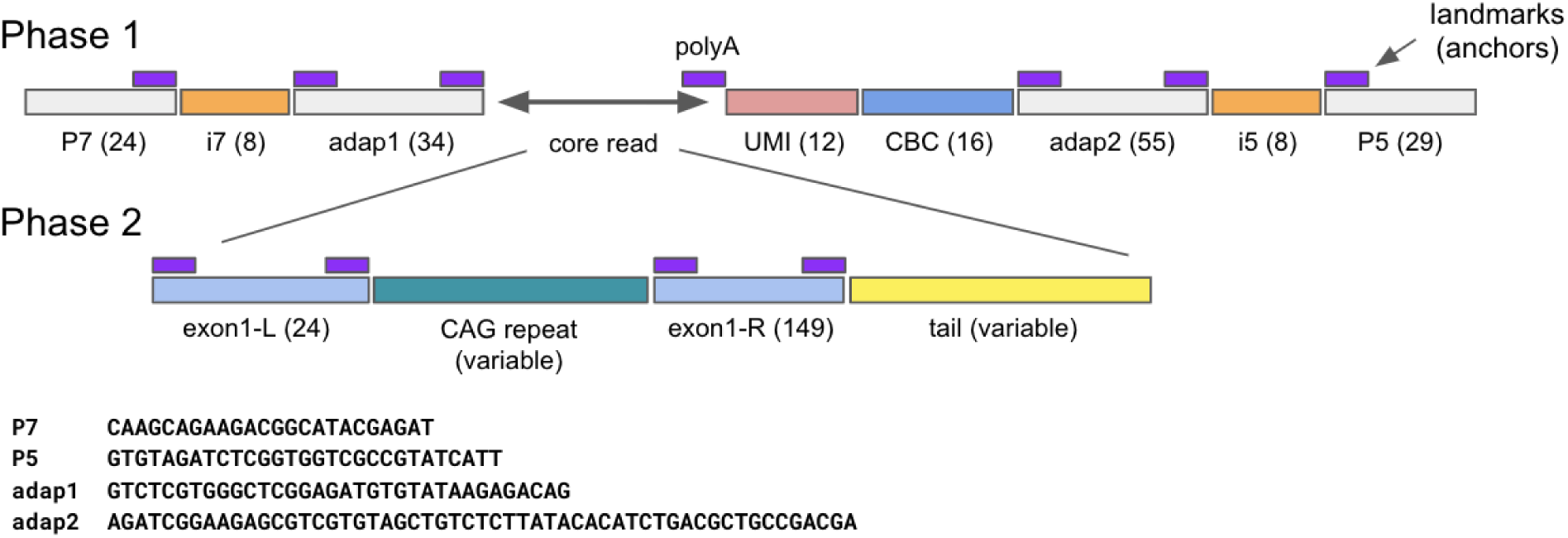
Schematic illustration of long-read decoding in the *HTT* CAG repeat assay. The colored boxes are a schematic representation of the contents of one long read from our *HTT*-CAG library. The computational decoding of each long read was done in two phases. Phase 1 decoded the outer wrapper of the library construct, which included the Illumina sequencing adapters (P7 and P5), Illumina indexes (i7 and i5) which were used to multiplex different snRNA-seq libraries on the same PacBio flowcell, Illumina Nextera and Truseq adapters (adap1 and adap2) and the unique molecular identifier (UMI) and cell barcode (CBC) attached during snRNA-seq library construction. Phase 2 decoded the sequence derived from the host cell’s cDNA, including the initial portion of *HTT* exon 1 (exon1-L), the variable length CAG repeat region, the remaining portion of *HTT* exon 1 (exon1-R) and a variable length tail sequence, which may contain downstream exons of *HTT* or, if the source transcript was unprocessed, sequence from *HTT* intron 1. Numbers in parentheses indicate the expected length (in base pairs) of each read segment. Purple rectangles indicate fixed sequences which were recognized using fuzzy matching to known sequences to identify landmarks that were then used as anchors to divide each read into its constituent components.

We decoded both reads that were aligned to exon1 of the *HTT* gene and unaligned reads. We found it was important to decode both aligned and unaligned reads as the aligner (pbmm2) exhibited bias in the success rate for correctly aligning reads to *HTT* exon 1 based on the CAG repeat length in the decoded read. When decoding aligned reads, we used the strand indicated by the alignment. For unaligned reads, we attempted to decode the read both as recorded in the input file and reverse-complemented and used the most confident decoding in the analysis.

Each read contained two index sequences (i5 and i7) which were used to identify the input snRNA-seq library (“reaction”) when multiple reactions are multiplexed together on one PacBio flowcell. Given a list of input reactions and the pair of indexes used for each, we employed fuzzy matching to find the best match for each index sequence in the decoded read and assigned the matching input reaction if one or both indexes were able to unambiguously identify the input reaction. After determining the input reaction for each read, the cell barcode (CBC) and unique molecular identifier (UMI) were extracted from the decoded reads and reads from the same reaction containing the same CBC and UMI were aggregated. The CAG repeat length was estimated as the consensus length across all reads sharing the same CBC and UMI using the half-sample mode (hsm function in R library modeest). This yielded one repeat length measurement for each CBC+UMI combination.

Several quality control steps were then applied. To remove low quality reads, reads were dropped if the decoded UMI was longer than 28 base pairs or when the CAG repeat “purity” (fraction of bases not matching a pure CAG-triplet motif) was less than 90%. A CBC+UMI combination was retained only if it was supported by 10 or more PacBio reads. Within each reaction, we computed the Levenshtein edit distance (allowing indels) between all pairs of CBC+UMIs (R function “stringdist”, method=”lv”). To avoid double counting, if any CBC+UMI had another CBC+UMI from the same reaction with an edit distance of less than 4 changes, only one measurement was retained, preferring the one supported by the most reads. In principle, we could have performed error correction on the CBC+UMIs, but in practice we found that among “nearby” CBC+UMI values (likely generated by sequencing error) usually only one CBC+UMI was supported by a preponderance of the reads. The reaction from which the *HTT*-CAG library originated also underwent transcriptome analysis and QC. We retained only measurements where the CBC was found to (exactly) match a CBC from the same reaction that had passed all transcriptome QC.

After QC, in the small fraction of cases where we made multiple repeat-length measurements for the same cell (CBC) but with distinct UMIs, we used these for quality assessment (Fig. 2b), but then retained only one measurement, keeping the UMI with the largest repeat measurement. This ensured that if we measured both the short and long alleles from a cell, we used the long allele in downstream analyses.

### Analyses of CAG-repeat effects on SPN gene expression – groups of SPNs defined by CAG repeat-length ranges (Wilcox text)

Analyses of the effect of CAG-repeat length on genome-wide gene expression in subsets of SPNs defined by CAG-repeat length ranges (Fig. 3b**,c**; **Supplementary Fig. 5-6**) utilized a Wilcox rank-sum test (wilcox.test() function in R), in which the individual SPNs (in two groups of SPNs defined by CAG repeat-length ranges) were ranked by their expression level of a gene (as a fraction of all UMIs detected in a cell), and then the distribution of ranks were compared between the two groups. We utilized the Wilcox test for these specific analyses because (i) it makes no assumptions about parametric properties of the expression data, and (ii) the p-value distributions it generates were appropriately null in control permutation analyses in which CAG-repeat lengths were permuted across the individual cells.

### Analyses of CAG-repeat effects on SPN gene expression – Negative Binomial Regression

For various analyses of the effect of CAG-repeat length upon SPN gene expression, Negative Binomial Regression (NBR) analyses allowed the exploration of a wide range of functional forms for models of continuous effects of CAG-repeat length (in contrast to the Wilcox test focus on comparisons of cell groups). Please see **Supplementary Note 3** for a detailed description and discussion of these methods including the variety of alternative methods and models that were evaluated.

### Identification of phase-D genes

Prominent among the genes found to change expression with CAG-repeat expansion beyond 150 CAGs (Fig. 3) were a set of genes that exhibited particularly large fold-changes; inspection of the data revealed that these large fold-changes resulted from these genes’ being almost completely repressed at baseline (in SPNs with <150 CAGs), and that these genes tended to have become de-repressed together in the same SPNs: generally, a specific subset of SPNs with particularly long expansions of the CAG repeat. To identify such genes systematically, we implemented a multi-stage approach, utilizing the single-SPN CAG-length and RNA-expression data from the six deeply sampled donors with manifest HD clinical motor symptoms (donors 1-6 in **Supplementary Table 1**). In stage 1, we identified genes for which transcripts were (i) detected in fewer than 1% of those SPNs with <100 CAGs, and (ii) detected in a significantly larger fraction (p < 10^-5^ by Fisher’s exact test) of those SPNs with > 200 CAGs. This initial screen identified 89 genes (29 of which are in the HOX gene loci, and 60 of which are at other loci dispersed across the genome). Analysis of the expression of these genes in the single-SPN CAG-length and RNA-expression data also revealed that these genes were expressed in only a subset of those SPNs with >200 CAGs. In stage 2, we used the genes identified in stage 1 to refine our definition of the cells of interest, so that these were limited to 174 SPNs with >200 CAGs in which we also detected at least 3 UMIs from these 89 genes collectively (a “test set” of SPNs); we also defined a “control set” of 5,282 SPNs with <150 CAGs in which we detected only 0-1 UMIs from these 89 genes collectively. In stage 2, we identified genes for which transcripts were (i) detected in fewer than 1% of those SPNs in the “control set”, and (ii) detected in a significantly larger fraction (p < 10^-5^ by Fisher’s exact test) of those SPNs in the “test set”. This analysis identified 173 genes, of which 39 are in the HOX loci. These 173 genes served as input to the Negative Binomial Regression analyses of phase-D gene expression in **Supplementary Note 3**.

### Modeling and simulation of CAG-repeat expansion dynamics

Please see **Supplementary Note 4** for details on these analyses and simulations. To create animated visualizations of repeat expansion dynamics, we first fitted a specific repeat expansion model (TwoPhasePowerModel/150, **Supplementary Note 4**) to model the dynamics for a specific donor and potentially a specific scenario (e.g. a hypothetical inherited repeat length) and then simulated the random walks of a large number of cells (n=3000). We initialized a vector of repeat lengths to the desired inherited allele length, then at each age from birth to the desired final length we modified the repeat length based on the transition probability matrix underlying the chosen model using a time step of 1 year. The final animations were created by plotting each individual movie frame separately (using R) as a png file and then combining the individual frames into an animated gif using the ffmpeg software (version 6.6.1).

